# Structural and genetic determinants of zebrafish functional brain networks

**DOI:** 10.1101/2024.12.20.629476

**Authors:** Antoine Légaré, Mado Lemieux, Vincent Boily, Sandrine Poulin, Arthur Légaré, Patrick Desrosiers, Paul De Koninck

**Affiliations:** Centre de recherche CERVO, Québec (Québec), Canada; Département de biochimie, de microbiologie et de bio-informatique, Université Laval, Québec (Québec), Canada; Département de physique, de génie physique et d’optique, Université Laval, Québec (Québec), Canada; Centre interdisciplinaire en modélisation mathématique de l’Université Laval, Canada

## Abstract

Network science has significantly advanced our understanding of brain networks across species, revealing universal connectivity principles. While human studies based on magnetic resonance imaging (MRI) have established several network principles at macroscopic scales, recent breakthroughs, including high-amplitude regional co-activation patterns and spatially contiguous functional gradients, remain unexplored at cellular resolution in animal models. Here, we employ whole-brain functional imaging at cellular resolution in larval zebrafish, combined with anatomical and spatial genetic expression profile databases, to investigate the structural and genetic basis of functional brain networks. We show that mesoscopic functional connectivity (FC) is a robust measure of brain activity that captures the individuality of larvae. Using a public dataset of thousands of single-neuron reconstructions, we reveal a strong coupling between FC and structural connectivity (SC). Numerous properties of the connectome that account for indirect pathways and diffusion mechanisms individually and collectively predict interregional correlations. The hierarchical modular structure of SC and FC significantly overlaps in space, and modules identified within the connectome constrain the shape of both spontaneous and stimulus-driven activity patterns. Using visual stimuli and tail monitoring, we identify a functional network gradient that maps onto the sensorimotor function of brain regions. Finally, we identify a set of genes whose co-expression in brain regions significantly predicts regional FC. Our findings reproduce several key features of mammalian brain networks in zebrafish, demonstrating the potential for studying large-scale network phenomena in smaller, optically accessible vertebrate brains.

## Introduction

Our understanding of the human brain has evolved from studies of diverse animal models, from tiny worms to non-human primates. Despite considerable anatomical differences between the nervous systems of mammals and more primitive animals, most are built from similar cellular architectures ^1^, synaptic structures ^2^, and neurotransmitter systems. These commonalities underlie remarkably conserved connectivity principles at the level of neuronal networks. Through the mathematical description of the brain as a set of nodes connected by edges, network science has been instrumental in characterizing the architecture of brain networks across species ^3–5^. Several organizational features have been consistently reported across animals and spatial scales, and are now considered canonical properties of brain networks. These features include economic wiring principles ^6^, densely interconnected and functionally specialized subnetworks called modules ^7^, efficient information routing through short network paths ^8,9^, and rich clubs of disproportionately interconnected hubs ^10,11^.

The rules through which brains are wired impose strong constraints on their dynamics. Many studies have characterized the structure-function relationship of human brain networks ^12–14^ through the comparison of two measurements: structural connectivity (SC), the strength of anatomical connections between two neurons or brain regions; and functional connectivity (FC), the statistical associations between neural activities — most frequently the correlation coefficient calculated on hemodynamic signals of two brain regions. SC has been shown to constrain both the brain’s minute-timescale activity correlations ^15^, as well as its transient patterns of activation observed at rest or under cognitive demand ^16^. The general consensus, however, is that predicting FC from SC only is insufficient ^13^, prompting new efforts to compile maps of biological features ^17^, such as regional gene expression levels, to enrich network models with node annotations and biological detail ^18^. Mounting evidence indicates that, alongside SC, transcriptomic signatures influence the organization of functional brain networks ^19^. Collectively, this body of work highlights how the brain’s structural wiring and genetic factors act in concert to support a complex functional repertoire. However, much of this research is based on magnetic resonance imaging (MRI), which is confined to the macroscopic scale of the brain where cellular mechanisms are indistinguishable. Although multiple network features derived from MRI studies are now considered universal, recent breakthroughs in network neuroscience have yet to be thoroughly investigated at cellular scales in animal models – such as the emergence of high-amplitude regional co-activation patterns ^20^ shaped by connectome modules ^21,22^, or the organization of brain functions along spatially contiguous gradients of connectivity ^23,24^. The mechanisms of numerous large-scale network phenomena remain obscure in humans due to technical challenges and limitations, and while animal models could provide greater experimental accessibility, it is unclear if these phenomena unfold in much smaller model systems.

Advances in genetically encoded sensors, spatial transcriptomics, and optical microscopy provide a fertile landscape to characterize the interplay between structure, function, and genes in brain networks at much greater spatiotemporal resolutions in small animal model systems ^25–27^. In particular, the larval zebrafish occupies a strategic niche, as a vertebrate model, for optical methods to achieve live whole-brain imaging at cellular resolution ^28,29^. Moreover, recent efforts to build atlases of the larval brain (reviewed in ref. ^30^) have generated open-access data on anatomical structures ^31,32^, spatially resolved gene expression markers ^33^, and neural connectivity ^32^, providing an essential scaffold for whole-brain functional connectivity studies in zebrafish. Prior work has characterized the organizational features of zebrafish FC using whole-brain lightsheet imaging data ^34^ at cellular resolution, noting similarities with larger scale functional networks ^35^. Other studies have employed network analysis to reveal brain-wide functional alterations following mutations in larvae ^36–38^, bringing new methodologies to the extensive genetic toolbox available in zebrafish.

Here, we combined whole-brain two-photon calcium imaging with recent structural connectivity and gene expression datasets ^32,33^ to investigate the structural and genetic determinants of functional brain networks in zebrafish. We first demonstrated the methodological robustness of FC across three whole-brain imaging datasets, as well as its ability to reliably distinguish individual larvae. Then, we observed strongly coupled functional and structural brain networks, with large amplitude co-fluctuations in brain activity that spatially overlapped with modules identified within the connectome. In addition, we used visual stimuli and behavioral monitoring to uncover a main functional gradient coinciding with the sensorimotor function of brain regions. Finally, we identified regional gene expression barcodes that could predict FC, independent of SC. Our results in zebrafish mirror several recent neuroimaging observations in humans and mice ^23,24,21,22^, highlighting the potential of intersecting multiple optical microscopy datasets to investigate large-scale network phenomena, typically studied in large mammalian brains, in the much smaller and optically accessible vertebrate brain of the larval zebrafish.

## Results

### Two-photon imaging of brain-wide functional networks in zebrafish larvae

We used resonant-scanning two-photon microscopy to record brain-wide neuronal activity at single-cell resolution in transgenic zebrafish larvae expressing a nuclear-localized pan-neuronal Ca^2+^ sensor ^39^ (Tg*(elavl3* :H2B-GCaMP6s), *n* = 22 larvae, 5-7 days post fertilization (dpf)). Larvae were head-restrained in agarose under the microscope, while their tail movements were monitored using an infrared camera (Fig. 1**a**, Methods). Concurrently, larvae were exposed to static whole-field illumination for 10 minutes to record spontaneous brain activity, followed by short successions of abrupt dark transitions (dark flash stimuli ^40^) over a 3-minute period (Methods). We imaged 21 non-overlapping planes at 1 Hz (Supp. Video 1, separate imaging planes; Supp. Video 2, 3D projection with behavior), spanning the entire larval brain with the exception of the most ventral neuronal populations (Fig. 1**b**, Supp. Fig. S1**a**). After correcting lateral motion and extracting nuclear Ca^2+^ signals ^41–43^, we used image registration ^44^ to align the calcium imaging volumes to a brain atlas (mapZebrain ^32^, Methods). We then mapped all imaged cells into 70 anatomical brain regions spanning both hemispheres (Fig. 1**c**, Supp. Fig. S1**b**), excluding a few regions that were poorly sampled across animals (Supp. Table 1, Methods). In total, we recorded activity from 54, 464 ± 5086 neurons distributed in 65 brain regions, representing approximately half of the total neuronal population at this stage of development ^28^.

**Table 1:**
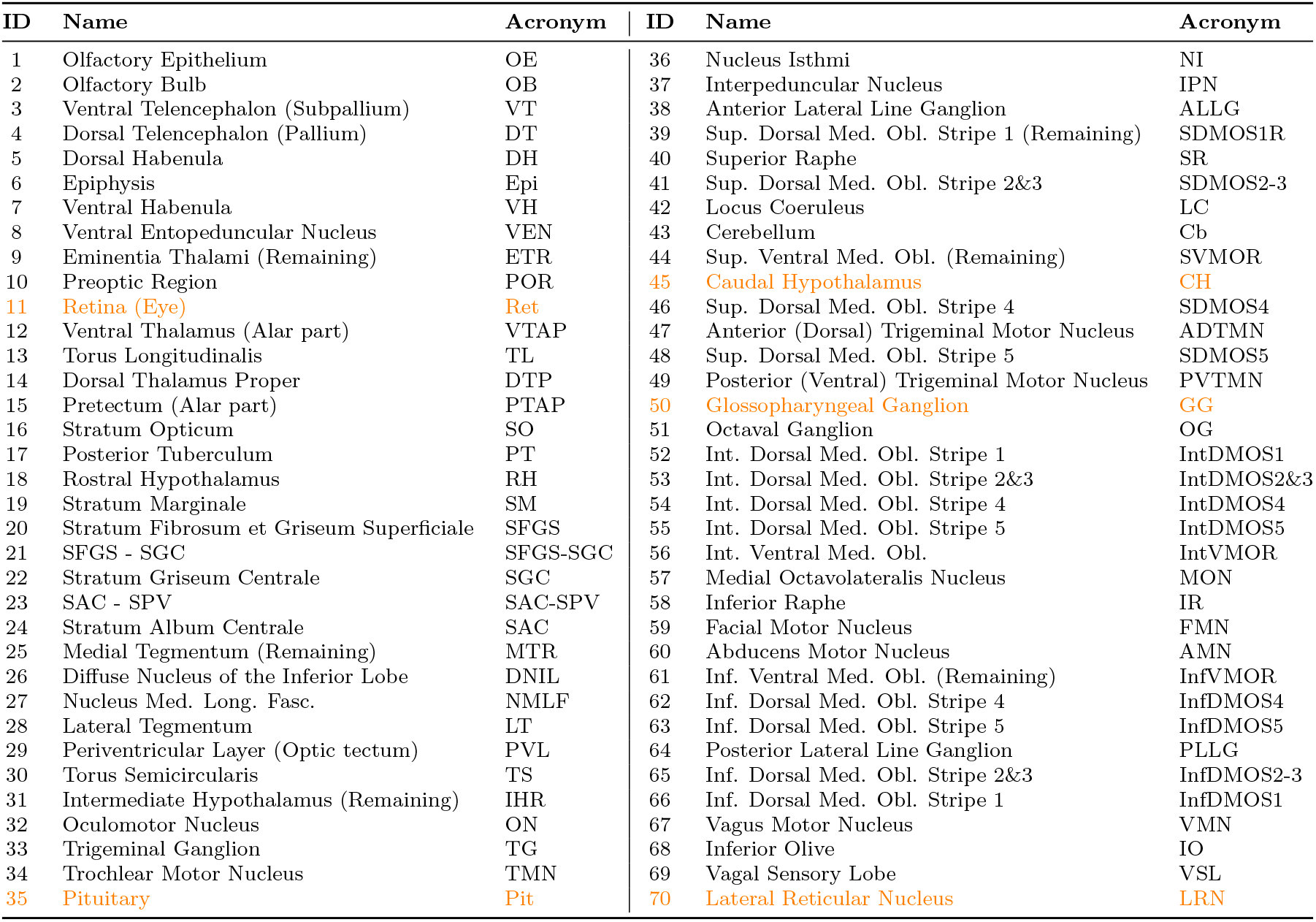
List of 70 mutually exclusive and collectively exhaustive brain regions from mapZebrain used throughout this study. Regions that were excluded from FC analysis are indicated in orange.

**Figure 1:**
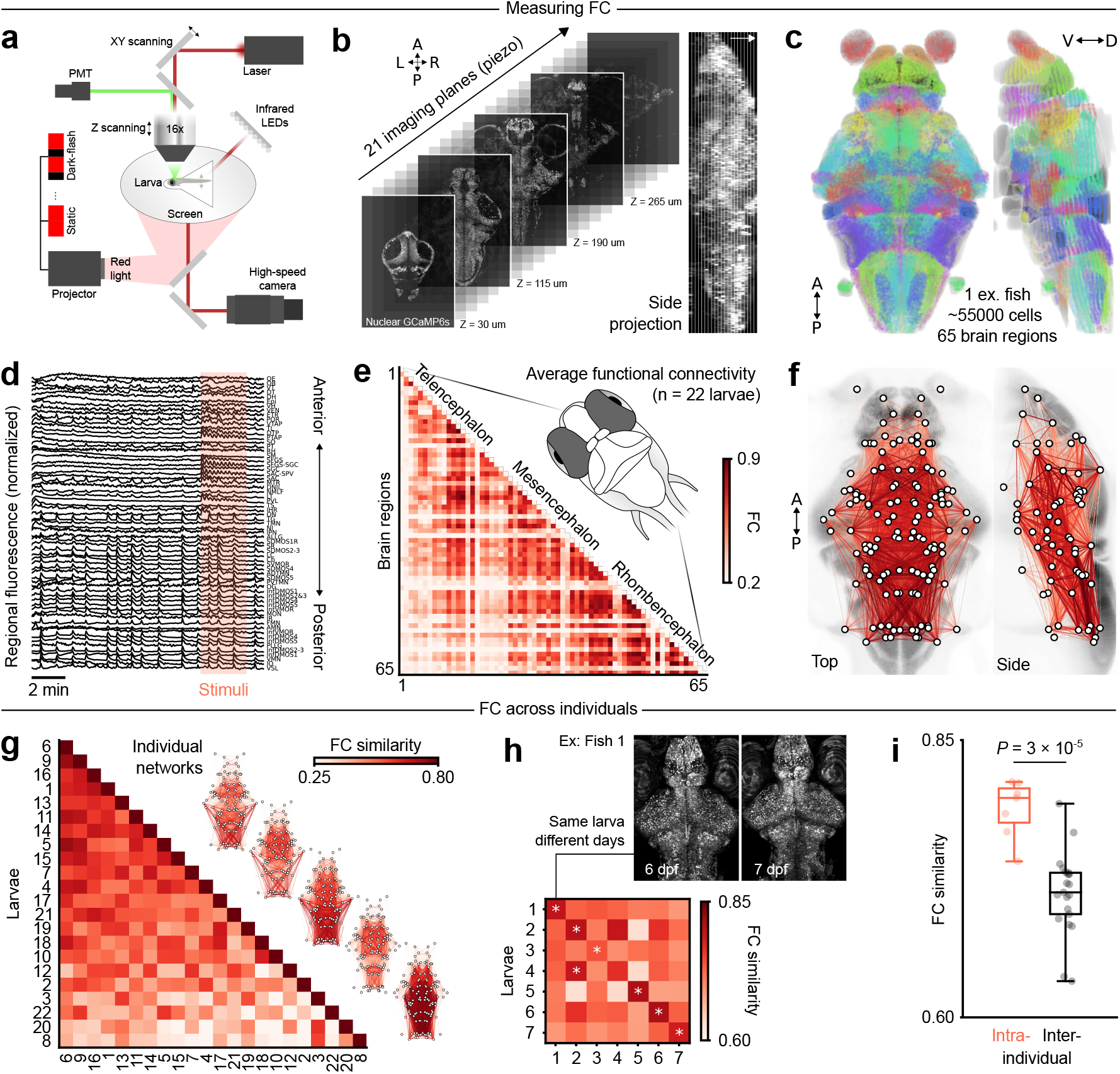
Brain-wide imaging of functional networks in zebrafish larvae. (**a**) Experimental configuration for whole-brain two-photon imaging, light stimulus delivery and behavioral monitoring in Tg(*elavl3* :H2B-GCaMP6s) zebrafish larvae. (**b**) 5 planes selected from 21 functional imaging planes at different depths (left); side projection in one example fish (maximum intensity, right); white arrow indicates dorsal side; refer to Supp. Fig. S1 for image scales. (**c**) Centroids of automatically identified neurons from one larva, registered to mapZebrain and mapped into 70 brain regions (pseudocolors, retina/eye is not displayed). (**d**) Regional fluorescence time series from one representative larva, ordered from anterior to posterior regions; a pink rectangle highlights a period of dark-flash visual stimuli. (**e**) Group-averaged FC matrix (*n* = 22 larvae), ordered from anterior to posterior regions. (**f**) Network visualization of group-averaged FC, using the same color map as the previous panel; nodes represent brain region centroids, and bottom quartile FC edges are not displayed; network edges are mirrored across both hemispheres for visualization. (**g**) Functional network similarity across larvae, ordered from the most globally similar to the most globally dissimilar individual; five example individual networks are plotted on the right. (**h**) Functional network similarity of 7 larvae imaged on consecutive days; similarity scores are averaged across both temporal directions, from dataset 1 to dataset 2 and vice-versa; white asterisks denote maximal similarity values on each row. (**i**) Network similarity scores are significantly higher when comparing individuals to themselves (diagonal values from matrix in **h**) rather than different individuals (off-diagonal values from matrix in **h**) (*P* = 3 × 10^−5^, *t*-test). A, anterior; P, posterior; V, ventral; D, dorsal; L, left; R, right.

To quantify the FC of the larval brain, we averaged the activity of neurons within each region of the brain during the spontaneous period (Fig 1d), then calculated the pairwise correlations between regional signals (Methods). Averaging FC matrices between individuals yielded a group estimate of functional interactions throughout larval zebrafish brain regions (Fig 1**e**). Hindbrain regions exhibited the highest correlations (Fig 1**f**, right), driven by large-amplitude synchronous calcium transients during motor activity (1**d**, bottom traces; viewable in Supp. video 2). Consistent with previous reports ^35^, correlations decayed with increasing distance between regions (Pearson’s *r* = − 0.489, Supp. Fig. S2**b**), and regressing out a global signal before computing FC revealed multiple negative correlations from the residual signals, primarily between the most spatially distant regions (Supp. Fig. S2**c**-**d**). Throughout this study, we utilized spontaneous group-averaged FC derived from two-photon imaging experiments as a succinct network-based description of brain activity. Visual stimuli and motor recordings are introduced in later analyses to differentiate visual and motor populations across the brain.

### Whole-brain functional connectivity is robust across imaging datasets and reflects individuality

A few studies on zebrafish have utilized network science approaches ^36,35,45,37,38,46^, however, the FC of the larval brain remains sparsely characterized. We started by assessing the robustness of whole-brain FC in larval zebrafish over a variety of experimental parameters. First, we compared our group-averaged FC matrix from Fig 1**e** with one generated from an open-access lightsheet microscopy dataset ^34^ registered to the same brain atlas (Methods). The lightsheet microscopy experiments were conducted in paralyzed larvae expressing a pan-neuronal, fast calcium sensor located to the nucleus, with approximately twice the volumetric sampling rate. Additionally, we compared our current dataset to an older two-photon microscopy dataset generated in our lab, which employed slightly different imaging parameters, including field of view, number of imaging planes, and image resolution (Methods). We obtained similar FC matrices across different microscopy techniques and calcium reporter dynamics (Supp. Fig. S3, Pearson’s *r* = 0.626 between two-photon and lightsheet datasets), and substantial reproducibility for data generated using the same microscopy technique and calcium indicator (Supp. Fig. S3, *r* = 0.901 between two-photon datasets). These experiments, conducted over several years and at different institutions ^34^, underscore the robustness of whole-brain FC in zebrafish larvae.

Zebrafish larvae undergo a strict neurodevelopmental course and exhibit many hardwired behaviors ^47,48^. Thus, we expected that FC across individuals at a similar developmental stage would be stereotyped. The functional networks measured in different larvae were significantly correlated, consistent with previous reports ^35^ (Fig 1**g**, average Pearson’s *r* = 0.491, spatial and temporal permutation tests, *P <* 10^−16^, Supp. Fig. S4, Methods). Although individuals share a common FC scaffold, we examined the individuality of FC by conducting two separate imaging sessions in the same group of larvae 24 hours apart (from 6 to 7 dpf, *n* = 7 larvae). We performed pairwise comparisons of individual FC matrices in both sessions and found that FC matrices from the same subject were most similar to each other compared to other larvae in 6*/*7 cases (Fig 1h). We validated that this high identification rate was statistically unlikely under all possible identity permutations (5040 permutations, *P* = 3 × 10^−4^). Additionally, FC similarity scores at the group level were significantly higher within, rather than between subjects (Fig. 1i, *P* = 3 × 10^−5^, t-test). Interestingly, the similarities between fish were lower after a global signal regression, yet the identification rate increased to 7*/*7 (Supp. Fig. S5**a**, Methods). It should be noted however that larvae could also be recognized based on their average fluorescence levels in both functional and anatomical brain volumes, which likely reflects differences in fluorescent protein expression or morphology (Supp. Fig. S5**a**). Nevertheless, FC-based measures provided a significantly greater contrast between intra- and inter-similarity scores than static fluorescence measures, suggesting that brain activity better separates individuals (Supp. Fig. S5**c**, *P <* 0.05 for raw FC, *P <* 0.001 for FC with global signal regression, one-way ANOVA followed by Tukey’s post hoc comparison).

Together, these results support the high reproducibility of larval zebrafish mesoscopic FC across experimental parameters, as well as its ability to discriminate different subjects, a hallmark of functional brain networks first demonstrated in humans ^49^.

### Structural network properties predict functional connectivity

The structure-function relationship of brain networks ^12,13^ — how physical connectivity predicts correlated activity and vice-versa — has been widely studied in multiple species using a variety of functional and structural measurements ^50,24,51–54,14^. Despite significant work to uncover organizational principles in functional networks of larval zebrafish ^35^, structure-function coupling has not been characterized in the zebrafish brain. To fill this gap, we leveraged a recently established morphological catalog of 4327 individual neurons collected from 6 dpf larvae and registered to the mapZebrain atlas to generate a regional wiring diagram ^32^ (Fig. 2a). We first compared four different methods to compile both directed and undirected SC from individual cell reconstructions (Supp. Fig. S7, Methods), opting for neurite terminal counts (putative synapses) as units of weighted connectivity. We expanded regional boundaries by 30 microns while compiling SC (Supp. Fig. S7**d**) to encompass numerous terminals situated just outside the original regional delimitations (Supp. Fig. S6) and to better account for the spatial extent of putative postsynaptic dendritic arbors at regional boundaries (Methods). Using this method, we found that regional SC (Fig. 2**b**) and FC matrices were highly correlated (Pearson’s *r* = 0.568, Fig. 2**c**; Supp. Fig. S7**i**-**l** for comparison between wiring methods). Note that while we used this definition of SC throughout the article, key results were also replicated with a sparser, more conservative SC matrix (Supp. Fig. S16). We evaluated the significance of the SC-FC correlation using a spatially constrained configuration model (SCCM) of SC, which randomizes network connections while preserving some network features, such as node degrees and the approximate distribution of connection lengths (Methods; Supp. Fig. S8; Supp. Fig. S9 for null distributions, *P <* 10^−16^). We consequently observed that structural and functional weighted degrees were highly correlated (Pearson’s *r* = 0.689, Fig. 2**e-f**), with particularly small dispersion at higher degrees, suggesting that FC and SC networks are characterized by similar connectivity hubs ^55^.

**Figure 2:**
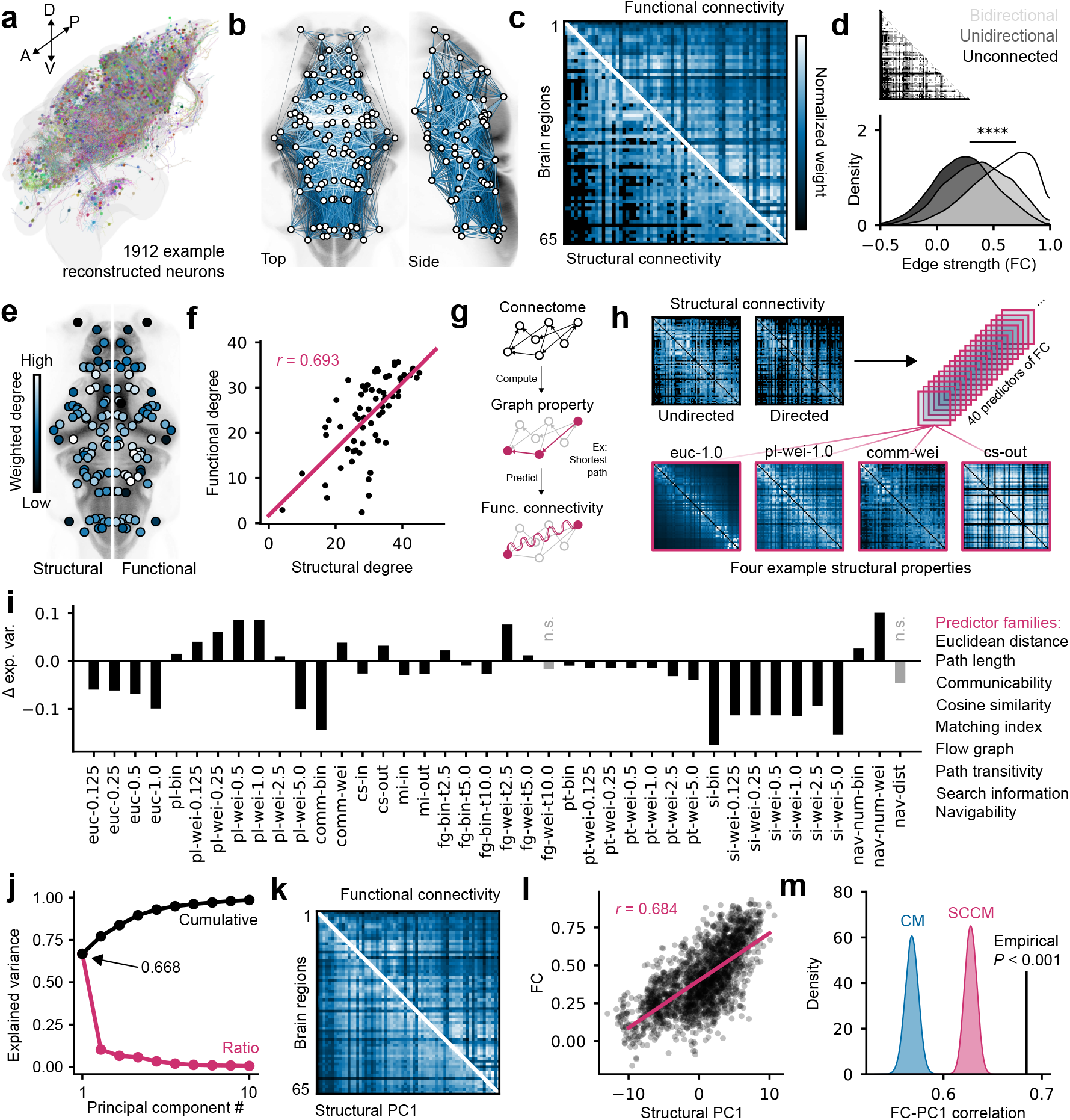
Structure-function coupling of zebrafish brain networks. (**a**) 3D rendering of 1912 reconstructed neurons generated on mapZebrain.com. (**b**) Mesoscopic structural connectivity overlayed on brain anatomy; edge weights are color mapped with the next panel, and bottom quartile edges are not displayed. (**c**) Undirected SC matrix compared with FC; matrices are z-scored and weight values are arbitrary for visualizaton. (**d**) Distributions of individually sampled FC values from edges that have either bidirectional, unidirectional or no underlying structural connections (one-way ANOVA, *F* = 5048, *P* = 0) (**e**) Functional and structural node degrees with arbitrary rescaled values for visualization; structural degrees are displayed on the left hemisphere, and functional degrees are displayed on the right. (**f**) Linear regression between structural and functional degrees (Pearson’s *r* = 0.693). (**g**) Depiction of the FC modeling approach used in following panels; properties of the structural network, such as the shortest path between two nodes, are used to predict functional connectivity. (**h**) Four example matrices from an array of 40 graph properties used as predictors of FC in this study. (**i**) FC variance explained by each of the predictors, subtracted by the variance explained by SC; predictor families are indicated on the right, in the same order as the bar chart labels. (**j**) Relative (red) and cumulative (black) variance explained by each principal component derived from the set of predictors. (**k**) Structural PC1 compared to FC; matrices are arbitrarily scaled for visualization. (**l**) Linear regression between upper triangle values of structural PC1 and FC (Pearson’s *r* = 0.684). (**m**) Null distributions of structural PC1 correlations derived from null SC matrices (1000 matrices per distribution, empirical *P <* 0.001).

Next, we asked whether the presence of bidirectional structural connections between regions predicted more correlated activities. Fig. 2-**d** shows significant shifts in FC edge weight distributions for bidirectionally connected, unidirectionally connected, or structurally unconnected regions, with bidirectional connections exhibiting the highest correlations (one-way ANOVA, *F* = 5048, *P* = 0). This characteristic was preserved to a lesser extent in the SCCM and in a standard configuration model (CM), but vanished when randomly shuffling the SC matrix (Supp. Fig. S10**e**-**h**). This suggests that the correlation between SC and FC degrees — a feature that is approximately preserved in both SCCM and CM — might partially contribute to these distribution shifts, as high degree regions are more likely to be both reciprocally connected and functionally correlated. These analyses highlight that the presence of structural connections is generally associated with greater functional connectivity, with bidirectional connections driving the most correlated activity.

However, FC may depend on mechanisms other than direct anatomical connections, such as polysynaptic interactions ^50^, extrasynaptic signaling ^53^, or global motor activity ^56,57^. To assess the influence of various mechanisms on shaping FC, we computed 40 structural properties of SC ^58^ (predictors) based on binary or weighted, directed or undirected SC matrices, using multiple parameterizations where applicable (Methods). We then evaluated how each property, encapsulated in a 2D matrix of inter-regional scores (Fig. 2**h**), contributes to the variance of FC relative to the SC matrix itself (Fig. 2**i**). These structural properties were derived from 9 families – one geometric and eight topological – each computed for each pair of nodes, displayed from left to right in Fig. 2**i**: Euclidean distance, shortest path length, communicability ^59,60^, cosine similarity, matching index ^61^, flow graphs ^62^, path transitivity ^50^, search information ^63^, and navigability ^64^ (see Methods for complete descriptions). Some properties encapsulate a direct routing of activity through polysynaptic regional pathways (shortest paths), while others describe the random diffusion of activity on the network (communicability or flow graph). Most structural properties were significantly correlated with FC, with only two failing to exceed the correlations of the same properties derived from null SCCM and CM matrix ensembles (Supp. Fig. S11). While many properties were poor predictors of FC relative to the variance explained by the SC matrix, some provided additional explanatory power. For example, the Euclidean distance between regions was a relatively poor predictor of FC (Fig. 2**i**, `euc`), whereas weighted shortest paths explained an additional 8.45% of the variance (`pl-wei-1.0`). Weighted communicability (`comm-wei`) explained 3.76% additional variance, and flow graph at diffusion time *t* = 2.5 (`fg-wei-t2.5`) explained 7.51% additional variance. The best structural predictor of FC was weighted navigation (`nav-num-wei`), with 9.93% additional explained variance. Overall, these results show that multiple SC properties capturing indirect pathways and diffusive communication mechanisms better predict FC compared to direct connections alone.

Building on this observation, we asked whether these predictors collectively contribute to FC. We first noted that all predictors were positively correlated with each other (*r* = 0.653 ± 0.162, mean ± standard deviation of interpredictor correlations), and that a single principal component (PC) explained 66.8% of the variance in the data between all node pairs when considering these predictors as variables (Fig. 2**j**). This first PC, representing a linear combination of all predictors with specific weights (loadings), had scores that were highly correlated with the FC matrix values across all pairs of nodes (Pearson’s *r* = 0.684, Fig. 2**k**,**l**), significantly more than null PC1s derived from SCCM and CM matrix properties (*P <* 0.001, Fig. 2**m**). Note that this consensus predictor obtained using PCA results from the natural variance among predictors and is not explicitly optimized to match the FC matrix. These results indicate that FC is captured to a large extent by a linear combination of structural properties, highlighting the significant role of their joint contribution in shaping FC.

Collectively, these analyses demonstrate a strong coupling between structural and functional networks in larval zebrafish, where direct structural connections account for a significant portion of the observed functional correlations. FC is better predicted by specific properties derived from SC, notably shortest paths, flow graphs, and navigability. These properties combine linearly to collectively explain a large fraction of FC variance, suggesting a cumulative effect of different signaling mechanisms on the underlying structural network.

### Structural and functional modules overlap at multiple hierarchical levels

Brain networks in numerous species are organized into modules (also referred to as communities ^65^) where groups of brain regions are preferentially connected to each other, a feature that promotes functional specialization, resilience, and improved computational capabilities ^7^. A modular hierarchy of FC in zebrafish has been described in a previous study ^35^, but the structural connectivity underlying these modules is unknown. Given the strong coupling between FC and SC, we expected functional modules to emerge perhaps from homologous structural modules. To detect structural and functional modules at different hierarchical levels, we used a consensus ^66^ modularity maximization ^67^ algorithm (Methods). Briefly, we generated consensus matrices that represent the probability that two nodes are assigned to the same module in thousands of partitions with different levels of network subdivision (Fig. 3**a**,**e**). Then, we hierarchically clustered these matrices to generate multilevel node partitions with varying numbers of modules (4 modules are displayed in Fig. 4 **b**,**c** for SC and **d**,**f** for FC). This analysis revealed a maximum of 10 structural modules and 7 functional modules that roughly separated the major anatomical subdivisions of the larval brain (Supp. Fig. S12, Methods). We verified that empirical networks were significantly more modular across all hierarchical levels than multiple randomized null models of SC and FC (Fig. 3**g**, curves represent 99th percentiles of null models, thus *P <* 0.01; Methods). Then, we quantified the overlap between structural and functional partitions using the adjusted Rand index. SC and FC modules overlapped significantly more throughout their hierarchy than they did with partitions generated from null network models (Fig. 3**h**-**i**, *P <* 0.01), except for SCCM which overlapped slightly more with FC at the coarsest hierarchical level where two modules separate the anterior and posterior brain. We also noted that SC and FC modules were spatially compact, significantly more than those of null models with the exception of SCCM, which preserves connection distances by definition (Fig. 3**j**, *P <* 0.001, one-way ANOVA followed by Tukey’s post hoc test for individual comparisons). Interestingly, the FC modules were slightly more spatially dispersed compared to the SC modules, which could reflect the spread of regional activity beyond the structural modules through indirect pathways. Together, these results demonstrate that both SC and FC the in larval zebrafish brain are modular, with overlapping, spatially compact modules subdividing the brain across multiple hierarchical levels.

**Figure 3:**
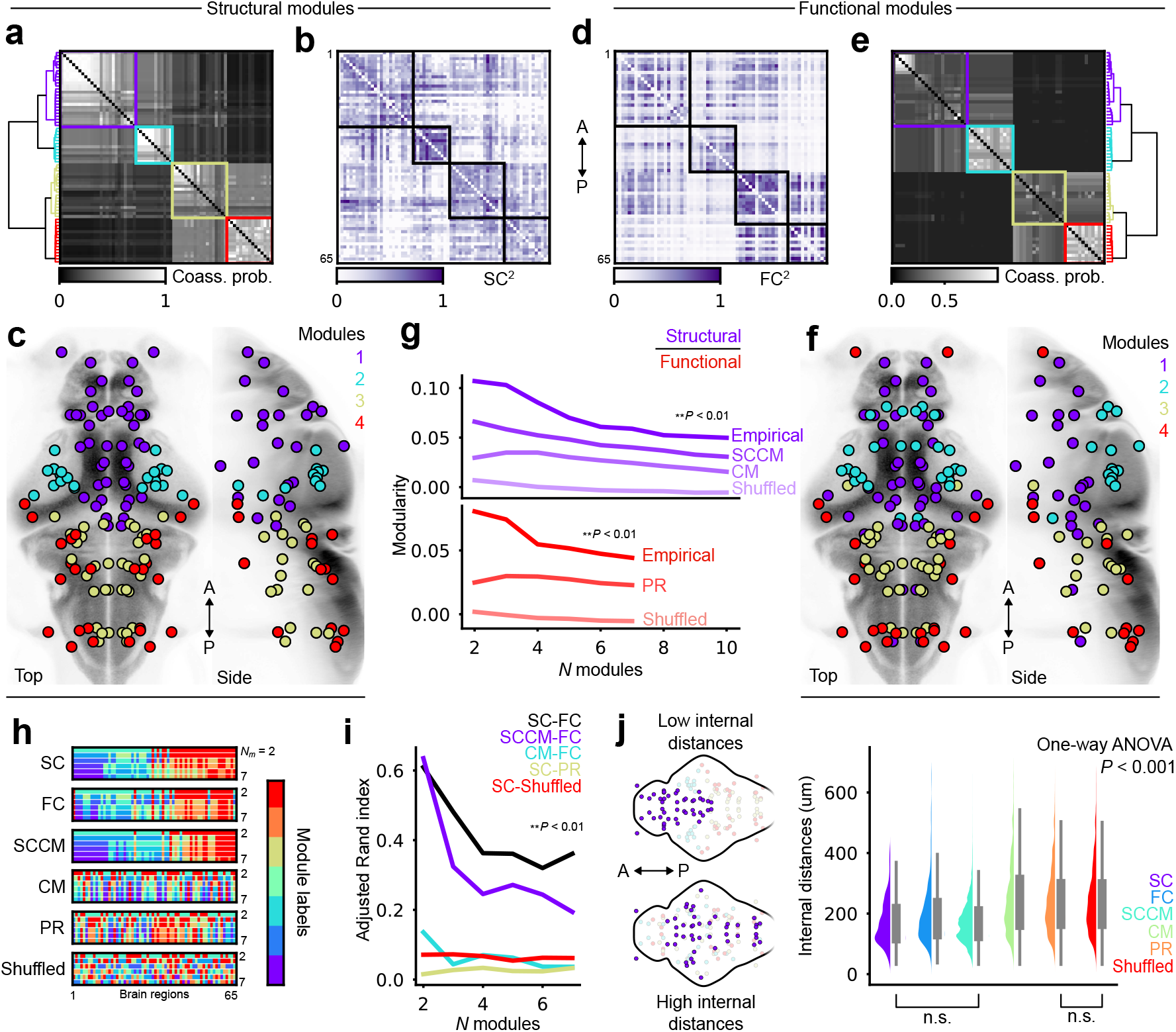
Structural and functional modules are overlapped at multiple hierarchical levels. (**a, d**) Module coassignment probability matrices, hierarchically clustered using Ward’s linkage (for SC and FC, respectively); 4 modules are chosen arbitrarily as an intermediary hierarchical level for visualization. (**b, e**) Connectivity matrices, reordered by modules according to the previous panels (SC and FC, respectively); colors are mapped to squared edge weights for better contrast. (**c, f**) Brain region centroids colored by module membership (SC and FC, respectively). (**g**) Structural and functional modularity at different hierarchical levels, compared against various null models. SC modularity curves are plotted above (purple), and FC curves below (red); 99th percentiles of null distributions are plotted for null models; SCCM, spatially constrained configuration model; CM, configuration model; PR, phase-randomized time series (see Methods). (**h**) Module identity vectors plotted at different hierarchical levels, for both empirical data and null models; *N*_*m*_ denotes the number of modules; representative labels are shown for each null model. (**i**) Adjusted Rand index computed on pairs of module labels from the previous panel at different hierarchical levels; 99th percentiles of null distributions are plotted. (**j**) Left, visual examples of spatially compact (top) vs distributed (bottom) modules; right, boxplots of pairwise internal distances between regions belonging to the same modules, compiled across all hierarchical levels; all distributions are different with the exception of two comparisons, as indicated below the figure (one-way ANOVA, *P <* 0.001, followed by Tukey’s post-hoc test for pairwise comparisons).

**Figure 4:**
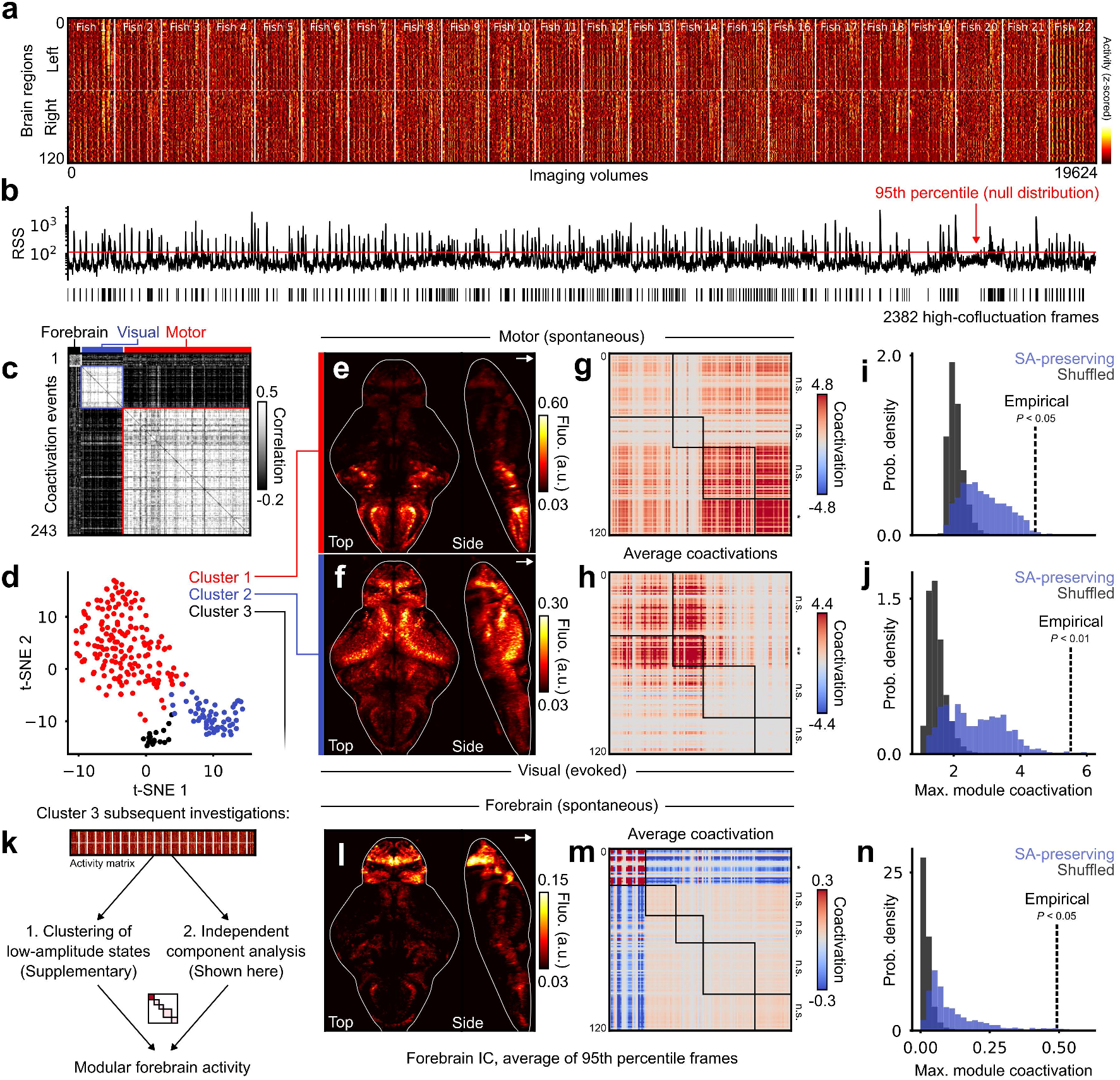
Mesoscopic co-activation patterns coincide with the modular organization of SC. (**a**) Raster plot of z-scored regional activity across 22 larvae. Left and right hemispheres are plotted in the upper and lower halves, respectively. (**b**) Root-mean-square (RMS) of regional co-activation values for each of the corresponding frames in panel (**a**); the red line indicates a statistical threshold for high co-activation events (*P <* 0.05, false-discovery rate fixed at *α* = 0.05 using the Benjamini-Hochberg procedure). (**c**) Correlation matrix of 243 detected events, separated into 3 clusters that are illustrated in the following panel. (**d**) t-SNE projection of 243 high-amplitude events, with 3 clusters identified using density-based clustering. (**e**-**f**) Raw data projections of the two main clusters, co-registered and averaged across larvae (*n* = 22); top, high-amplitude motor events; bottom, high-amplitude visual responses to dark stimuli; contrast is adjusted after averaging, see Methods. (**g**-**h**) Average co-activation matrices of the two main clusters, reordered according to 4 structural modules; statistical significance of module co-activations is indicated on right matrix borders. (**i**-**j**) Null distributions of maximal module co-activation values in SA-preserving and SA-breaking surrogates; dashed lines indicate empirical co-activation values. (**k**) Two independent analyses recover modular forebrain activity that was weakly detected as cluster 3 in the previous clustering analysis. (**l**) Averaged raw fluorescence from frames corresponding to the 95th percentile of forebrain independent component (IC) activity. (**m**) Average co-activation matrix of frames belonging to the 95th percentile of forebrain IC activity. (**n**) Null distributions of forebrain modularity, similar to (**i**-**j**).

### Co-activation patterns of brain regions are shaped by structural modules

Recent modeling and neuroimaging studies in humans and mice suggest that FC is shaped by the sum of a small number of short-lived and large-amplitude regional co-activation events ^20,68^, which seem to emerge from the modular organization of the structural connectome ^21,22^. Using the hierarchy of modules established in the previous section, we examined whether large-amplitude events were present in zebrafish brain activity and whether they spatially aligned with structural modules. To identify large-amplitude activity patterns, we *z*-scored regional time series across both hemispheres (Fig. 4**a**) and then quantified the co-activation of brain regions by computing the root-mean-square (RMS) of pairwise node co-fluctuations at each time point ^20^ (Methods). This calculation yielded a signal whose peaks indicate moments where multiple brain regions are either strongly coactivated or co-deactivated relative to their means (Fig. 4**b**). We detected 243 high-amplitude events in 22 animals, whose amplitudes significantly exceeded null RMS distributions obtained through circular permutations of regional time series (*P <* 0.05, false-discovery rate adjusted at *α* = 0.05). We projected all events onto a 2D plane using the dimensionality reduction algorithm t-SNE ^69^, then used the density-based clustering algorithm HDBSCAN ^70^ to identify three main event clusters (Fig. 4**c-d**, Supp. Fig. S13 for cluster reproducibility, Methods). The largest cluster consisted of high activity in hindbrain regions, coinciding with episodes of motor activity (Fig. 4**e**, raw data projection, Methods). The second largest cluster consisted of high activity in the optic tectum and other visually responsive regions following dark stimuli after the spontaneous imaging period (Fig. 4**f**). The third substantially smaller and less coherent cluster corresponded to spontaneous periods of elevated activity in forebrain regions (Supp. Fig. S13). We focused our analysis on the first two clusters to ask whether these activity configurations were modular by reorganizing functional co-activation matrices according to previously identified SC modules (Fig. 4**g**-**h**) and then by computing the average co-activation levels within each module (Methods). We found that each event cluster had a single module that was significantly co-activated compared to null spatial surrogates of regional activity that approximately preserved spatial autocorrelation ^71^ (SA) (Fig. 4**i**-**j**, *P <* 0.05 for motor, *P <* 0.01 for visual; 1000 surrogates per distribution). We conducted additional validations to evaluate the significance of module co-activations across the full hierarchy of SC modules (Supp. Fig. S14**a**-**b**-**d**-**e**) and to ensure that statistical significance was unlikely when substituting empirical SC modules with null modules derived from SCCM matrices (Supp. Fig. S14**c**-**f**).

We next investigated the third forebrain cluster, which involved fewer co-activated brain regions and was therefore in-appropriately detected by a global high-amplitude threshold. Using two separate approaches (Fig. 4**k**), we recovered spontaneous configurations of high forebrain activity. First, we applied hierarchical consensus clustering on the remaining low-amplitude frames to retrieve a significantly modular forebrain cluster (Supp. Fig. S15). Second, we used independent component analysis ^72^ to identify a forebrain component (Methods). We used peaks in its associated time course to identify frames during which the component displayed high activity (95th percentile of forebrain IC activity). During these “forebrain events”, contrastingly low activity was observed throughout the rest of the brain (Fig. 4**l**, average of the 95th percentile of IC activity, *n* = 22 larvae). By reorganizing the average forebrain co-activation matrix with SC modules, we again observed striking modularity (Fig. 4**m**), with co-activation values in the forebrain module significantly exceeding those of spatial surrogates (Fig. 4**n**, *P <* 0.05). Surprisingly, significant co-activations within this module were preserved in null SCCM modules (Supp. Fig. S14**i**), likely a consequence of the protruding geometry of the forebrain that prevents local forebrain connections from being properly shuffled in null models where distance constraints are enforced on edge swaps, thus preserving this module after randomization (Supp. Fig. S8**d**-**f**, see top left matrix corners).

Taken together, these results suggest that high-amplitude co-activations associated with motor, visual, and forebrain activity (rendered in Supp. Video 3) involve brain regions that are strongly interconnected within SC modules, thus replicating and extending recent neuroimaging results from mammals ^20,22^ to zebrafish.

### Neurons associated with visual and motor functions collectively span most of the larval brain

With visual and motor functions as the main drivers of coordinated inter-regional activity, we next took advantage of cellular resolution data to further characterize neurons underlying these functions at a microscopic scale. First, we used high-speed tail recordings and machine vision to track different tail segments of head-restrained larvae, which exhibited smooth undulations during swimming (Fig. 5**a**,**b**, Methods). We then trained a recurrent neural network to automatically identify these swimming events across experiments (Fig. 5**c**, Methods; *n* = 18 larvae with tail recordings). In parallel, we probed visually responsive neurons using abrupt light-dark transitions (6s dark followed by 9s light, repeated 12 times; viewable in Supp. Video 2). To identify neurons significantly correlated with swimming or visual responses, we convolved motor events or stimulus vectors with an exponential kernel (*τ* = 3.5s) to generate hypothetical calcium responses (regressors), and then correlated these fictive signals with real neuronal time series ^73,74^. We applied different statistical thresholds to identify four neuronal populations in each animal: motor-correlated cells (motor positive), motor-anticorrelated cells (motor negative), dark-correlated cells, and light-correlated cells (*P <* 0.01, thresholds inferred from null correlation distributions, see Methods; example activity traces in Fig. 5**e**). We compiled these cells in all larvae in 3D atlas coordinates, then applied a spatial statistical threshold to exclude cells found in locations where few cells from the same category were observed across individuals (spatial *P* -value test ^75^, *P <* 0.025, Methods). This procedure generated four consensus spatial densities which, together, spanned most of the brain (Fig. 5**d**, overlapped densities; Fig. 5**f**, individual densities; Supp. Fig. S19, brain atlas regions; Supp. Fig. S17, unthresholded densities).

**Figure 5:**
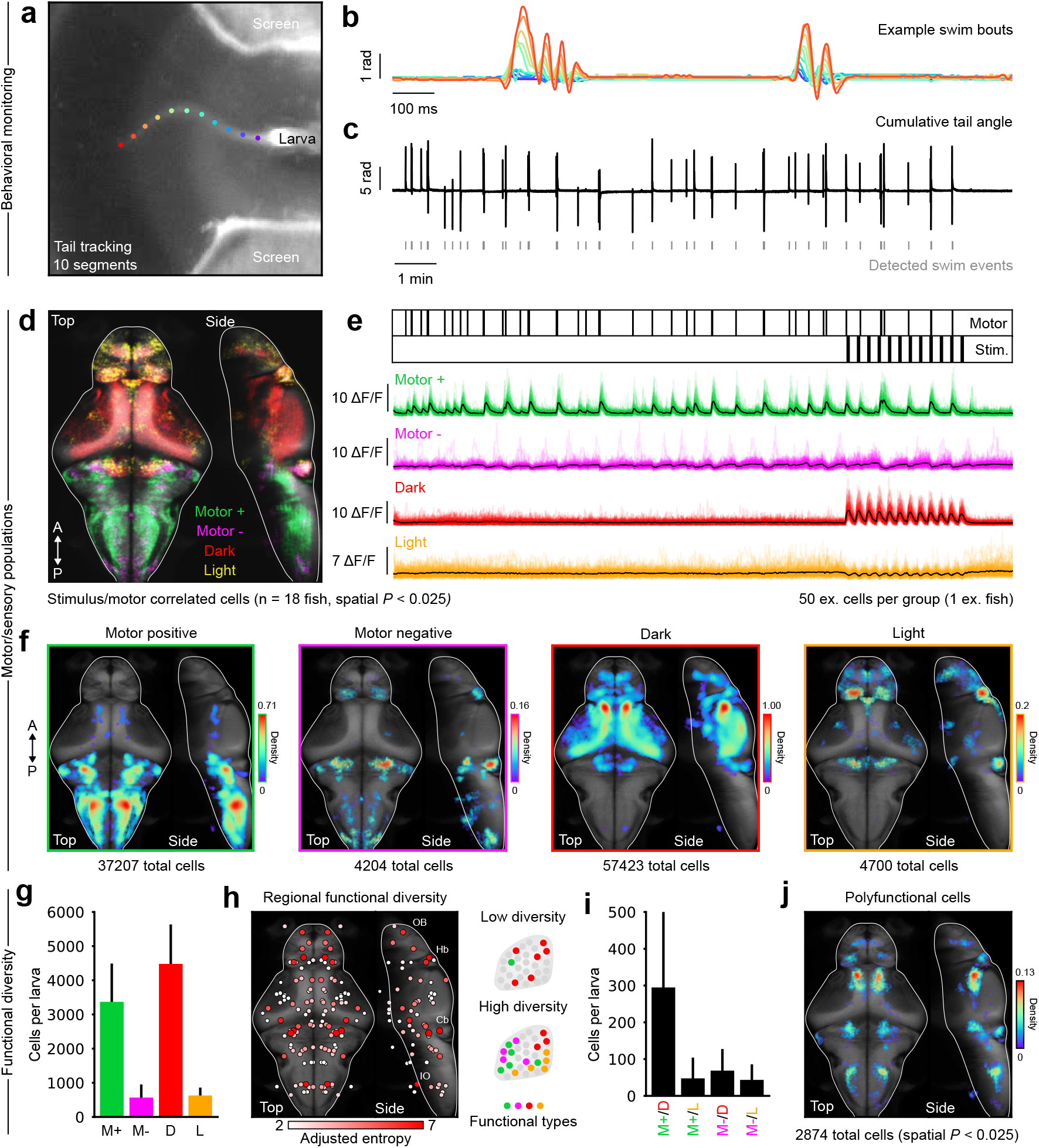
Identification of sensorimotor neuronal populations. (**a**) Example frame from high-speed monitoring of a head-restrained larva with tail tracking points. (**b**) Angular time series of 10 tail segments displaying two successive swimming events; colors correspond to tracking points on the previous panel. (**c**) Cumulative tail angle of one larva over a full experiment, with detected swimming events below. (**d**) Overlapped densities of motor-positive, motor-negative, dark- and light-responsive cells, projected on mapZebrain template brain (*n* = 18 larvae, spatial *P <* 0.025. (**e**) Top, motor and stimulus event vectors; bottom, 50 example cells per functional category (color traces) with average population activity (black traces). (**f**) Individual cell densities from panel **d**; the colormap represents arbitrary spatial density units, relative to the maximum density observed in dark-responsive cells; the number of cells aggregated across animals and contributing to each distribution is indicated below each panel; spatial *P <* 0.025. (**g**) Number of cells per functional category per larva; error bars indicate standard deviation; statistical comparisons are avoided due to slight methodological differences between motor and visual cell identification (see Methods). (**h**) Left, regional measure of functional diversity using adjusted Shannon entropy; nodes reflect region centroids, and node sizes are redundant with colormap. Right, schematized examples of low and high diversities in a brain region. (**i**) Number of overlapping cells for each non-orthogonal pair of functional categories per larva; horizontal graph labels combine the colors from previous panels; error bars indicate standard deviation. (**j**) Spatial density of polyfunctional cells detected across all animals, spatial *P <* 0.025. A, anterior; P, posterior; M+, motor-positive; M-, motor-negative; D, dark; L, light.

In each animal, we detected 3369 ± 1085 motor-positive neurons, 567 ± 347 motor-negative neurons, 4480 ± 1120 dark-responsive neurons, and 628 ± 195 light-responsive neurons (mean ± standard deviation, Fig. 5**g**). Motor-positive neurons were found broadly throughout the hindbrain, while motor-negative neurons formed multiple small and segregated clusters throughout the hindbrain, cerebellum and habenula (Fig. 5**g**, green and magenta). We used noradrenergic (NE) markers from mapZebrain to contextualize some of the motor negative clusters (Supp. Fig. S18). Notably, some overlapped with NE neurons within the locus coeruleus (Supp. Fig. S18**a**) and the medulla oblongata (putative NE-MO cells ^76^, Supp. Fig. S18**b**), consistent with the known functional role of these cells, which initiate the freezing behavior in head-restrained preparations ^76,77^. We also observed a sharp laminar organization of positive and negative motor cells in the caudal hindbrain (Supp. Fig. S18**c**), suggesting that the negative correlations were not motion artifacts as they mapped on well-delineated anatomical substrates ^78^. Unlike motor populations, most dark and light neurons were located within the anterior brain, in the optic tectum, cerebellum, habenula, and forebrain regions (Fig. 5**g**, red and orange). Strikingly, light-responsive neurons were mostly concentrated within the left habenula (Fig. 5**f**), consistent with previous observations ^79^. A detailed anatomical quantification of these populations within brain atlas regions is provided in Supp. Fig. S19.

Although we observed very little overlap between motor and visual populations, a few regions such as the cerebellum contained cells from all four categories. To quantify functional diversity between brain regions, we calculated the entropy ^80^ from the regional proportions of functional cell types (Supp. Fig. S19**b**), normalized by the standard deviation of this measure across individuals, thus penalizing poorly or unevenly sampled regions (Methods). This adjusted entropy measure was highest in the cerebellum, habenula, forebrain, midbrain, anterior hindbrain, and inferior olive regions (Fig. 5**h**), most of which are known to integrate signals from multiple modalities ^81–83,34,84–86^. We also identified cells that belong to numerous functional categories at once (polyfunctional neurons), corresponding in majority to both motor-positive and dark-responsive cells (Fig. 5**i**). Polyfunctional cells were sparse, found within only a few nuclei across the brain (Fig. 5**j**, spatial *P <* 0.025; separate overlap densities are shown in Supp. Fig. S20).

Overall, our analyses of sensorimotor populations uncover numerous functionally specialized clusters that collectively span a considerable fraction of the larval brain (about 84% of brain regions, Supp. Fig. S19**b**; rendered in Supp. Video 4). While some regions and individual cells integrate signals from different functional modalities, visual and motor populations remain mostly non-overlapping at both cellular and regional levels, a key observation for the subsequent analysis.

### Visuomotor populations coincide with the main functional network gradient

We identified visual- and motor-correlated neurons located sequentially along the brain’s anteroposterior axis, an anatomical layout which perhaps reflects the flow of sensorimotor transformations along the larval brain. Many recent neuroimaging studies have shed light on the contiguous spatial organization of brain functions by investigating functional gradients (or connectopies ^87^) ^23,88,89,24,90^. These gradients, derived from either structural or functional connectivity measurements, indicate where each node is situated along the network’s major axes of connectivity variance, with nearby gradient values indicating nodes with similar connectivity profiles with other brain regions (Methods). Connectivity gradients have been shown to gradually unfold across visual, motor, and transmodal cognitive regions, providing a data-driven mapping of the brain’s processing streams and hierarchies ^91,23^.

Using our previous identification of visual and motor neurons, we next investigated if connectivity gradients of SC and FC could reflect the sensorimotor organization of the zebrafish brain. We calculated connectivity gradients using the diffusion map algorithm ^92^ on both FC and SC matrices, yielding eigenvectors that represent the principal axes of random diffusion on the network (Methods). Both networks were characterized by a main diffusion gradient that opposed anterior and posterior brain (Fig. 6**a**, gradient 1, Pearson *r* = 0.795 between SC and FC gradient 1). Focusing on the first 50 SC and FC gradients, whose eigenvalues (i.e. their relative importance) quickly decayed (6**b**,**d**), we established an optimal linear mapping between SC and FC gradients, as similar gradients were found in a different order within both networks (Fig. 6**c**, Methods). We then calculated the average correlation between SC and FC gradients (mean of matrix diagonal in Fig. 6**c**), which significantly exceeded mean correlations with null connectivity gradients (*P <* 0.001, 1000 null SCCM matrices, data not shown) or with spatially shuffled gradients (*P <* 0.001, 1000 SA-preserving permutations, data not shown). We further evaluated the significance of single gradient comparisons, observing 7*/*50 significantly correlated gradient pairs under the null connectivity hypothesis (*P <* 0.05, 1000 SCCM gradients per gradient pair) and 46*/*50 significantly correlated pairs following SA-preserving permutations (*P <* 0.05, 1000 spatial permutations per gradient pair). The main anteroposterior FC and SC gradients were significantly correlated to each other under both of these conditions (*P <* 0.001 and *P* = 0.026 for SCCM and spatial permutations, respectively). Thus, SC and FC network gradients are significantly correlated at the ensemble level, and while not all gradients have a significant 1-to-1 mapping, generally similar connectivity gradients can be detected within both networks, in particular their main gradient which unfolds from anterior to posterior brain.

**Figure 6:**
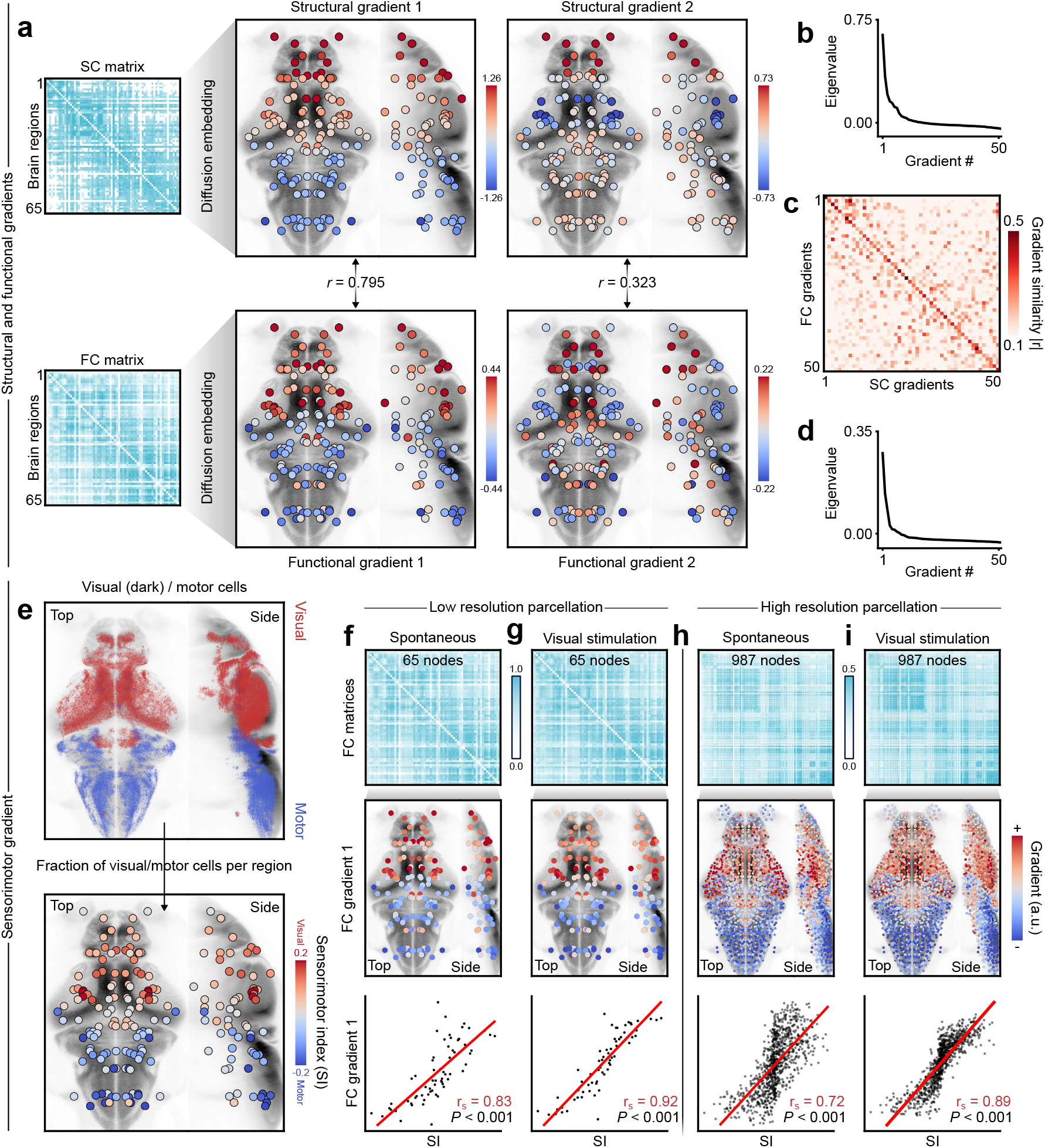
Regional sensorimotor functions coincide with the main functional gradient. (**a**) Top, SC matrix and its first two diffusion gradients, with nodes denoting brain region centroids; bottom, FC matrix and its first two diffusion gradients that match the first two structural gradients. (**b**) Eigenvalues of the first 50 structural diffusion gradients. (**c**) Absolute correlation |*r*| between the first 50 SC and FC gradients; functional gradients are reordered to optimally match the structural gradients. (**d**) Eigenvalues of the first 50 functional gradients. (**e**) Top, dark and motor positive neuron centroids from Fig. 5, in red and blue respectively; bottom, sensorimotor index of each brain region. (**f**) Top, FC matrix in spontaneous conditions; middle, first FC gradient of the corresponding matrix above; bottom, correlation between sensorimotor index (SI) and first FC gradient; *r*_*s*_ denotes the Spearman correlation coefficient, and *P*-values are obtained through SA-preserving permutations (1000 permutations). (**g**) Similar to the previous panel, except with FC computed while including visual stimuli. (**h**) Similar to the previous panel, except with high-resolution FC computed from spontaneous activity. (**i**) Similar to the previous panel, except with high-resolution FC computed while including visual stimuli.

Next, we used the two larger functional populations identified earlier, dark and motor positive cells, to annotate the sensorimotor affiliation of brain regions and investigate the cellular basis of the principal gradient (Fig. 6**e**). We computed the average fraction of dark and motor-positive neurons per region, and then subtracted these quantities to obtain a “sensorimotor index” where positive values indicate visual regions, and negative values indicate motor regions (Methods). Fig. 6**f** shows that this sensorimotor index strongly and significantly correlated with the main FC gradient (Spearman’s *r* = 0.85, *P <* 0.01, 1000 SA-preserving surrogates, Supp. Fig. S21**a**). The main SC gradient was also highly, but not significantly correlated with the sensorimotor index (Spearman’s *r* = 0.778, *P* = 0.073, 1000 SA-preserving surrogates). The mapping increased when we derived gradients from FC calculated over the full experiment duration, capturing both spontaneous and visually stimulated activity (Fig. 6**g**, Spearman’s *r* = 0.93, Supp. Fig. S21**c**, *P <* 0.01). The spatial smoothness of this FC gradient prompted us to further validate its correspondence with visuomotor populations at a higher network resolution by fractioning anatomical brain regions into smaller and roughly equal parts (Methods). Despite a tenfold increase in the number of network nodes, correlations between the first FC gradient and the sensorimotor index remained high and significant (Fig. 6**h**-**i**, Spearman’s *r* = 0.68 for spontaneous FC, *r* = 0.89 for visually stimulated FC; Supp. Fig. S21**c**-**d**, both *P <* 0.01).

We observed another functional gradient which mostly accentuated forebrain regions in all FC matrix definitions (Supp. Fig. S22**b**). Along with the first visuomotor gradient, the two diffusion eigenvectors formed a roughly triangular and contiguous embedding of brain regions with visual, motor, and forebrain regions occupying each corner (Supp. Fig. S22**c**-**d**). Similar embedding geometries have been observed in mammals ^23,24^, with visual, motor, and transmodal regions separated by the two main connectivity gradients. This has been interpreted as a hierarchy of brain regions, with unimodal sensory regions at the basis and transmodal associative regions at the apex ^23^. The forebrain gradient observed here could reflect a separation from sensorimotor to putative transmodal regions, in qualitative agreement with the numerous complex functions housed within the zebrafish forebrain (see Discussion).

Together, these results demonstrate that the main functional role of brain regions, either visual or motor, coincides largely with the principal functional gradient of larval zebrafish FC. This gradient is observed in both spontaneous and visually stimulated functional networks, regardless of their spatial resolutions. Other gradients, such as the forebrain functional gradient, likely capture other functions not covered by our experiments.

### Functional connectivity is predicted by regional gene expression levels

Our results so far highlight how multiple features of functional brain networks and whole-brain activity can be traced back to an underlying structural network architecture. The previous experiments and analyses, however, are based on an indistinct pan-neuronal fluorescent labeling, thus omitting the cellular diversity present in the developing larval brain. Alongside SC, genetic factors are crucial determinants of brain function, influencing both cellular diversity and network-level organization ^19^. Their influence is apparent in studies combining gene expression atlases with brain connectivity measurements, with transcriptomic similarities in specific genes related to neurotransmission generally predicting higher functional connectivity across brain regions ^93,94^. To investigate the influence of gene expression on larval zebrafish FC, we leveraged a recent dataset of spatially resolved gene expression markers ^33^. We compiled 290 fluorescence *in situ* hybridization RNA expression markers (HCR ^95^) from mapZebrain ^33^ (five examples in Fig. 7**a**), then quantified their regional expression levels from normalized intensities of regional marker fluorescence (Fig. 7**b**, Methods). Next, we computed the pairwise correlations of regional gene expression profiles (the rows of Fig. 7**b**), yielding a co-expression matrix (correlated gene expression, CGE) that we then compared with the FC matrix. We observed a modest correlation between CGE and FC (Spearman’s *r* = 0.33, Fig. 7**c**, left), suggesting that transcriptomic profiles evaluated across the full spectrum of genes do not predict functional network organization.

**Figure 7:**
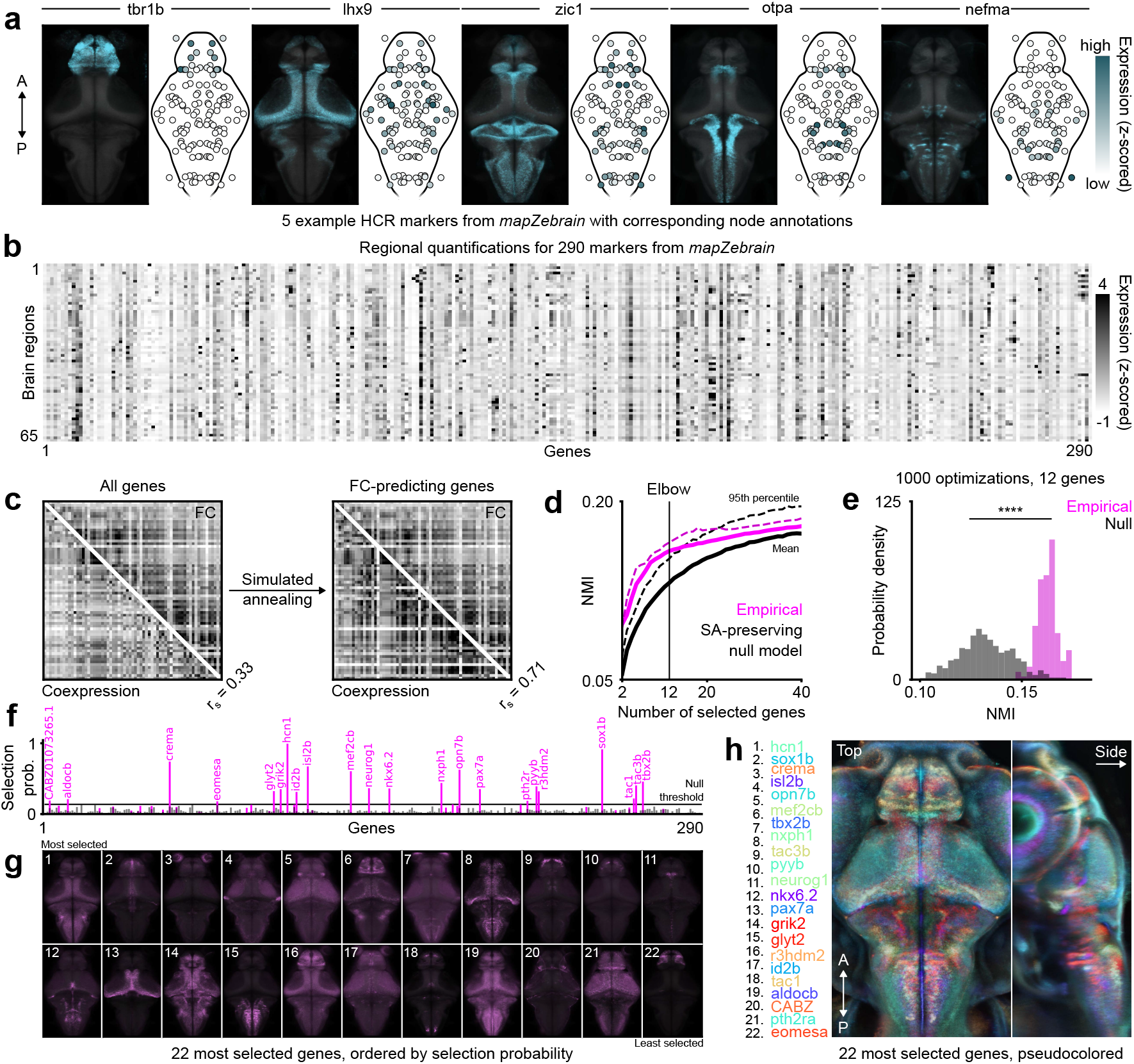
Gene co-expression predicts functional connectivity. (**a**) Five example fluorescence *in situ* hybridization markers from mapZebrain (left, top view, 90th percentile projections), with corresponding regional annotations (right, in black outlines); intensities are scaled arbitrarily per gene for visualization. (**b**) Regional gene expression matrix; each row (brain region) is z-scored independently. (**c**) Left, CGE matrix across all 290 genes, compared with the FC matrix; *r*_*s*_ denotes Spearman’s coefficient; right, CGE matrix of 12 optimized genes obtained through simulated annealing, compared with FC. (**d**) Average (full lines) and 95th percentile (dotted lines) of optimized normalized mutual information (NMI) between CGE and FC for varying numbers of genes used in simulated annealing runs; pink curves correspond to empirical gene sets, while black curves correspond to spatially shuffled genes; a vertical line indicates the elbow of the average empirical curve; 1000 optimization runs per number of genes. (**e**) NMI distributions at the predetermined elbow point for empirical and null optimization results using 12 genes (*P <* 0.001, t-test). (**f**) Selection frequency of each gene across 1000 simulated annealing runs; pink bars correspond to empirical genes, while black bars correspond to spatially shuffled gene datasets; a horizontal bar denotes statistical significance, with significant gene names indicated above. (**g**) 90th percentile intensity projections of 22 significant gene markers; gene names are indicated in the next panel, ordered by selection probability (left to right, top to bottom). (**h**) Overlap of 22 pseudocolored gene markers; pixelwise color contrast is used to highlight most prominent genes at each location; colors are used to accentuate the global patterns, rather than to precisely distinguish individual genes (which are displayed separately in the previous panel). A, anterior; P, posterior; SA, spatial autocorrelation.

We next searched for smaller subsets of genes whose co-expressions were more strongly correlated with FC, using a simulated annealing optimization approach ^96,94^ with normalized mutual information (NMI) as the optimization criterion (Methods). Across a wide range of parameters, we identified optimized subsets of genes whose CGE matrices were highly correlated with FC (Fig. 7**c**, right, one example of optimized CGE matrix with Spearman *r* = 0.71). To find an optimal gene set size, we performed thousands of optimizations while computing CGE from groups of 2 to 40 genes, observing progressively diminishing returns in performance for sets larger than 12 genes (Fig. 7**d**, elbow method, 1000 optimizations per number of genes). To evaluate the statistical significance of these results, we generated 1000 surrogate gene expression matrices by independently permuting markers while preserving their spatial autocorrelation ^71^, and then performed similar optimizations on shuffled gene datasets. For small numbers of genes, the real dataset significantly outperformed the surrogates in predicting FC (Fig. 7**d**, dotted curves indicate the 95th percentile, thus *P <* 0.05; Fig. 7**e**, NMI distributions for optimal gene set size, *P <* 0.0001). Note that for larger gene sets (*>* 22), the null model eventually outperformed the real dataset, likely a consequence of the higher spatial redundancy of real unshuffled genes (Fig. 7**d**, Supp. Fig. S23).

After determining the optimal gene barcode size, we identified which genes most frequently appeared in the optimal solutions. A comparison of 1000 optimization runs using 12 genes on both unshuffled and shuffled datasets revealed 22 genes that were consistently selected more frequently than others (Fig. 7**f**). These gene markers are shown in Fig. 7**g**, and overlayed on the zebrafish brain projection in Fig. 7**h**, highlighting their spatial distribution and complementarity. The most selected gene was *hcn1*, which encodes a voltage-gated potassium channel expressed mostly in the hindbrain. The second-most selected gene was *sox1b*, a transcription factor involved in neurodevelopment expressed mostly in forebrain and midbrain structures, but also in the spinal cord ^97^. The functions of these 22 genes, sorted by their selection probabilities, are listed in Supp. Table 2. Some are directly involved in neurotransmission, such as the presynaptic transporter of glycine (*glyt2*), found within hindbrain glycinergic populations, while others encode functions that do not directly relate to neuronal activity (see Discussion).

**Table 2:** Brief descriptions of the functions of the 22 most selected genes from simulated annealing optimizations, ordered by selection probability.

| Gene | Selection % | Function |
| --- | --- | --- |
| hcn1 | 99.4 | Hyperpolarization-activated cation channel; voltage-gated potassium channel activity |
| sox1b | 91.4 | DNA-binding transcription factor |
| crema | 74.1 | DNA-binding transcription factor |
| isl2b | 67.3 | DNA-binding transcription factor |
| opn7b | 62.6 | G protein-coupled receptor activity and photoreceptor activity |
| mef2cb | 60.5 | DNA-binding transcription factor |
| tbx2b | 45.8 | DNA-binding transcription factor |
| nxph1 | 43.9 | Neurexophilin; signaling receptor binding activity; promotes adhesion between dendrites and axons |
| tac3b | 40.4 | Regulation of blood pressure |
| pyyb | 39.1 | Neuropeptide receptor binding activity |
| neurog1 | 36.6 | Enables E-box binding activity and protein homodimerization activity |
| nkx6.2 | 36.2 | DNA-binding transcription factor |
| pax7a | 35.3 | Transcription factor |
| grik2 | 35.1 | Glutamate ionotropic receptor kainate type subunit 2 |
| glyt2 | 32.4 | Glycinergic synaptic transmission |
| r3hdm2 | 32.1 | Enables RNA binding activity. |
| id2b | 31.2 | Negative regulation of transcription by RNA polymerase II |
| tac1 | 22.0 | Substance P receptor binding activity |
| aldocb | 21.1 | Fructose 1,6-bisphosphate metabolic process and glycolytic process |
| CABZ01073265.1 | 19.1 | Unknown |
| pth2ra | 18.7 | Receptor for parathyroid hormone |
| eomesa | 18.0 | DNA-binding transcription factor |

Overall, these analyses indicate that the co-expression of a small number of genes at a regional level in the larval zebrafish brain can predict FC, suggesting an important role for regional cellular properties in the organization of functional brain networks.

## Discussion

Decades of research combining MRI and network analysis have solidified our understanding of the functional and structural architectures of the human brain in both health and disease ^98,99^. Here, we used larval zebrafish as a strategic vertebrate model system to study brain networks at a brain-wide regional scale ^29^ – similar to the scale of neuroimaging studies – while gaining access to cellular detail within the same set of experiments. We adapted methods from mammalian brain studies to zebrafish and, despite broad differences in brain size and complexity, obtained results consistent with those reported in larger animal models.

We first used whole-brain two-photon calcium imaging to demonstrate that the high-dimensional activity of over 50,000 cells can be reduced into a single correlation matrix, which, despite severe coarse-graining, is a very stable and meaningful observable of neurodynamics. Indeed, we measured similar FC across different datasets, microscopy techniques, and experimental settings. We also observed a high similarity between the FC of different larvae, in line with previous reports ^35^. Although larval FC is similar between specimens, we were able to discriminate individual larvae imaged on separate occasions using their FC fingerprint, similarly to humans ^49^ and mice ^100^. Although more experiments are required to demonstrate the ability to recognize individuals within a larger population and to benchmark functional and anatomical fingerprinting methods (Supp. Fig. S5**c**), this is to our knowledge the first demonstration of subject identification using brain activity in zebrafish. This result might not be so surprising, however, as larvae exhibit diversity in body morphologies and growth rates, locomotion levels ^101,102^, stimulus responsiveness ^103^, and learning abilities ^104^, among numerous traits that distinguish them. Some of this diversity likely arises from neurological differences that reflect the different genetic backgrounds of each animal ^103^. Our experiments highlight the high sensitivity of FC in recognizing the neurological fingerprints of zebrafish larvae undergoing rapid development, which will be instrumental in future studies investigating whole-brain fingerprints of neurologically altered subjects after mutations ^105,38^, pharmacological interventions ^36^, or environmental stresses ^106,107^. While statistical comparisons between animals and experimental conditions are facilitated by mesoscopic brain parcellations, analyzing such datasets at the single-cell level remains the ultimate goal. Whole-brain imaging in zebrafish provides a unique opportunity to bring network analyses closer to the cellular scale, where a much greater level of detail comes at the price of a substantially higher computational burden. Delineating finer brain parcellation based on, for instance, neurotransmitter domains within regions might strike a good balance between interindividual comparability and a more fine-grained definition of functional network nodes.

Using single-cell morphologies compiled independently into a single 3D volume ^32^, we generated an inter-regional wiring diagram and studied the structure-function relationship of brain networks. There are several limitations to this SC estimation method: (1) undersampling, as the size of the dataset represents about 4% of the total number of neurons; (2) a lack of discrimination between axons and dendritic arbors; (3) the absence of pre- and post-synaptic information; and (4) the absence of neurotransmitter identity. We benchmarked different wiring methods based on the spatial layout of neurites, but the SC matrices generated here remain imperfect and incomplete approximations of inter-regional connectivity. Different methods have been proposed to estimate connectivity from the spatial relationships of independent tracing experiments ^108,109^. Until complete electron microscopy reconstructions of the nervous systems of larval zebrafish ^110,111^ and larger models ^112^ are achieved, these approaches will remain valuable for studying connectivity at a mesoscopic, neural populations level. By estimating inter-regional pathways, we observed a strong coupling between structural and functional networks. Direct anatomical connections partially predicted FC, and numerous network properties that account for indirect signaling mechanisms were even more predictive of FC. A recent study in C elegans exploiting optogenetics to photostimulate every single neuron ^53^ remarkably demonstrated the cumulative effects of synaptic, polysynaptic, and extracellular communication on driving correlations between neurons. Such exhaustive experiments are not yet feasible in larger nervous systems, but optogenetic approaches can be used to probe single neurons and anatomical pathways in zebrafish and infer connectivity ^113–115^. Correlative approaches based on graph properties can also provide hypotheses on the relative importance of different signaling policies (e.g. direct vs diffusive routing) across the entire brain at regional scale. Pharmacological or genetic perturbations are promising avenues to alter whole-brain FC and evaluate how the relative contributions of different FC predictors change, providing more mechanistic insight on the factors that modulate the structure-function coupling ^14^.

We found a modular architecture in both functional and structural networks, with significantly overlapped network modules at different resolution levels. This finding suggests that previously identified functional modules ^35^ are anchored by a similar anatomical layout of structural modules. Furthermore, by resolving instantaneous regional co-fluctuations, we observed motor, visual, and forebrain activity patterns that coincided significantly with modules identified within the connectome. This finding aligns with recent modeling ^21^ and neuroimaging ^22^ results, where brief large amplitude events ^20^ emerge from densely interconnected modules. While our results emphasize a clear neuronal origin to these events, their underlying mechanisms in mice, humans, and zebrafish remain unclear. Here, large-scale motor and visual activity patterns were directly related to locomotion and vision, but elevations in forebrain activity were not overtly associated with stimuli or ongoing behavior. It is unclear whether this spontaneous activity reflects the intrinsic recurrent dynamics of the forebrain or if external inputs drive such fluctuations. Our modularity analysis identifies the forebrain as a densely interconnected structure, in agreement with recent higher-resolution connectivity analyses conducted on the same structural dataset ^109^. Thus, an appropriate substrate for the synchronization of neuronal populations ^116,21^ appears to be present within the zebrafish telencephalon, but more studies are required to untangle the mechanisms of spontaneous forebrain fluctuations and those of high-amplitude modular activity in general. Our observations reinforce the well-established hypothesis that connectome modules are functionally specialized subnetworks ^7^. Future connectivity studies at the nanoscale ^110,111^ should provide a much better subdivision of this hierarchical structural architecture, allowing a finer mapping of structure to function ^117,118^ in the zebrafish brain.

We identified a main functional gradient from visual- and motor-correlated neurons, which we annotated at a regional level using proportions of functional cell types. Dark-flash stimuli elicit robust behavioral responses in larvae ^40^ and were used here to drive visual responses in large neuronal populations ^34^. However, zebrafish larvae boast a wide array of hardwired responses to other stimuli ^47,48^, which future studies should incorporate to assess how various sensorimotor populations coincide with functional connectivity gradients and other network topological properties, such as modules. From a limited set of stimulus and motor regressors, we identified brain regions boasting a high functional diversity, but a larger array of stimuli would also benefit this analysis and provide important brain annotations to dissect the structural pathways underlying various sensorimotor transformations. For simplicity, we limited our regression analysis to four main functional cell categories, which were enough to contextualize the main connectivity gradient, but several functional and motor subpopulations were most likely hidden within these broad categories. For instance, we combined all swimming events into a single motor regressor and neuronal population, but separate motor populations in the hindbrain are associated with different swimming types ^119,120^. While diverse locomotor patterns can in principle be observed in head-restrained larvae under the microscope, most swimming events observed here under spontaneous conditions were large escape swims. We noted heightened activity in putative NE-MO cells during motor quiescence (Supp. Fig. S18), which are known to initiate freezing behavior when visual feedback becomes disconnected from motor activity^76^. Several limitations encountered here could be addressed by introducing additional stimuli and providing larvae with visual feedback in a closed-loop system. Nevertheless, dark-flash stimuli were sufficient to uncover a principal visuomotor gradient of whole-brain activity observed even in the absence of stimulation. This perhaps reflects the phylogenetic importance of darkness, a prominent feature of natural aquatic habitats ^121^ which numerous studies have harnessed to generate stereotyped responses in behavioral assays ^122,123^. In addition to this first gradient, we detected a second FC gradient which separated visual and motor populations from the forebrain. This finding evokes earlier questions: what gives rise to modular forebrain activity, and what functional roles are associated with the forebrain gradient? Using meta-analyses in humans, the main FC gradient has been associated with cognitive functions of increasing complexity, solidifying its interpretation as a unimodal-transmodal axis of the brain ^23^, while the second gradient coincided with sensorimotor regions. In zebrafish, the sensorimotor axis seems to be the main driving force behind FC, which perhaps reflects the early developmental stage of the animals considered in our study. Standardized data on zebrafish cognitive functions are lacking, precluding a quantitative assessment of the forebrain gradients observed here. However, many functions are housed within the zebrafish telencephalon, such as social interactions ^124^, learning ^125–128^, and spatial navigation ^129^, to name a few. Therefore, based on this diversity of functions, it may be appropriate to consider the forebrain FC gradient as a putative transmodal axis of the zebrafish brain, but more efforts are required to support this idea. As the field grows to accommodate more sophisticated experimental designs during brain imaging ^130,131,75,129^, the brain-wide neural correlates of complex and naturalistic behaviors should yield precious data to gain a more quantitative understanding of the functions associated with numerous architectural features of the zebrafish brain, including its major connectivity gradients.

While anatomical connectivity is perhaps the most fundamental constraint on brain dynamics, gene expression profiles strongly dictate the functional properties of cells and neuronal populations ^19^. In support of this idea, we identified 22 genes that, collectively, were significant predictors of FC. This observation suggests that neuronal populations with similar transcriptomic profiles are more likely to be correlated in their activities, perhaps by sharing many ion channels, receptor subtypes, and electrophysiological properties. Although our simple quantification scheme did not take full advantage of HCR markers due to resolution limitations, empirical gene expression levels significantly outperformed spatially shuffled genes in predicting FC. As the dataset used in this study is small relative to the full genome, the optimal barcode size determined here is likely to grow when replicating similar analyses on a larger database. Some identified genes are directly associated with neurotransmission, while others, such as transcription factors, may control the expression of proteins that regulate neuronal activity ^132^, and dictate the fate, type, and function of cells ^133^. Meanwhile, genes that may not appear related to neuronal function could be correlated in their expressions with other activity-related genes that are not present within the dataset. Future studies investigating the genetic basis of FC would benefit from combining spatially resolved gene expression data with larger single-cell RNA sequencing databases to explore a much broader range of genes and their interactions. An important issue to note is that genes expressed in one location can, in some cases, have functional outcomes elsewhere in the brain ^105^. The gene combinatorial approach used here identifies a regional transcriptomic barcode that predicts time-averaged correlations at the gene ensemble level, but direct comparisons between activity patterns and single gene expression maps should be approached carefully. We do not expect these approaches to be capable of appropriately pinpointing specific mutations, but function-altering mutations to individual genes are expected to have a significant impact on FC, as shown in other studies ^36–38^. Network-based approaches and whole-brain functional imaging are promising tools for enriching studies of single-gene mutations/knockouts in zebrafish, which have relied primarily on behavioral and histochemical assays to characterize neurological effects ^31,134,105,135^. Furthermore, comparing how various mutations, associated with a specific neurological or psychiatric condition, individually impact on FC measurements could reveal whether common traits in brain-wide FC may underlie these conditions. It should be noted that while the structural wiring of the brain is also coordinated by gene expression, the prediction of SC from gene expression requires more intricate rules than simple correlation-through-co-expression ^136^ which goes beyond the scope of our study that focused mainly on FC. As structural and genetic datasets grow to match the scale and resolution of whole-brain functional imaging data, more exhaustive studies on the reciprocal interactions between these three different factors will become possible at a cellular level.

In summary, this work demonstrates that many brain network phenomena observed at much larger neuroimaging scales — often through indirect hemodynamic measurements — can be paralleled in a much smaller nervous system using cellular-resolution imaging, providing evidence for their generality and reproducibility across species. Moreover, it highlights striking functional similarities between the nervous system of teleost fish and that of mammals. With a wide array of experimental tools to observe and manipulate neural circuits, the zebrafish model offers abundant opportunities for future connectomics studies to unravel the intricate relationship between structure, function, and genetics within brain networks.

## Methods

### Animal model

All experiments were conducted on 5-7 dpf Tg(*elavl3* :H2B-GCaMP6s) ^39^ zebrafish larvae. Larvae were raised in embryo medium (13.7mM NaCl, 0.54mM KCl, 1.0mM MgSO_4_, 1.3mM CaCl_2_, 0.025mM Na_2_HPO_4_, 0.044mM KH_2_PO_4_, 4.2mM NaHCO_3_, pH 7.2 with HCl 12N) at a density of 1 larva per mL in Petri dishes placed in an incubator at 28°C on a 14:10 hour day/night cycle. From 5 dpf onward, the medium was replaced daily and larvae were fed live *Tetrahymena thermophila* CU428.2 (Cornell University). *T. Thermophila* were grown in Petri dishes in sterile SPP medium (2% proteose peptone, 0.1% yeast extract, 0.2%glucose, 33 *µ*M FeCl_3_, 250 *µ*g/mL Penicillin/streptomycin, 0.25*µ*g/mL Amphotericin B) at room temperature and harvested in the stationary phase. They were then washed from their SPP medium using three centrifugation steps (2min at 0.8g each), followed by pellet resuspension in embryo medium. All protocols were approved by the animal care committee of Université Laval.

### Two-photon microscopy

Larvae were embedded in 2% low-melting point agarose (Invitrogen 16520100) in a 30 mm glass bottom petri dish (Mattek P35G-1.0-14-C) and positioned upright for brain imaging using a toothbrush bristle without direct contact with the animal. Agarose was carefully removed below the swim bladder and around the tail using a scalpel (Fisher 35-205), and then the preparation was submerged in embryo medium and placed under the microscope for imaging. All imaging experiments were conducted on a Scientifica SliceScope resonant two-photon microscope (SciScan LabView software) equipped with a Spectra Physics Insight X3 tunable laser. Fluorescence was collected using a piezo-driven 16x water-dipping objective (Nikon, 0.8 NA), reflected by a T585lpxr dichroic mirror, and bandpass filtered (505/119nm) before detection with a GaAsP detector. High-resolution anatomical stacks were first collected (512 × 720 pixels, 200 planes, 2 microns in z-spacing, 24x frame averaging) using an excitation wavelength of 860 nm, which is near the isosbestic point of the calcium indicator. Following the structural scan, whole-brain calcium imaging was performed at an excitation wavelength of 920 nm. We sampled brain activity from 22 imaging planes (512 × 720 pixels) spaced approximately 12 microns apart, and excluded the first plane due to piezo recoil artifacts induced by the sawtooth scanning pattern. We delimited the imaging volume by positioning the first plane at the top of the brain where labeled nuclei are first encountered, and then positioned the last plane in ventral neuronal populations, slightly above the lower boundary of the pan-neuronal expression pattern. All images were obtained with a lateral pixel resolution of approximately 0.9 × 1.1 microns (width × height), which is below the average nucleus diameter of approximately 5 microns. Power measurements after the objective were kept at 20 mW, and functional imaging experiments lasted 15 minutes. For the small dataset of 7 larvae used for fingerprinting experiments (6-7 dpf, 2 imaging sessions per animal), as well as the older datasets used for reproducibility analysis in Supp. Fig. S3, we recorded spontaneous brain activity for 10-30 minutes in a slightly truncated field of view, spanning the same set of brain regions with the exception of a few caudal hindbrain regions.

### Tail monitoring and tracking

Tail movements were monitored from below the imaging chamber at 400 frames per second using a high-speed camera (Flir Blackfly S USB3, BFS-U3-28S5M-C) mounted with an adjustable macro lens (Navitar Zoom 7000). 850 nm LED arrays were used to illuminate the fish at a 45 degree angle from above. Infrared light was sent through a dichroic mirror, and then reflected and band-pass filtered (Thorlabs, 852nm center wavelength, 10nm width) into the camera. A 4x binning factor was used to increase the signal-to-noise ratio of the camera frames, resulting in small 160 × 160 pixel images which were sufficiently resolved for accurate tail tracking. The camera was triggered and synchronized with the microscope using an Arduino and a custom-written Bonsai ^137^ workflow.

To track and reconstruct the tail in each frame, we used a standard intensity-based tracing algorithm based on previously published methods ^138,139^. First, we used a custom GUI to remove the background from the video by subtracting the intensity mode (most frequent intensity value) of each pixel computed along the temporal axis. Then we applied Gaussian filtering and local contrast enhancement (CLAHE) with varying parameters if required, depending on the quality of each recording and the size of each larva. Then, we manually selected a base point for the tail located in the swim bladder of the fish and used 10 segments of equal length *r* to reconstruct the rest of the tail. Due to variations in the length of the fish, the length of the tail segments was manually adjusted for each animal. The initial tail segment was obtained by sweeping a 120-degree cone in small increments of 6 degrees in the expected direction of the tail (in this case, left) and then placing a tracking point at a distance *r* following the angle where the intensity of the image is highest along a line profile. The subsequent tracking points were obtained similarly and iteratively by sweeping a 120-degree cone from the previously defined tail point, centered on the direction of the previous segment, and by placing a new point in the direction of maximal intensity. The coordinates of each tracking point were further refined at every step by computing the local intensity center of mass within a small 5 pixel window, yielding an improved sub-pixel resolution for tracking coordinates and more appropriately centering each point on the tail.

After reconstructing the tail, the angles of each tail segment were computed at every frame, relative to the main axis of the larval body (in this case the horizontal axis). This yielded 10 angular time series capturing the detailed movement of each tail segment. Angular time series were unwrapped to avoid discrete jumps from − *π* to *π* and vice-versa, and then their means were subtracted to ensure that the resting position of the tail is 0. Angular time series were finally filtered to remove sparse tracking glitches by applying a median filter twice consecutively (window of 3 time steps or 7.5 ms), and then applying a Gaussian filter (*σ* = 1 time step).

### Detecting swim events

To identify swim events from tail tracking signals, we manually segmented the onset and offset of hundreds of swim bouts in 7 animals and then used these annotations to train a gated recurrent unit (GRU) neural network ^140^ with 64 hidden units using Pytorch ^141^. The model received 10 angular time series as input, and we thresholded its probability output at to retrieve binary time series, where 1 indicates an ongoing swimming event and 0 indicates no swimming event. We trained the model using 9000 temporal segments of length 800 time steps (roughly 2 seconds of recording) and a batch size of 64, and we mirrored left and right swimming events to augment the dataset, for a total of 18000 training examples. 25% of the training examples were empty tracking sequences without swimming events (mainly noise) to ensure that the model performs well on lengthy and sparse tracking data. We trained the model for 20 epochs using the binary cross entropy loss function, the Adam optimizer ^142^ and a learning rate of 10^−3^. Once trained on a few animals, we used the model to segment the rest of our swimming data. We found this method to be more robust and less sensitive to rare tracking glitches and noise spikes, which other threshold-based methods often identified as false positives and had to be filtered out later.

Note that we delayed the model output by 15 time steps (37.5 milliseconds) relative to the manual annotations during training to provide a temporal integration window. Because swimming events take a few time steps to reach non-negligible amplitudes, a small integration window is required for the model to disambiguate swimming from noise and to confidently detect ongoing events. Once trained and deployed on new data, we shifted the model’s outputs by 15 time steps in the opposite direction to retrieve accurate onset times.

### Visual stimulation

Visual stimuli were delivered using red light (AAXA Technologies KP-750-00 DLP Projector with a Kodak dark red #29 Wratten 2 filter) projected under the imaging chamber using a dichroic mirror (Thorlabs). We utilized standard white printing paper secured against the glass of the imaging chamber as a diffusive screen for projection. The screen spanned the entire visual field of the larvae, with the exception of a small cut-out region to allow tail monitoring. Larvae were centered in the chamber during agarose embedding to ensure that both eyes received equal illumination. Infrared light was used for tail monitoring to avoid interference with the visual system. For spontaneous brain activity recordings, we maintained larvae under constant illumination that spanned the entire visual field. For dark-flash stimulation, we alternated between 6s dark periods and 9s light periods, repeated 12 times, which is equivalent to 3 minutes of periodic visual stimuli. Stimuli were rendered in `.avi` video format using Python and triggered by the microscope using a custom-written Bonsai ^137^ workflow.

### Calcium imaging data processing

Each calcium imaging recording was pre-processed to correct unwanted lateral drifts and movements in the imaging chamber, and to automatically extract the activity of all recorded cells. First, we converted raw `16-bit` microscopy data into 21 separate `.tif` imaging planes, and then corrected lateral motion in each imaging plane independently using the piecewise non-rigid registration algorithm NoRMCorre ^143^ (implemented in Python within the CaImAn package ^42^). We used a large patch size of 120 × 120 pixels with a 60 pixel overlap to minimize local registration artifacts, which can occur when correcting motion in peripheral regions of the brain where few landmarks are present in smaller patches. Experiments where significant z-drift was observed (e.g. cells moving out of focus) were discarded.

After motion correction, we segmented fluorescent nuclei from the temporal averages of the corrected imaging planes using a simple maxima detection algorithm ^43^. Briefly, the algorithm identifies local intensity maxima as putative nuclei in the raw image, and merges these maxima with those identified from a second processed version of the image where local contrast is enhanced. First, average calcium frames were smoothed using a Gaussian filter (*σ* = 0.5), and then thresholded using the triangle threshold method to exclude background regions. Then, local maxima were identified using the `peak_local_max` function from `skimage` ^144^, with a minimum distance between neighboring maxima corresponding to the lower bound of the expected radius of nuclei (2 pixels). A second set of maxima was then extracted independently from a locally normalized version of the image, where each pixel is divided by the maximum value found in neighboring pixels within a small disk of radius 2 pixels, again corresponding to a lower bound on the expected size of a nucleus. The normalized image was thresholded using the background found previously on the raw image, and smoothed with a similar Gaussian kernel before finding maxima. To merge the maxima obtained from both complementary images, pairwise distances were computed between both sets of maxima, and then the nearest neighbors were identified. Neighbors from the second set that were found within a radius of 4 pixels from any maxima of the first set were assumed to be associated with the same nucleus and thus discarded. Maxima from the second set that were located further away than this merge radius were considered as newly identified cells that had been missed in the original image and incorporated into the final set of maxima.

After finding the coordinates of putative nuclei, the fluorescence signals *F* (*t*), where *t* represents time (s), were extracted from a small disk of 2 pixel-radius centered on each maximum, corresponding to a conservative lower bound on the expected radius, thereby ensuring no overlap between cells. We excluded pixels belonging to rare cases of overlapping regions of interest to ensure no overlap between neighboring nuclei. Raw fluorescence traces were then detrended for slow baseline drifts and converted to relative fluorescence change values, Δ*F/F*_0_, using the following sequence of operations. First, to identify the temporal baseline *F*_0_(*t*) of each neuron, which typically decay due to photobleaching, calcium signals were Gaussian filtered t(*σ* = 3s), and raw baseline values were calculated from the filtered signals using a minimum filter with a large and conservative temporal window of 300 seconds. The baselines were then smoothed using a wide Gaussian filter (*σ* = 60s), and finally used to calculate the detrended raw signals using the following equation:

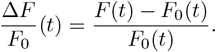

We opted for a conservative baseline removal strategy with wide temporal windows to avoid subtracting rapid physiological fluctuations.

### Brain atlas registration

To map the 3D coordinates of segmented cells from different larvae into a common 3D referential, we used a two-step registration process. First, we used Advanced Normalization Tools (ANTs ^44^) to compute a sequence of three transforms (an initial rigid transform, followed by an affine transform, and finally a non-rigid transform using the SyN algorithm) that map the 21 piezo imaging planes, acquired using an excitation wavelength of 920 nm, to their higher-resolution anatomical counterpart, acquired at 860 nm. We used a set of parameters that was optimized for live brain registration in a previous study ^145^:

~~~
antsRegistration -d 3 --float 1 --output [,$filename_aligned] --interpolation WelchWindowedSinc --use-histogram-matching 0
-r [$filename_fixed,$filename_moving,1] -t rigid[0.1] -m MI[$filename_fixed,$filename_moving,1,32,Regular,0.25] -c [200×200×200×0,1e-8,10]
--shrink-factors 12×8×4×2 --smoothing-sigmas 4×3×2×1vox -t Affine[0.1] -m MI[$filename_fixed, $filename_moving,1,32,Regular,0.25]
--convergence [200×200×200×0,1e-8,10] --shrink-factors 12×8×4×2 --smoothing-sigmas 4×3×2×1vox -t SyN[0.2,6,0]
-m CC[$filename_fixed,$filename_moving,1,2] -c [200×200×200×200×10,1e-7,10] --shrink-factors 12×8×4×2×1 --smoothing-sigmas 4×3×2×1×0vox
~~~

This sequence of transforms uses mutual information as the alignment metric for the rigid and affine transforms, and cross-correlation for the non-rigid SyN transform. `$filename_fixed` corresponds to the reference image stack saved in `.nrrd` format with pixel sizes indicated in the file header, and `$filename_moving` is the image that gets warped (in this case, the functional imaging stack). Second, we computed similar transforms from the high-resolution anatomical stack acquired at *λ* = 860 nm to a 6 dpf template brain of the same nuclear GCaMP line from mapZebrain. We visually inspected each transformed stack, overlapped onto its target stack, to ensure that no considerable registration errors were made at any of the two stages. Finally, the 3D coordinates of neurons in the original calcium imaging planes were written to a .*csv* file in physical units (microns), and transformed into atlas coordinates by applying the `antsApplyTransformsToPoints` function twice, first from the functional stack to the anatomical stack coordinates, then from the anatomical stack to the brain atlas coordinates. After transforming cell coordinates into the brain atlas referential, we converted them back into pixel coordinates, and then mapped each cell into one of 70 anatomical brain regions by indexing their corresponding 3D binary masks. All masks were downloaded from mapZebrain and compressed into a single sparse `.csz` file for efficient storage and access. We used the 1.0 version of brain region masks, which differs slightly from the current 2.0 region masks.

### Computing functional connectivity

Before computing FC matrices from the calcium imaging data, we first identified a few regions to exclude from the analysis. We excluded region 11 (retina), which does not contain accessible fluorescent neurons due to the opacity of the eyes. We also excluded all regions that contained less than 25 segmented cells in at least half of the larvae, leading to the exclusion of four additional regions: region 35 (pituitary), region 45 (caudal hypothalamus), region 50 (glossopharyngeal ganglion), and region 70 (lateral reticular nucleus). All brain regions are indicated in Supp. Table 1, with exclusions highlighted in orange.

We averaged the calcium fluorescence signals over all segmented neurons in each brain region and then generated 65 × 65 FC matrices for each larva by computing the Pearson correlation coefficients between pairs of region signals. We slightly Gaussian filtered the regional time series (*σ*=2s) before computing correlations. We then proceeded to compute the average FC matrix by averaging the functional connectivity values across all individual FC matrices. Because a few larvae had excluded brain regions due to insufficient sampling, some FC edges were computed on a subset of all 22 animals. However, the majority of FC edges were computed on the full population and we did not account for these slight statistical heterogeneities throughout subsequent analyses.

We note that atlas brain region masks were drawn across both hemispheres and that we used this regional definition without splitting nodes into separate hemispheres. Some analyses use bi-hemispheric networks, roughly doubling the number of nodes, such as the fingerprinting analysis in Fig. 1 (see next section), or the analysis of whole-brain co-activation patterns in Fig. 4. However, most of this study uses combined hemispheres. Therefore, all figures except Fig. 4 use left-right mirrored edges in network representations, as well as mirrored node annotations (for instance, network modules) for ease of visualization.

### Comparing FC across datasets

To validate the reproducibility of FC in zebrafish larvae, we compared the current dataset with older data generated in our lab, using similar imaging parameters and a slightly reduced field of view. We used identical data pre-processing and mapped these experiments in the same anatomical parcellation before computing FC as described previously. Due to the reduced field of view, we truncated the dataset used in this study to encompass the 58 anterior-most regions. To evaluate the similarity of both FC matrices, we correlated their vectorized upper triangles. We repeated this procedure with previously published lightsheet microscopy data ^34^, which was initially mapped in a different brain atlas (Z-Brain atlas ^31^). We used ANTs with the parameters described above to compute a transform from Z-Brain to mapZebrain by registering the *elavl3* :H2B-RFP and *elavl3* :H2B-GCaMP6s template brains to each other, and then transformed lightsheet cell coordinates using the `antsApplyTransformsToPoints` function. We detrended calcium signals, then computed regional group-average FC matrix from the spontaneous periods, which were interspersed throughout the lightsheet experiments during which multiple visual stimuli were delivered ^34^. To properly match the fields of view of both datasets, we truncated them to encompass the 53 anterior-most regions.

### Comparing FC across individuals

To measure the inter-individual similarity of FC, we used upper triangle correlations between all pairs of FC matrices, ignoring unsampled (`NaN`) values. This correlation matrix between individuals is shown in Fig. 1**g**. We evaluated the magnitude of these similarities by comparing the distribution of inter-individual similarities with two null models (Supp. Fig. S4). The first one was generated through temporal shuffling by applying cyclic permutations of the regional time series before computing FC. This generated 22 null FC matrices where correlations are lower and reflect randomized signals, while preserving the frequency spectra of the original time series. We compiled null inter-individual similarities by repeating this process 25 times. For the second null model, we used spatial shuffling of regional centroids to establish a new rostro-caudal order of the matrix, thereby preserving its values, but permuting them. Briefly, for each larva, we generated a random 2D embedding of regional centroids using t-SNE, roughly preserving neighborhoods but randomly rotating and reordering the rostro-caudal axis. Then we reordered the matrix rows/columns on the basis of the vertical axis of this embedding, and finally computed the inter-individual similarity scores. We repeated this process 25 times. Both null models yielded significantly lower similarity values than the empirical inter-individual FC similarities (one-way ANOVA, *P <* 0.0001).

### FC fingerprinting

To study the individuality of FC, we imaged 7 larvae on two consecutive days, at 6 and 7 dpf. Larvae were embedded in agarose as described above, and whole-brain calcium imaging was conducted for 30 minutes under constant red illumination. Larvae were then carefully removed from the agarose embedding and left overnight in individual petri dishes filled with embryo medium. The following day, each larva was embedded and imaged again in the same order as the day prior. We computed FC matrices across both brain hemispheres, effectively doubling the network size. We kept only the brain regions that were properly sampled in both hemispheres and across all larvae, for a total of 58 regions and 116 network nodes. We performed pairwise comparisons of FC matrices from day 1 to day 2 and vice versa, yielding two asymmetrical matrices that were averaged to generate one symmetrical 7 × 7 similarity matrix, encoding the similarity of two individuals’ FC regardless of the temporal direction in which datasets are being compared. The maximum values on each row were observed on the diagonal in 6*/*7 cases, most individuals being maximally similar to themselves. We alternatively used a global signal regression to evaluate how arbitrary signal processing steps can impact identification accuracy. For each animal, we computed the average brain activity from regional time series and then performed a linear fit of this signal onto each individual regional time series before subtracting it and computing regressed FC matrices. This operation globally lowered the similarity scores between all pairs of FC matrices, but maintained (and even increased) the identification rate. We evaluated statistical significance by randomly shuffling identity labels in one of the dataset before computing pairwise similarities and counting the number of row-wise maximal similarities found along the matrix diagonal.

### Computing structural connectivity from single-cell morphologies

We tested four methods to generate mesoscopic SC matrices from a dataset of 4327 single cell morphologies ^32^, and then compared these methods against each other by correlating them with the FC matrix (Pearson correlation between the upper triangles of SC and FC matrices, *r*_*F C*_). We detail these connection weighing schemes in Supp. Fig. S7. In all four cases, we generated raw connectivity matrices *C* iteratively, one neuron at a time. For each neuron, we first identified the presynaptic region *j* from the location of the neuron’s soma, then inferred the target regions *i* from its neurites, and finally evaluated the raw connection weights *C*_*ij*_ using different morphological quantities and strategies that we describe hereafter. For Method 1 (Supp. Fig. S7**a**), we attributed connection weights using the length of neurites passing through *any* brain regions, while for Method 2 (Supp. Fig. S7**b**), we restrained weight counts only to regions where the neuron’s terminals are present, thereby excluding those where neurites are only passing through. Once compiled across all cells, the final weights of these two methods reflect the total neurite length that one region is sending through and into another.

For Method 3 (Supp. Fig. S7**c**), we weighed connections by counting the number of terminals within each downstream region, while for Method 4 (Supp. Fig. S7**d**), we slightly expanded the regional boundaries to encompass additional terminals (see Supp. Fig. S6), and then counted the terminals similarly as per Method 3, only with slightly larger and overlapping regions. Once compiled, the final weights of these two methods reflect the total number of terminals that one region establishes into another. The raw directed connectivity matrices, which sum these quantities across all neurons, are displayed in Supp. Fig. S7**e**-**h**. We converted these raw directed matrices into volume- and log-normalized, directed or undirected *W* matrices using the following relationships

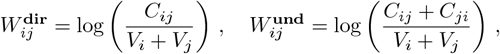

where *V*_*i*_ indicates the volume of brain region *i*. This normalization procedure ensured that larger regions, such as the periventricular layer of the optic tectum, did not get disproportionately large weights simply by geometrically intercepting more neurites and terminals, or due to the larger number of cells present in the region. Following this normalization, we observed significantly log-normal weight distributions in all four cases (Supp. Fig. S7**i**-**l**, *P >* 0.05 in all cases, Kolmogorov-Smirnov test). We used the undirected versions of these matrices to compare them with FC (Supp. Fig. S7**i**-**l**, upper triangles shown). We found that counting terminals was slightly better at predicting FC than the summed wiring length in a brain region (*r*_*F C*_ = 0.473 for wiring length in all downstream intersected regions, *r*_*F C*_ = 0.466 for wiring length in downstream regions with terminals, *r*_*F C*_ = 0.485 for number of terminals, Supp. Fig. S7**i-k**). We also found that expanding regional boundaries increased the correlation with FC (*r*_*F C*_ = 0.568 for Method 4, Supp. Fig. S7**l**). Therefore, we prioritized terminal count with slightly expanded regional boundaries (30 micron expansion) for this study.

### Validation of wiring method

To validate the SC weighing scheme, we evaluated the effect of regional boundary expansion on FC predictability. As mentioned previously, we expanded regional boundaries to recover multiple unmapped terminals (Supp. Fig. S6), as well as to mimic the spatial extent of dendritic arbors of postsynaptic cells, which are entirely unknown in the structural dataset. We reasoned that dendrites are not confined to their somatic region and could establish synapses in adjacent regions or within neighboring neuropil areas. Thus, terminals located outside one region could still contribute to its connectivity count, although this becomes less likely with increasing distance. Overall, the regional expansion procedure increased the correlation between SC and FC, peaking at an expansion magnitude of 30 microns (Supp. Fig. S7**m**). Larger expansion magnitudes eventually decreased this correlation, perhaps reflecting a 30 micron optimum that captures the average spatial extent of the presumptive dendritic arbors. We validated that this gain in FC predictability could not be replicated by randomly distributing the same amount of extra terminals uniformly in the raw matrix *C* before computing *W*, or by shuffling the additional normalized connection weights once *W* is computed (Supp. Fig. S7**n**-**o**, 1000 null matrices per shuffling method, *P <* 0.001). These validations ensured that the increased correlation with FC is not trivially driven by increasing the density of the sparse SC matrix. We note that this method of estimating inter-regional connectivity almost certainly increases the number of false positives, which should, however, be rivaled by a decrease in false negatives. However, this trade-off can not precisely be quantified without ground truth connectivity data.

### Null models of structural connectivity

Throughout this study, we assessed the significance of many real network properties using a spatially constrained configuration model (SCCM) of SC, which is identical to the standard configuration model (CM) ^146^, with the addition of a simple distance constraint on edge swaps. In the case of CM, a graph is randomized through a sufficiently large number of double edge swaps, each swap being selected at random, through which each edge is swapped at least once, but ideally at least 10 times. This procedure preserves node degrees, but randomizes the topology of the network. In the case of the SCCM, we performed a similar edge swapping procedure, with the exception that some swaps were prevented when the difference in length of each new edge relative to the original edge length exceeded a certain threshold, which we set to 30 microns. We found that this threshold offered a good compromise between preserving many properties of the original network without being too restrictive and limiting exploration of new random network configurations (see Supp. Fig. S8). We generated 1000 null SC matrices using 50000 edge swaps each, more than 10 times the number of edges in the network. We repeated this process for both directed and undirected networks and used this null matrix ensemble throughout the rest of the study. For standard CM, we removed the distance threshold and generated a similar amount of null matrices. Both of these null models preserved exactly the weight distribution of the SC matrix and the binary node degrees. Additionally, we constrained swaps to occur only at the presynaptic nodes in the directed case, which preserved exactly the weighted input degree distribution.

### Structural predictors of functional connectivity

In Fig. 2, we used a wide range of structural properties to predict FC. Some studies have used the properties of SC, encoded in dense matrices, to predict FC^89^. However, the set of properties used as predictors in these studies is typically limited, and the few that are selected then become arbitrary. Based on a previous study ^58^, we aimed to eliminate such selection bias by evaluating a large number of properties. We provide summary descriptions of each predictor used and refer to network neuroscience references that include extensive lists of network properties for alternative descriptions and citations to the original sources ^147,148,58^.

In all cases, the predictors follow a similar nomenclature in the labels of Fig. 2**h** and **i**. First, the name of the predictor (for example, `euc` or `nav`), followed by weighted (`wei`) or binary (`bin`), and finally the numerical value of a single parameter, such as a matrix inversion parameter *γ* or Markov time *t* used to compute the predictor. All predictors were computed from the full directed or undirected SC (*W*) matrix (70 × 70), and then the five excluded brain regions were truncated to match the size of the FC matrix for comparisons. We used the Python version ^149^ of the Brain Connectivity Toolbox ^147^ (`bctpy`) to compute most of these predictors. Some predictors required a “cost” matrix *C*, rather than connections *W*, where small weights are associated with high costs, and large weights are associated with low costs. This mapping can be achieved simply by inverting the weights of the SC matrix *W*_*ij*_ using the formula 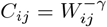, where the *γ* parameter allows different weight-to-cost transformations.

### Euclidean distance

Euclidean distance *D* between brain regions (more precisely, between regional centroids) was first calculated within and between hemispheres and then averaged to yield a single representative distance between two regions across both hemispheres. Then, the distance matrix *D* was inverted element-wise using 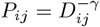 to produce a proximity matrix *P* where high values indicate nodes that are close to each other in space. We used *γ* = {0.125, 0.25, 0.5, 1.0}, generating 4 predictors that are annotated as `euc-0.125` to `euc-1.0` in Fig. 2.

### Shortest path length

We computed binary and weighted shortest paths from different directed cost matrices *C* with different values of *γ* using Dijkstra’s algorithm. Then, we inverted and symmetrized the shortest path length matrix element-wise to obtain a predictor matrix that is positively correlated with FC. For the binary case, a weight-to-cost transformation was not required, and therefore a single binary path length matrix was computed. Cost matrices were generated with *γ* = {0.125, 0.25, 0.5, 1.0, 2.5, 5.0}, yielding a total of 7 predictors that are annotated as `pl-bin`, followed by `pl-wei-0.125` to `pl-wei-5.0` in Fig. 2.

### Communicability

Communicability evaluates how two regions exchange information using a weighted sum of all parallel paths between two nodes, thereby encapsulating a more diffusive signaling process. In the binary, undirected case ^59^, it is calculated as *G* = *e*^*W*^ using the standard eigendecomposition procedure. In the weighted, undirected case ^60^, the connectivity matrix is normalized using 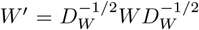, where *D*_*W*_ is the diagonal matrix of node strengths (sum of weights for each node), before being exponentiated, resulting in *G*_*wei*_ = *e*^*W*′^. These two predictors were calculated using undirected SC and are annotated as `comm-bin` and `comm-wei` in Fig. 2.

### Cosine similarity

Cosine similarity measures the cosine of the angle between two vectors and is used here to evaluate the similarity between the connectivity neighborhoods of two brain regions, defined as the rows (inputs) or columns (outputs) of the weighted, directed connectivity matrix *W*. Given two *n*-dimensional vectors, *υ*_1_ and *υ*_2_, it is calculated as the normalized scalar product 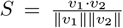. This measure was applied to directed SC to assess the similarity of both inputs and outputs between all pairs of brain regions. The 2 predictors are annotated as `cs-in` and `cs-out` in Fig. 2 for inputs and outputs, respectively.

### Matching index

The matching index ^61^ quantifies the similarity between the connectivity profiles of two nodes by measuring the proportion of shared connections relative to their total number of connections. If Γ_*k*\*ℓ*_ denotes the set of all nodes connected to node *k*, excluding node *ℓ*, then the matching index between modes *i* and *j* is calculated as 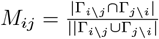, where | |, ∩, and ∪ denote, respectively, the cardinality (number of elements) of a set, the intersection, and the union of sets. Extended to weighted networks ^50^, the matching index is defined as 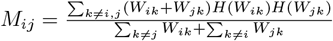, where *H* is the Heaviside step function such that *H*(*W*_*ij*_) = 1 if *W*_*ij*_ *>* 0 and 0 otherwise. We computed this measure on directed, weighted matrix *W* using both inputs (rows) and outputs (columns) of brain regions. The resulting predictors are annotated as `mi-in` and `mi-out` in Fig. 2, respectively.

### Flow graph

A flow graph ^62^ is represented by a weighted connectivity matrix 𝒲, where edge weights encapsulate information about a diffusion process occurring between pairs of nodes of on an original graph with weighted connectivity matrix *W*. When the process is a continuous-time random walk, the flow between nodes *i* and *j* at time *t* is defined by 𝒲 (*t*)_*ij*_ = (*e*^−*tL*^)_*ij*_*d*_*j*_, where *L* is the normalized Laplacian *L*_*ij*_ = *δ*_*ij*_ − *W*_*ij*_*/d*_*j*_, *d*_*j*_ = ∑_*k*_ *W*_*ij*_ is the strength of node *j*. We used both binary and weighted undirected matrix *W* to calculate this measure, at *t* = {2.5, 5.0, 10.0}, generating 6 predictors that are annotated as `fg-bin-2.5` to `fg-bin-10.0`, and `fg-wei-2.5` to `fg-wei-10.0` in Fig. 2.

### Path transitivity

Path transitivity ^50^ combines the shortest path between two nodes with the matching index to evaluate how many local detours are present at each step alongside the shortest path. To quantify the path transitivity between nodes *i* and *j*, the set of nodes belonging to their shortest path *π*_*i*→*j*_ = (*i, k, l*, …, *m, n, j*) is first identified. Then, the shared neighborhoods of all nodes from the shortest path are cumulated as follows: 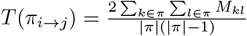 where *M*_*kl*_ is the matching index, and *π* denotes *π*_*i*→*j*_ for brevity. This measure is high whenever multiple other viable routes towards a target node are present at any point along the shortest path. We used binary and weighted undirected *W*, as well as shortest paths evaluated with *γ* = {0.125, 0.25, 0.5, 1.0, 2.5, 5.0} before computing path transitivity, resulting in a total of 7 predictors annotated as `pt-bin`, followed by `pt-wei-0.125` to `pt-wei-5.0` in Fig. 2.

### Search information

Search information ^63^ measures the amount of information required to move from one node to another through the shortest path that links them. Consider the shortest path between nodes *i* and *j, π*_*i*→*j*_ = (*i, k, l*, …, *m, n, j*). Then, the probability of following that path is *P* (*π*_*i*→*j*_) = *p*_*ik*_ × *p*_*kl*_ × … × *p*_*mn*_ × *p*_*nj*_ where 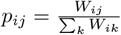. Finally, the information required to follow the path is *S*(*π*_*i*→*j*_) = − log_2_[*P* (*π*_*i*→*j*_)]. We used binary and weighted undirected *W*, as well as its shortest paths computed with *γ* = {0.125, 0.25, 0.5, 1.0, 2.5, 5.0}, to generate a total of 7 predictors annotated as `si-bin`, followed by `si-wei-0.125` to `si-wei-5.0` in Fig. 2.

### Navigability

Going through the shortest paths of a network requires global information, which could make this routing strategy implausible in circumstances where such information is not locally available. However, certain navigation strategies on networks could end up uncovering these shortest paths (or other near-optimal paths) without directly searching for them. For example, if nodes *i* and *j* are not directly connected, a good strategy to move from one to another might be to always follow connections toward nodes that are closer in Euclidean space to the desired target. This strategy has been shown to be efficient for navigating connectomes ^64^. We applied a similar navigation procedure to all pairs of nodes, first by converting directed connection weights *W* into a cost matrix *C* and then evaluating three different quantities for each pair of nodes: the binary length of a navigation path on the network (`nav-num-bin`), the cumulative weights of the same edges along the navigation path (`nav-num-wei`), and the total Euclidean distance traveled along the path (`nav-num-dist`).

### Module detection

To identify network modules, we used a hierarchical consensus algorithm based on the iterative Louvain algorithm ^150^ for maximization of modularity ^67^. We generated an ensemble of 10000 node partitions using either directed or undirected versions of modularity calculated on SC and FC matrices, respectively (`modularity_louvain_dir` and `modularity_louvain_und` in `bctpy`), varying the resolution parameter *γ* from 0.7 to 2.2. This yielded partitions that subdivided the network into 2 to 35 modules, where the upper boundary corresponds to roughly half of the network size beyond which one-node modules become necessary to further subdivide the network (although in practice some emerge earlier in the hierarchy). Then we computed the consensus matrix *C* whose values *c*_*ij*_ indicate the empirical probability that nodes *i* and *j* are assigned to the same module across the 10000 partitions. Finally, we used hierarchical clustering with Ward’s criterion ^151^ (`linkage` and `fcluster` from `scipy`) on the distance matrix *D* = 1 − *C*, thus converting high co-assignment probabilities into small distances. We generated varying amounts of modules by splitting the hierarchical dendrogram at different levels (`maxclust` criterion). The maximum number of modules for SC and FC was identified as the level in the hierarchy where a single (one-node) module emerges when further subdividing the network. We also identified null modules from ensembles of null matrices (SCCM, CM, phase-randomized FC, or shuffled FC, 1000 matrices each) using a reduced number of 1000 partitions per matrix before generating and clustering the consensus matrices with the same procedure as above. These null partitions were then used throughout the study to benchmark (1) the modularity of real networks, (2) the overlap between SC and FC, (3) the spatial compactness of modules, and (4) the modularity of inter-regional co-activation patterns.

### Identifying high-amplitude co-fluctuation events

To identify high-amplitude co-fluctuation events, we extracted regional fluorescence time series *f* (*t*) from both hemispheres, and then excluded brain regions that had at least one unsampled hemisphere (less than 25 imaged neurons) in at least one animal. This ensured that we kept the minimal subset of brain regions that were properly and symetrically sampled throughout the population, producing 120 total network nodes. Then, we z-scored each regional time series 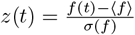, where the mean and standard deviation are calculated over time steps, and computed pairwise co-fluctuation matrices *C*(*t*) at each time step, whose values *C*_*ij*_(*t*) = *z*_*i*_(*t*)*z*_*j*_(*t*) reflect products of z-scored activity between regions *i* and *j*. Then we computed the root mean square (RMS) of each co-activation matrix to obtain a single number at each frame *t* whose magnitude reflects how the co-activated brain regions are, relative to their averages. To find a suitable threshold for event detection, we independently applied circular permutations to regional time series by splitting them at random time points and swapping the two segments, thus preserving their length, approximate frequency content, and numerical values, but changing their temporal order. We then calculated null co-activation matrices and RMS values from the shuffled time series across the population of 22 larvae, repeating this process 500 times to yield a large distribution of null RMS values. We used the 95th percentile of this null distribution to threshold the empirical RMS values, resulting in 2382 frames with significant co-fluctuation amplitudes. Then, we applied the Benjamini-Hochberg procedure ^152^ to control for false discovery at a rate *α* = 0.05, which further reduced the number of significantly co-activated frames to 1755. We finally merged all successive frames into single events, yielding a total of 243 high-amplitude co-fluctuation events. We clustered these events using t-SNE followed by HDBSCAN ^70^, which identified 3 clusters from the 2D embedding across multiple runs (Supp. Fig. S13). A few events were excluded by the density-based clustering algorithm, but were reassigned to the existing clusters based on which cluster centroid they were maximally correlated with.

### Rendering raw fluorescence in brain atlas

To visualize the raw whole-brain fluorescence associated with co-activation event clusters, we first detrended each motioncorrected calcium imaging plane by subtracting a piecewise linear regression at each pixel along the temporal axis, thereby subtracting the pixel’s temporal mean and removing photobleaching effects. This procedure greatly improved visual contrast within the raw data without amplifying noise. Then we averaged all frames belonging to a certain co-activation cluster and generated 3D temporal averages of all 21 imaging planes for each animal, which we then projected into mapZebrain using the `antsApplyTransforms` function. Once transformed, we averaged the stacks to obtain smooth snapshots of brain activity within the brain atlas. We observed that the warping and 3D interpolation used in the transforms reduced the contrast between active and inactive cells within individual stacks. Therefore, we squared the intensity values to improve contrast before projecting them in Fig. 4 and rendering them in 3D in Supp. Video 3.

### Identification of motor and visual cells

To find visual- and motor-correlated cells, we used a common linear regression approach for calcium imaging data ^73^. We generated regressors representing hypothetical calcium responses by convolving an exponential kernel that models slow calcium dynamics (*τ* = 3.5s) with vectors representing the occurrence or amplitude of events of interest. To retrieve dark- and light-responsive cells, we generated binary vectors of a length similar to the calcium time series, with ones assigned to frames where a stimulus of interest was being presented, and zeros otherwise, before convolving them with the exponential kernel. We correlated these regressors with fluorescence time series during the corresponding stimulus period, and then temporally offset the regressors to generate a null distribution of correlations from spontaneous neuronal activity in each animal, where no stimuli were present. We used the 99th percentile of this null correlation distribution as a threshold for significantly dark or light-responsive cells. For motor and anti-motor neurons, we first generated downsampled swimming event vectors of similar size to the calcium time series, with values indicating the maximal amplitude of swimming events occurring at each calcium imaging frame. We then convolved this vector with the exponential kernel, yielding hypothetical calcium responses of varying amplitudes, under the assumption that more vigorous swimming events are associated with larger calcium responses (which is validated empirically). Because swimming occurred throughout the experiment for most animals, we could not temporally offset the regressors to find periods of complete motor quiescence where null correlations could be estimated. Instead, we randomly shuffled the swim event vectors 250 times before convolving them with the exponential kernel, and then correlated these shuffled regressors to real fluorescence time series, thus yielding an alternative null distribution of correlations for each animal. Neurons that exceeded the 99th percentile of this distribution were considered significantly motor-correlated, while those that were below the 1st percentile were considered significantly motor-anticorrelated, or anti-motor cells. We observed more cells that correlated significantly with dark stimuli than cells that correlated with swimming; however, this difference could reflect the different statistical methods from which null distributions were generated for each functional category. As such, no statistical comparisons are made between cells from different functional categories.

After finding the significantly correlated neurons, we applied a second statistical filtering procedure established in a previous whole-brain imaging study, a spatial *P* -value analysis ^75^. We used this method to exclude cells that were located at coordinates of the brain where few other cells from the same functional category were found nearby in other animals. This ensured that only the regions that were the most reliably responsive regions to a stimulus or motor event were preserved in the final consensus spatial densities of functionally specialized cells. Briefly, for each neuron identified previously using regression, we found its nearest neighbors (NN) from the same functional category in brain atlas coordinates within the other animals. This generated a vector of nearest distances [*dr*_1_, *dr*_2_, …, *dr*_*m*_] where *m* = 22 larvae for stimuli and *m* = 18 for motor activity. We computed the average of this vector to yield a single scalar representing the average proximity of this neuron to its functional neighbors in other individuals. Then we generated 1000 null distance vectors by randomly sampling cell coordinates from the total number of neurons within the other animals, and computing distances with null nearest neighbors. We then averaged each null distance vector and compiled them into a single null distribution of average NN distances. We repeated this procedure for all functionally correlated neurons and preserved only those whose empirical average *dr* was lower than the 2.5^th^ percentile of its associated null distribution (thus *P <* 0.025).

### Quantifying regional functional diversity

We estimated the regional diversity in functional cell types by first counting the number of cells *n*_*i*_ from each functional category *i* present within a given brain region, measured separately for each animal. Then we converted these values into proportions such that 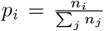, with the denominator set to 1 if there were no cells from the four categories within the region. This produced proportion vectors *p* = [*n*_1_, *n*_2_, *n*_3_, *n*_4_] for each brain region, representing motor +, motor −, dark and light cells, respectively, from which we computed the regional Shannon entropy *H* = ∑_*i*_ *p*_*i*_ log(*p*_*i*_). We then derived a regional diversity index by averaging these entropy values across animals for each brain region and dividing this measure by its standard deviation, such that 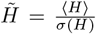. This operation penalized regions whose functional diversities were uneven across animals, which corresponded mostly to regions with very low cell counts. These regions tended to reach high diversity scores by chance in some animals, increasing the regional population average but also its standard deviation.

### Computing network gradients

To compute FC and SC network gradients, we used the diffusion map algorithm ^92^, which exploits the properties of random diffusion on graphs to separate data points (network nodes) based on their distances in a diffusion space. Briefly, the method consists of generating a Markov transition probability matrix *M* from a connectivity matrix *W*, and then using its eigenvectors *ϕ*_*i*_ (weighted by their corresponding eigenvalues 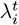, where the exponent *t* is the diffusion time) to define a new coordinate system where data points cluster naturally based on their topological proximity within the graph. In the case of brain networks, two points that are close to each other within this space can be interpreted as two brain regions *i* and *j* that are strongly connected or that share many connectivity neighbors, such that random walkers are likely to move from brain region *i* to brain region *j* and vice versa after a certain diffusion time *t*. Each eigenvector (gradient) of *P* can also be interpreted individually as indicating a major axis of diffusion along the brain network. If a gradient is sufficiently smooth, that is, if there are no abrupt jumps in its numerical values, then random diffusion occurs smoothly along the gradient on the network, from positive to negative values and vice versa. Thus, diffusion maps allow 1) the embedding of brain networks to visualize the relationships between connected neural elements in diffusion space and 2) the visualization of diffusion axes (putative processing streams) along the brain surface or volume.

We used the Python implementation of the BrainSpace toolbox ^153^ to compute gradients from both FC and undirected SC. The *α* parameter, which controls the degree-normalization of the connectivity matrix in the calculation of the diffusion operator, was set to the standard value *α* = 0.5 where the operator *M* approximates the Fokker-Planck diffusion. The code implementation uses the following normalization of the eigenvalues, 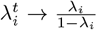, which provides a trade-off between all possible diffusion times *t* and thus eliminates the additional choice of parameter *t*^154^. We computed all gradients for both FC and SC, but restricted our analyses to the first 50 gradients in Fig. 6**a**-**d**. To compare gradients, we used an optimized linear sum assignment to establish a mapping between structural and functional gradients. Briefly, we computed the correlation matrix of functional and structural gradients and then optimize the sum max ∑_*i,j*_ *C*_*ij*_*X*_*ij*_, where *C* is the correlation matrix between SC and FC gradients, and *X* is a boolean matrix where *X*_*ij*_ = 1 if gradient *i* is assigned to gradient *j*. In other words, this optimization finds a global optimum of values on the diagonal of the matrix by reordering the rows and columns of *C*. We averaged the optimized values on the matrix diagonal to evaluate the global similarity between gradients, and we assessed the significance of this value through comparisons with null diagonal values obtained by finding an optimal mapping between SCCM and FC gradients. We also compared optimally mapped modes one by one, evaluating their statistical significance using both null connectivity gradients derived from SCCM as well as SA-preserving spatial permutations.

### Sensorimotor index

To compare the first functional gradient with the sensorimotor functions of brain regions, we first derived a “sensorimotor index” from the spatial densities of dark- and motor-correlated neurons. We used the consensus densities, obtained by combining correlated neurons from all larvae and applying a spatial *P* -value test, to annotate the proportion of cells *ρ* within each category across brain regions (see Supp. Fig. S19**a**). Then, we calculated the index *SI* by simply subtracting the proportions of dark and motor cells to each other, *SI* = *ρ*^dark^ − *ρ*^motor^. This yielded a regional vector of proportions in which positive scores are associated with regions with more dark-responsive neurons, and negative scores are associated with regions that contain more motor-correlated cells. We replicated these results in a high-resolution parcellation. First, we estimated a number of subdivisions for each region in order to split them into roughly 1000 nodes of similar size within the left hemisphere. Then, we divided each region mask using K-means clustering with the previously determined number of clusters, and then used the cluster centroids as node coordinates within our new parcellation. We mirrored nodes to the right hemisphere and assigned individual neurons to these nodes based on their nearest neighboring node, thus fractioning the brain into a Voronoi diagram. We computed nodal time series, averaged across individual cells and across both hemispheres, and then calculated individual FC matrices, which we averaged across individuals. We excluded all nodes that were unsampled in at least one hemisphere for at least one animal. Similarly, we computed a high-resolution sensorimotor index based on the nodal proportions of dark and motor cells. We used the first 10 minutes of recording for the spontaneous FC matrices, and the full-length time series (15 minutes) for the visually stimulated FC.

### Quantifying HCR markers

We downloaded 290 HCR gene expression markers ^95^ from mapZebrain ^32^, from which we estimated regional levels of gene expression. Individual fluorescent transcripts could not be resolved, as each marker was imaged at insufficient spatial resolution and averaged across multiple individuals. Thus, we used raw regional fluorescence levels to estimate gene expression. We compiled regional intensity levels by averaging the intensity of HCR markers within each brain region’s binary mask, and we normalized these intensity values by the level of background intensity within each marker, measured from a 50 × 50 pixels patch outside of the brain, roughly 40 microns below the upper delimitation of the atlas volume. We z-scored the final expression levels across genes for each brain region to account for spatial heterogeneities in the signals, for instance, to compensate for ventral regions, where intensity levels are reduced due to light scattering.

### Simulated annealing optimization

We evaluated the similarity of gene expression levels across brain regions by computing pairwise correlations of the rows of the gene expression matrix, yielding a 65 × 65 co-expression matrix that was poorly correlated with FC (Fig. 7**c**, left, Spearman’s *r* = 0.33). We used a simulated annealing algorithm ^155^ to truncate the gene expression matrix, keeping only a small subset of genes (columns) from which to evaluate the co-expression matrix, which greatly improved upon the previous correlation (Fig. 7**c**, right, Spearman’s *r* = 0.71). Briefly, in simulated annealing, the solution space is explored by moving from one potential solution to another by randomly swapping one element at a time, with the acceptance of new solutions determined by a probability function that depends on a temperature parameter *T*. This temperature decreases over time, reducing the likelihood of accepting worse solutions as the algorithm progresses. This provides a balance between exploration in the high temperature regime, followed by exploitation at low temperatures. Each optimization started with a random set of *M* genes, where *M < N*. Then, we evaluated the co-expression *C*^opt^ of their regional expression levels *G* - a 65 × *M* truncated version of the full gene expression matrix in Fig. 7**b** - by correlating the rows with each other, yielding a 65 × 65 co-expression matrix. Then we evaluated an objective function *F* (*FC, C*^opt^) between *FC* and *C*^opt^, which produced a similarity score *υ*_*i*_, where *i* represents the current iteration in the optimization. We used normalized mutual information (NMI) as an objective function, which better quantified subtle differences between the two matrices than linear correlations. In the following iteration, we randomly swapped a single gene from the optimized set to a new one, computed a new co-expression matrix, and re-evaluated the objective function *F* to yield a new score. If *υ*_*i*_ *> υ*_*i*−1_, the new solution was always accepted. Otherwise, the solution was accepted with a probability 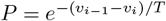, which is unlikely if the current score *v*_*i*_ is significantly lower than the previous iteration *v*_*i*−1_, but relatively likely for slightly worse solutions and for higher temperatures. We repeated this procedure for 30000 iterations, with an initial temperature of *T* = 2.5 that decreased gradually over iterations following *t*_*i*_ = *t*_*i*−1_ × 0.9995. We considered the final set of *M* genes after all iterations were completed as an optimized gene set, and we compiled thousands of such sets by conducting optimizations at varying *M* sizes. We selected the optimal *M* = 12 value by evaluating the elbow of the NMI curve in Fig. 7**d**, which is the point along the curve that is most distant from a line that connects the two extremities of the curve. Across multiple runs at *M* = 12, some genes were selected very often within optimized gene sets. We identified these genes by running a similar number of optimizations on spatially shuffled gene matrices and then evaluating the maximal number of times that a shuffled gene was selected. We kept only the genes whose selection probabilities were higher than this stringent chance-level threshold.

### Statistics

The statistical significance of multiple results throughout this study was assessed either by comparisons with null network models (SCCM or CM), or with spatially permuted node properties, such as activity or gene expression. For null network models, we typically benchmarked empirical network properties such as network modularity with similar properties calculated on 1000 null network matrices. In some cases, the empirical values exceeded all values from the null distribution, which we denoted as *P <* 0.001. Otherwise, *P* -values corresponded to the percentiles of empirical values relative to the null distribution. For spatial permutations, we used the BrainSMASH method ^71^, which reorganizes values within a 1-dimensional node property vector while approximately preserving their spatial autocorrelation based on the spatial distances between nodes. We applied this method to shuffle regional activity patterns, the sensorimotor index, or gene expression levels, and constrained shufflings with the inter-regional Euclidean distances *D*. Empirical *P* -values were reported relative to null distributions as described above.

## Data availability

The processed data such as cell coordinates and fluorescence time series will be made publicly available. Raw calcium imaging movies and behavioral data (*>* 1 TB) are available upon reasonable request to the authors.

## Code availability

Data analysis was performed using Python. The code required to replicate the analyses and figures is available on GitHub (https://github.com/pdesrosiers/Legare-2024-zebrafish-brain-networks). The repository is subdivided into several notebooks that perform analyses related to each figure. Figures were generated with Matplotlib and completed using Inkscape.

## Acknowledgments

We thank members of the PDK Lab and the Dynamica Research Lab for helpful discussions that contributed to this work; the staff from the Laboratoire aquatique de Recherche en sciences environnementales et médicales of Université Laval and the animal facility at CERVO for zebrafish husbandry, and Sandra Mignault for technical assistance; Minoru Koyama, Vincent Breton-Provencher and Alexandre Cléroux-Cuillerier for helpful comments on the manuscript; Ismael Djerourou, Cynthia M. Solek, and Youngseung Yoo for helping with the design and printing of the imaging chamber during the 2023 Frontiers in Neurophotonics Summer School. A.L. was supported by PhD scholarships from the Natural Sciences and Engineering Research Council of Canada (NSERC), Fonds de recherche du Québec – Nature et technologies (FRQNT), and Unifying Neuroscience and Artificial Intelligence - Québec (UNIQUE). S.P. was supported by a scholarship from FRQNT. This work was funded by NSERC (P.D. RGPIN-2019-06887 and RGPIN-2024-06492, P.D.K RGPIN-2023-05980), the Sentinel North program of Université Laval (Canada First Research Excellence Fund), the Northern Contaminants Program of Canada, and the Next Generation Networks for Neuroscience (Neuronex)/FRQS (295823). We are grateful to Calcul Québec and the Digital Research Alliance of Canada for their technical support and computing infrastructure. The Tg(Elavl3:H2B-GCaMP6s) zebrafish line was developed by Misha Ahrens (Janelia, USA) and was obtained from Edward Ruthazer’s lab (Montreal Neurological Institute, McGill, Canada).

## Author contributions

An.L., M.L., P.D. and P.D.K. conceived the study; An.L. and V.B. developed the imaging protocols and behavioral monitoring system; An.L. and S.P. built the visual stimulation system; An.L. performed all the experiments and analyzed the data; Ar. L. helped with data processing and analysis; An.L. wrote the initial draft; An.L., M.L., P.D and P.D.K. edited the final manuscript.

## Supplementary material

**Figure S1:**
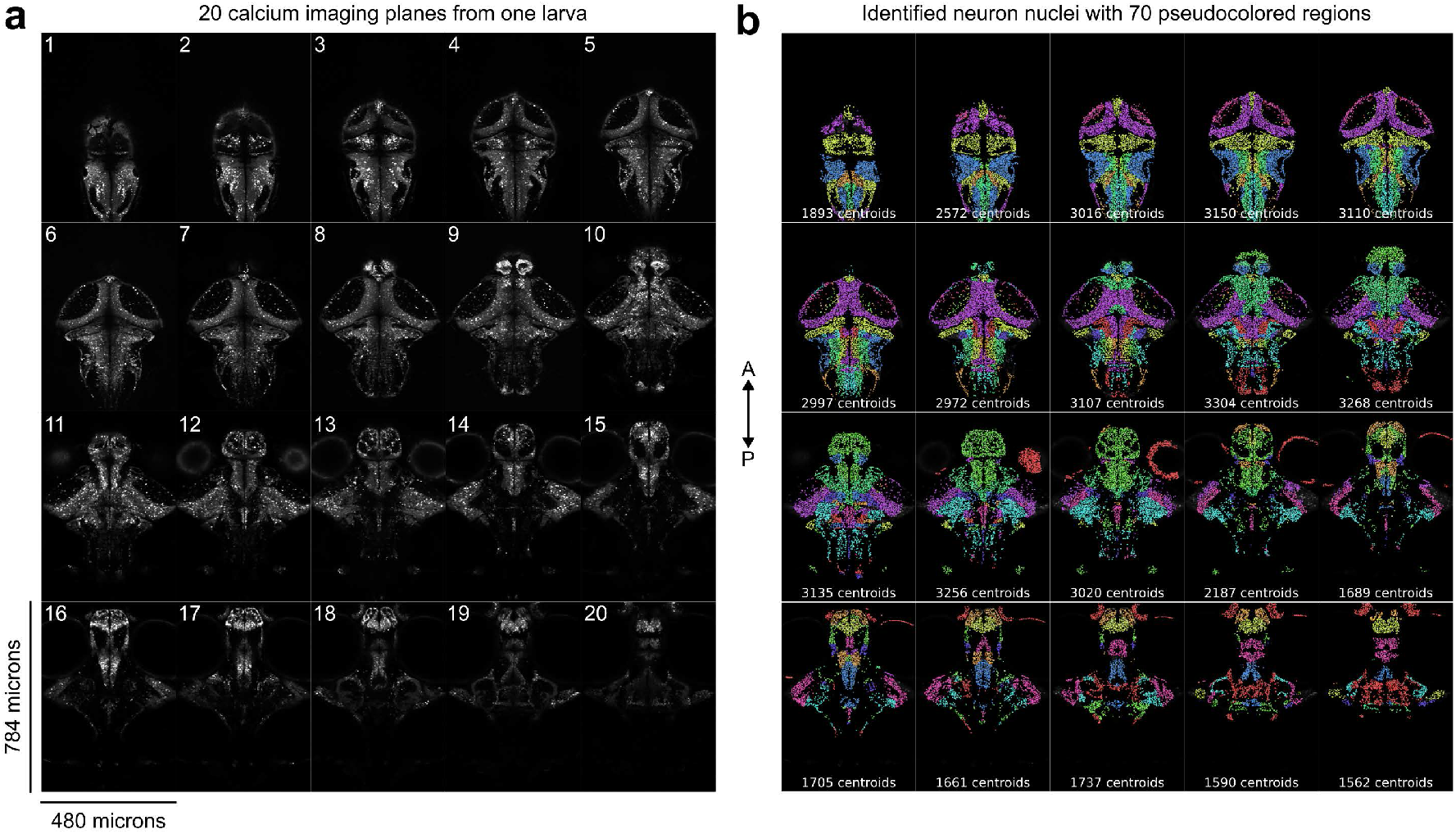
(**a**) 20 out of 21 mean intensity projections of calcium imaging planes from one example larva, after motion correction has been applied; planes are spaced roughly 12 microns apart in depth. (**b**) Identified nuclei on each corresponding plane from the previous panel, pseudocolored by brain region; the number of nuclei detected per plane is identified on each slice. Note that the autofluorescence of the eyes (from plane 11 onward) is incorrectly detected as nuclei; these artifacts are excluded from analysis, along with four other regions (Methods). A, anterior; P, posterior.

**Figure S2:**
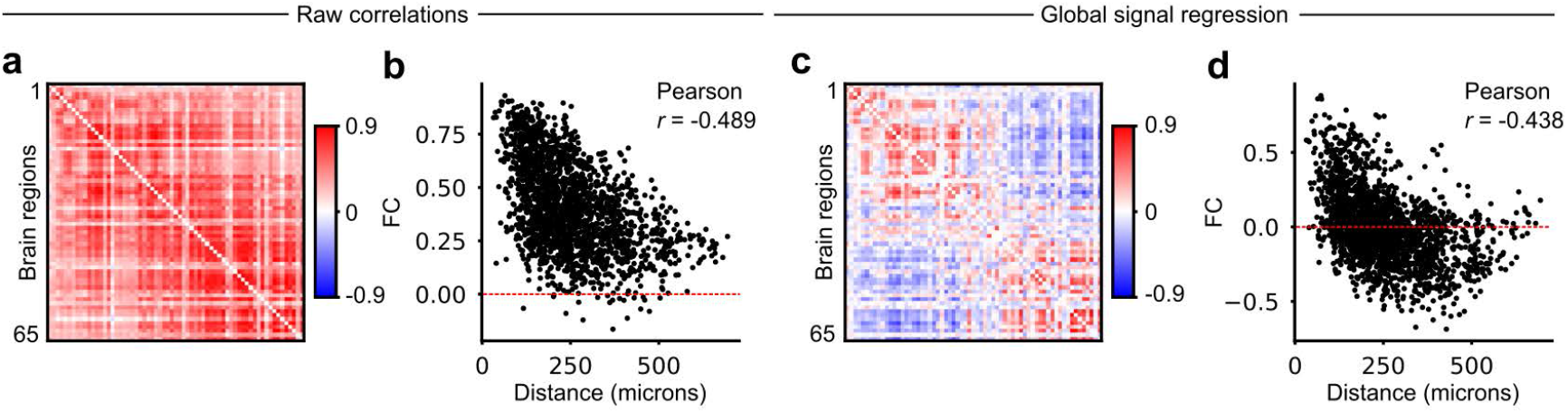
Distance dependance of functional connectivity. (**a**) Group-averaged FC matrix (*n* = 22 larvae), raw correlations (similar as Fig. 1**e**). (**b**) FC against spatial distance between brain regions; each point corresponds to one FC edge; a red dotted line delimits positive and negative correlations. (**c**) Group-averaged FC matrix (*n* = 22 larvae) after applying a global signal regression on regional signals of each larva. Note the emergence of negative correlations (in blue), especially between anterior and posterior brain regions (bottom left/top right matrix corners). (**d**) FC against spatial distance after applying global signal regression.

**Figure S3:**
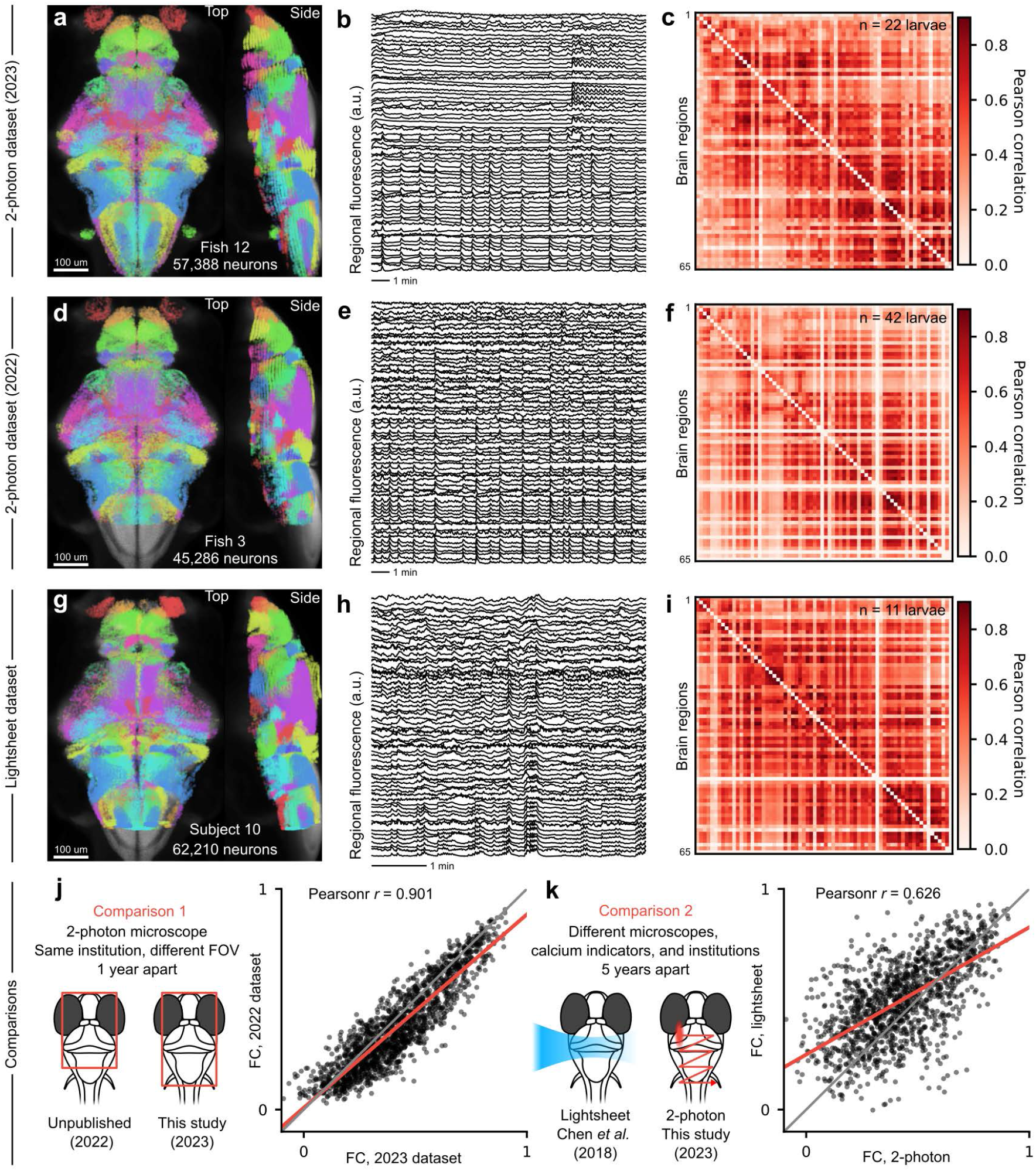
Group-averaged functional connectivity from 3 datasets (top, two-photon microscopy from this study; middle, unpublished, two-photon microscopy; lightsheet microscopy from Chen *et al*. (2019)). First column (**a, d, g**): neuron centroids from one representative fish in each dataset, projected in mapZebrain. Second column (**b, e, h**): regional calcium time series from the same fish; the number of imaged regions varies across fish and datasets. Third column (**c, f, i**): group-average functional connectivity from each dataset, using the 65 region parcellation from this study; some regions are unsampled in the second and third dataset, denoted by white rows. (**j**) Correlation between group-averaged FC of datasets 1 and 2, encompassing 58 most rostral regions. (**k**) Correlation between group-averaged FC of datasets 1 and 3, encompassing 53 most rostral regions. Red lines represent linear fits, and gray lines represent the unit slope where correlations are equal between datasets.

**Figure S4:**
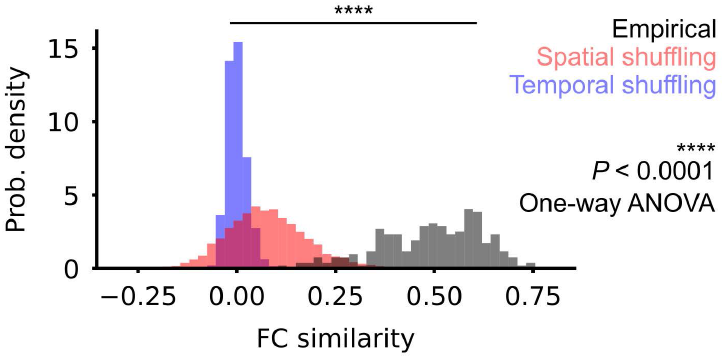
Distributions of inter-larva FC similarity (Pearson correlation of FC matrices). Gray distribution, empirical similarity scores between larvae. Red distribution, similarity scores after spatial reshuffling of FC matrices, which preserves numerical values (500 permutations, see Methods). Blue distribution, similarity scores after cyclic permutations of regional time series, leading to much lower correlation values and inter-larval similarities.

**Figure S5:**
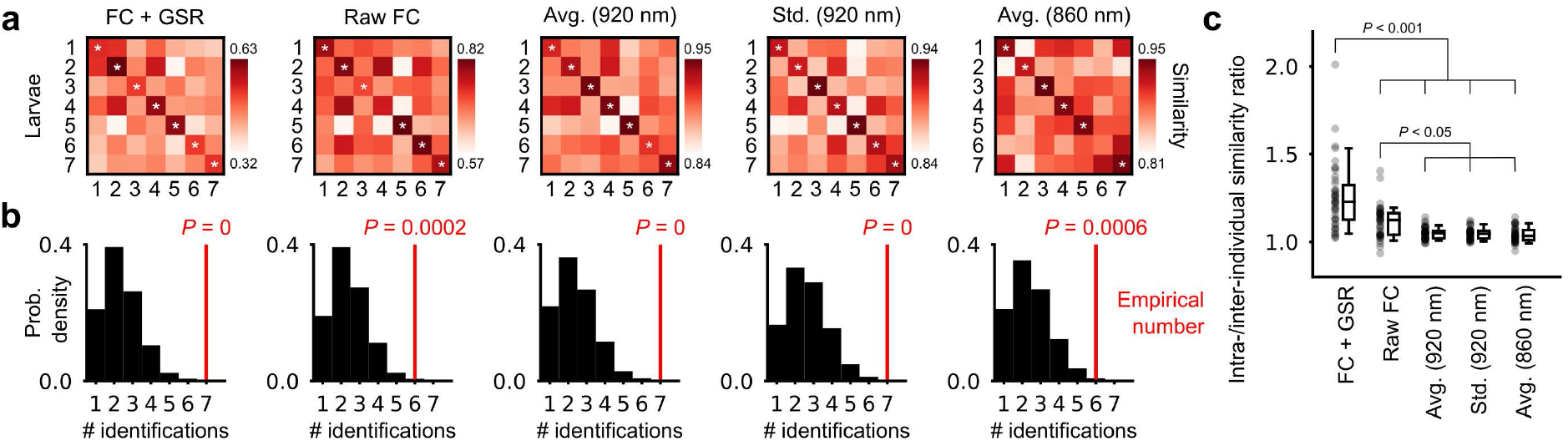
Statistical validations of identification success rate using FC fingerprinting and benchmarking of different fingerprinting methods. (**a**) Inter-larva FC similarity matrices between 6 and 7 dpf for five different fingerprinting methods; white asterisks denote successful identifications corresponding to row-wise maxima located on the matrix diagonal; the first two methods correspond to FC-based fingerprinting (with and without GSR), and the three subsequent methods use regional fluorescence intensity vectors (average or temporal standard deviation) to fingerprint larvae; a 920 nm excitation wavelength uses calcium-dependent fluorescence (methods 3 and 4), while 860 nm is near the isosbestic point of the sensor and reflects baseline fluorescence levels, independent of neuronal activity (method 5). (**b**) Null distributions of identification successes for each of the fingerprinting methods in the panel above; distributions are obtained using the previous FC matrices or fluorescence vectors with identities shuffled in one of the two datasets; empirical *P*-value indicated on the graph. (**c**) Distributions of intra-vs inter-individuality ratios for each fingerprinting methods; values correspond to matrix diagonal elements in **a**, divided by each off-diagonal value along the same row (comparisons with other animals), thus capturing the relative magnitude of intra-individual similarity; *P* -values for one-way ANOVA followed by Tukey’s post hoc comparisons are indicated on the graph. GSR, global signal regression; Avg., average; Std., standard deviation.

**Figure S6:**
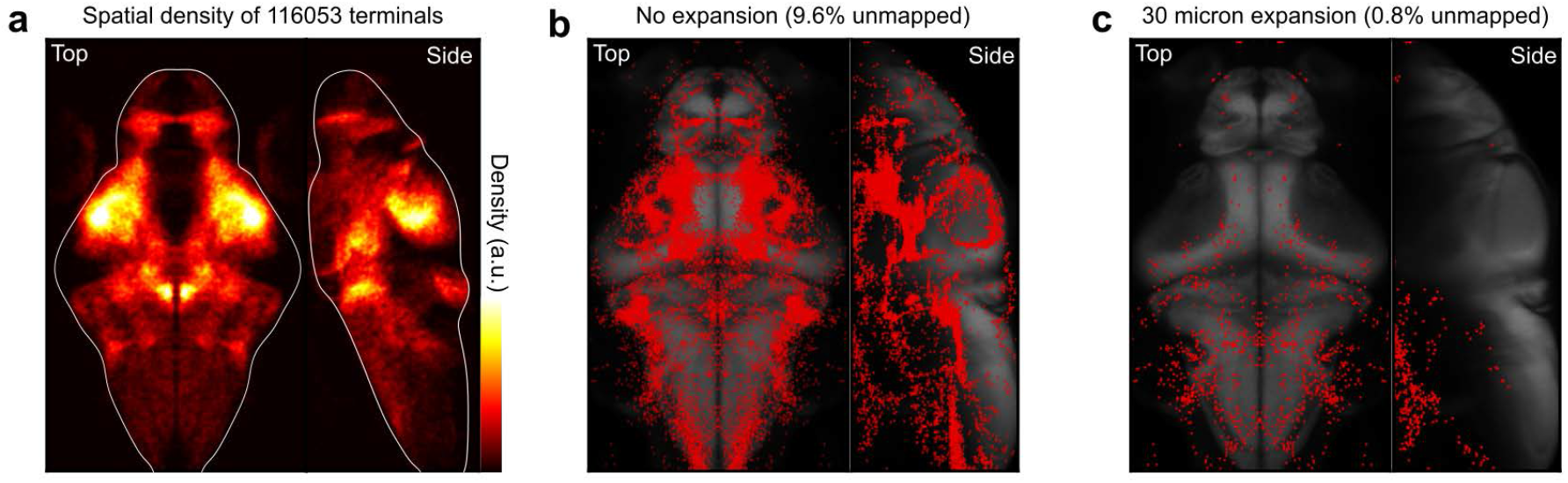
Visualization of neurite terminals used in SC reconstruction. (**a**) Heatmap of 116053 neurite terminals from 4327 individually reconstructed cells in mapZebrain, mirrored across hemispheres; the colormap reflects arbitrary spatial density values obtained using a gaussian kernel applied on terminal coordinates; white outlines roughly indicate brain boundaries. (**b**) Centroids of terminals that are not part of any brain region ROI (9.6% of terminals). (**c**) Centroids of terminals that are not part of any brain region ROI following a 30 micron expansion of regional masks (0.8% of terminals). The remaining unmapped terminals after regional expansion are mostly located in ventral, neurite-dense regions.

**Figure S7:**
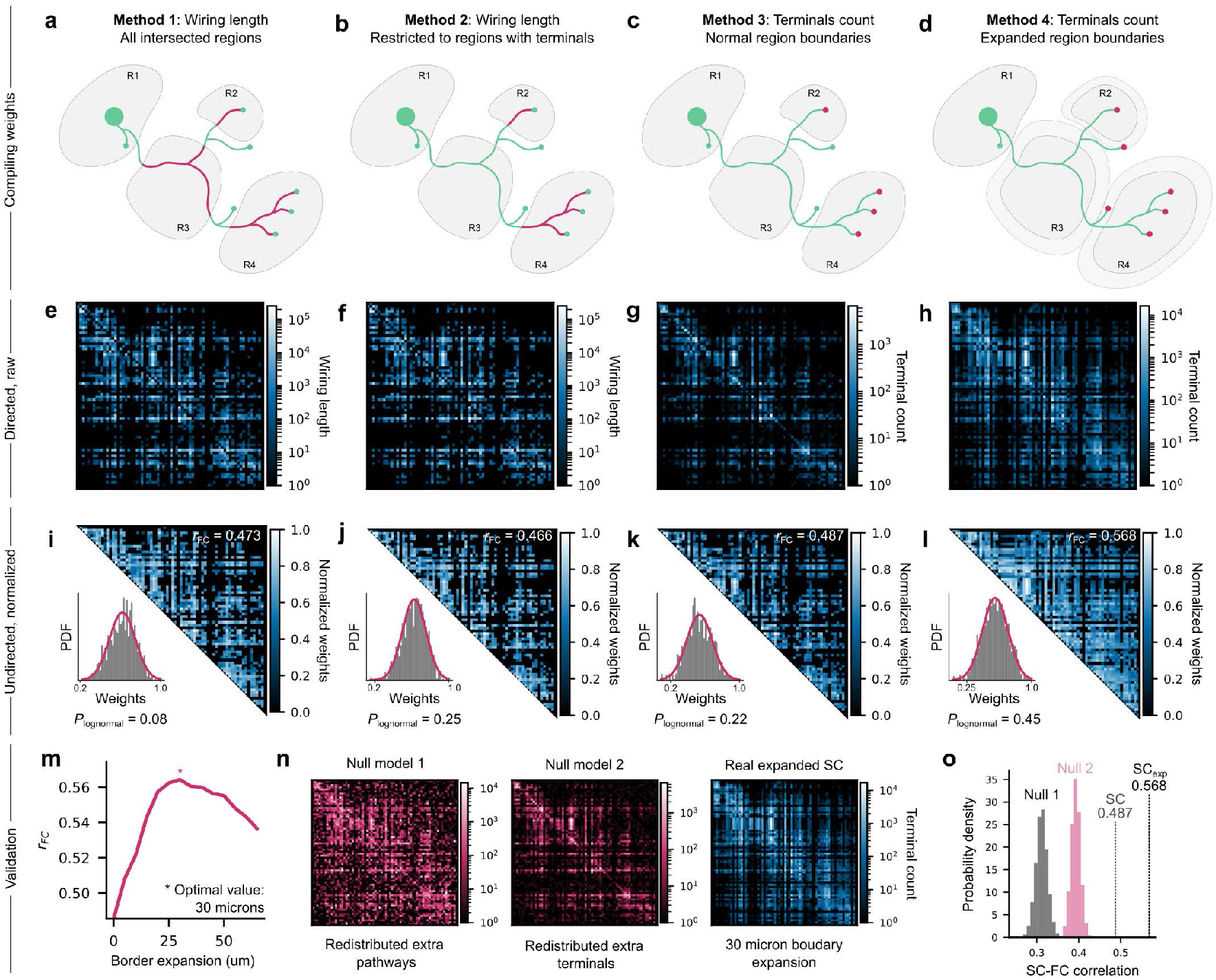
Generating a mesoscopic structural connectivity matrix from single-cell neuron reconstructions. (**a**-**d**) Graphical depiction of four connection weighing schemes. In green, a single neuron whose soma (large green circle) is located in region R1, sends projections through region R3, then terminates (small green and magenta circles) in regions R2 and R4. Magenta denotes which segment of processes and terminals are used in each scheme to compile connections weights. For precise descriptions of each weighing scheme, see Methods. (**e**-**h**) Raw directed connectivity matrices obtained from each of the four methods after compilation of the entire single-neuron dataset. Matrices are ordered correspondingly with the methods above. (**i**-**l**) Undirected matrices generated by symmetrizing, then log-normalizing the raw directed matrices obtained previously (directly above). *r*_*F C*_ indicates the Pearson correlation of each matrix with group-average FC. Log-normal weight distributions in the lower-left corner are evaluated using a Kolmogorov-Smirnov test, where higher *P* -values indicate a better fit. (**m**) *r*_*F C*_ values obtained for matrices generated using Method 4, for varying amounts of regional boundary expansion. An asterisk indicates the maximal correspondence between structure and function used throughout this study, at a 30 micron expansion. (**n**) Example matrices from two null models of adding extra terminals to count additional connections and replicate the increased density of the expanded boundary method (see Methods). (**o**) Distributions of SC-FC correlations for 1000 matrices generated with both null models; both correlation distributions are lower than the original unexpanded SC matrix (*P <* 0.001).

**Figure S8:**
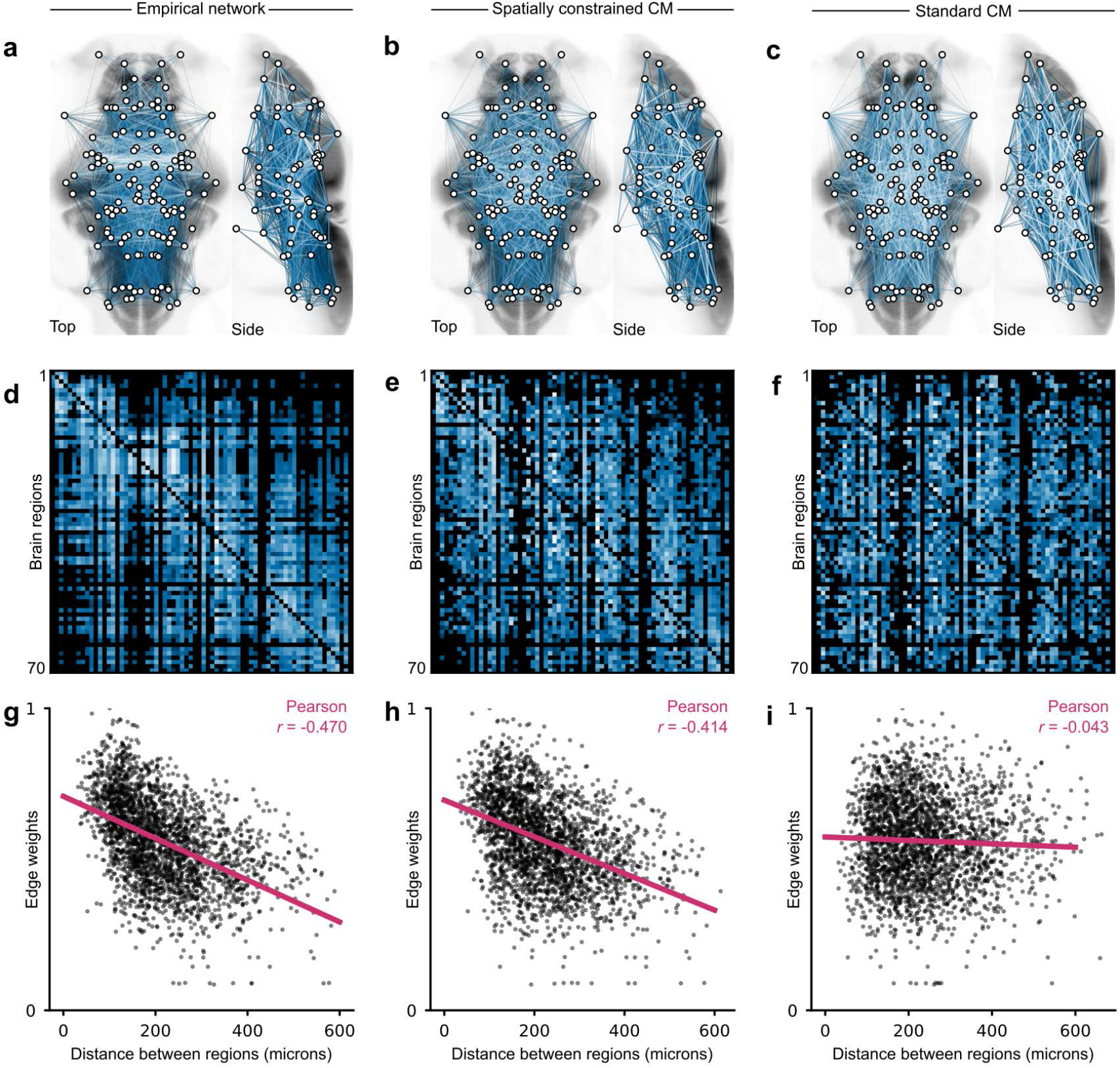
Comparison of two null models of SC with empirical connectivity. (**a**-**c**) Undirected network representations overlayed on brain anatomy for empirical SC, a spatially constrained configuration model (SCCM) and a standard configuration model (CM); bottom quartile edges are not displayed for visibility, and lighter colors indicate higher edge weights. (**d**-**f**) Example directed connectivity matrices for the three networks displayed above. (**g**-**i**) Distance-weight relationships for the networks displayed above; SCCM approximately preserves this negative relationship, while CM abolishes it. Note that SCCM and CM are shuffled versions of the real network, thus one representative example of a shuffling is shown in each case, and correlations in the bottom row may differ slightly across the matrix ensemble.

**Figure S9:**
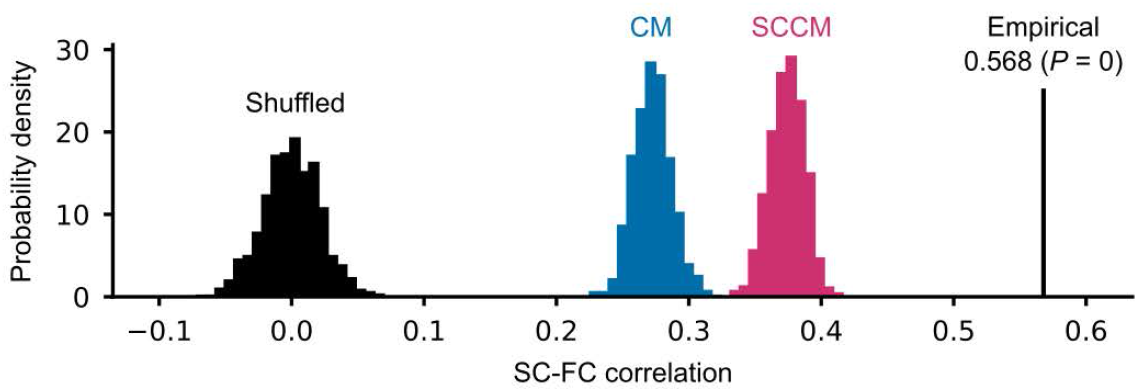
Comparison of the empirical SC-FC correlation with three different null models (spatially constrained configuration model (SCCM), configuration model (CM) and randomly shuffled matrices); the empirical correlation exceeds all distributions (*P <* 0.001, 1000 matrices per distribution).

**Figure S10:**
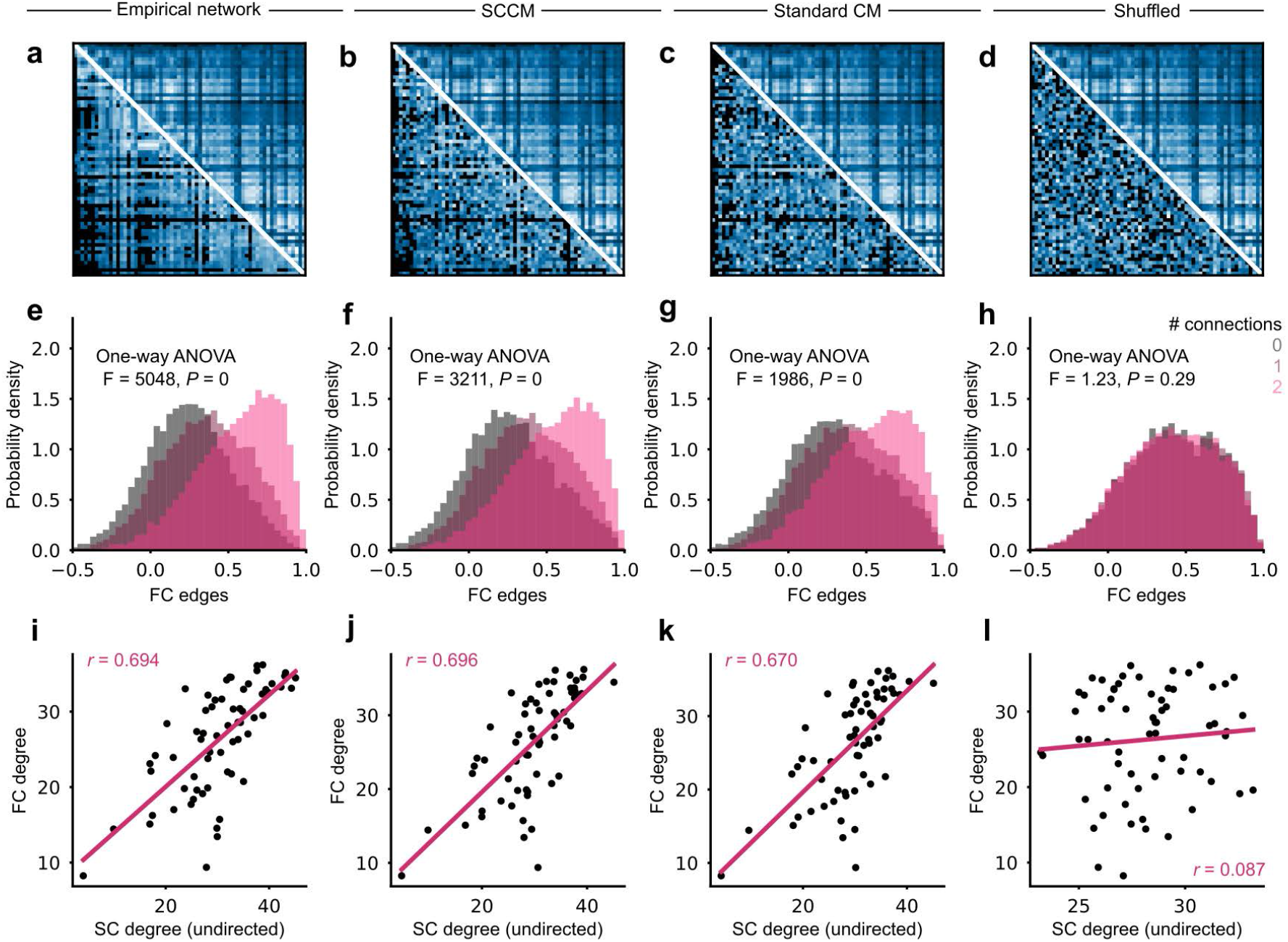
Relationship between FC and the number of direct structural pathways connecting two regions for empirical networks and various null models. (**a**-**d**) Comparison of FC (top right) with undirected SC (bottom left) for empirical networks, a spatially constrained configuration model (SCCM) of SC, a standard configuration model (CM) of SC, and a randomly shuffled SC matrix. (**e**-**h**) Distributions of compiled FC weights between regions that exchange bidirectional structural connections, a single unidirectional structural connection, or that are structurally unconnected; the relationship between directionality and functional weight is preserved in both degree-preserving null models (shifts in distributions, one-way ANOVA, *P* = 0 in both cases), but not for the randomly shuffled SC matrix (one-way ANOVA, *P* = 0.29); note however the degradation in the *F* -statistic. (**i**-**l**) SC-FC degree correlations (Pearson’s *r*) for all network cases; both SCCM and CM approximately preserve weighted degrees by design, whereas shuffled networks lose this correlation. These results suggest that degrees are a driving factor of the shift between FC weight distributions.

**Figure S11:**
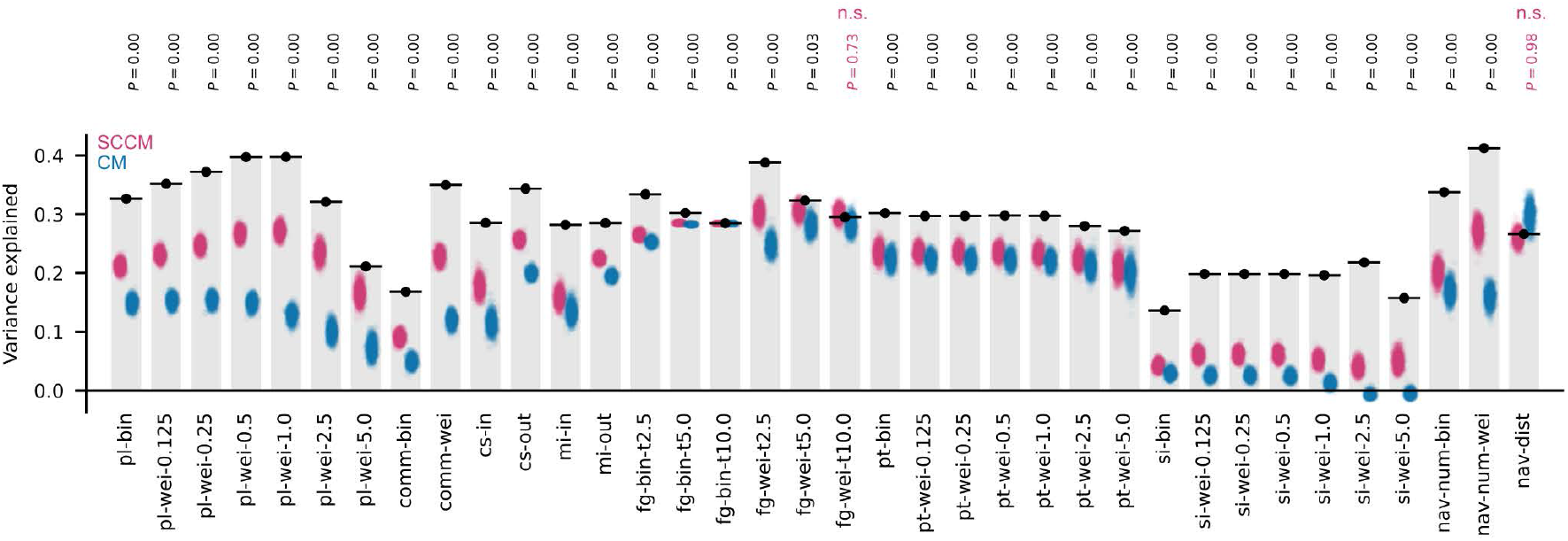
FC variance explained by different network properties derived from empirical SC (black lines/dots), and from matrix ensembles generated using a spatially constrained configuration model (SCCM) and a standard configuration model (CM) (1000 matrices each, pink and blue point distributions respectively); *P* -values are indicated above each structural property; only weighted flow graph with *t* = 10 (fg-wei-t10.0) and Euclidean distance traveled by navigation (nav-dist) fail to significantly exceed null distributions.

**Figure S12:**
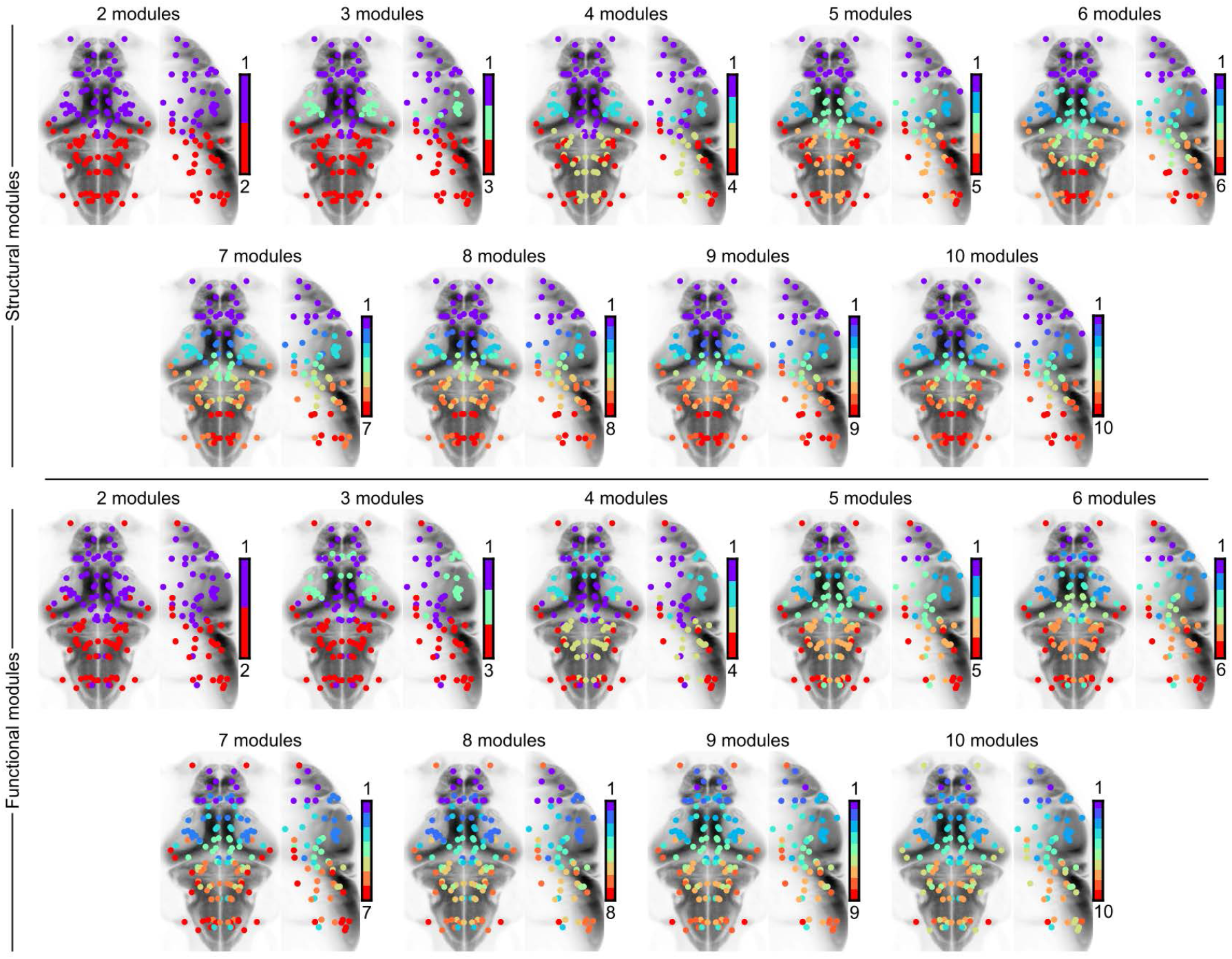
Hierarchical modules of SC (top) and FC (bottom), ranging from 2 to 10 subdivisions of the network. Note that singlets (modules of one node) emerge beyond 7 modules for FC, while they emerge beyond 10 modules for SC (not shown); we use the emergence of singlets as the maximal hierarchical subdivision of each network.

**Figure S13:**
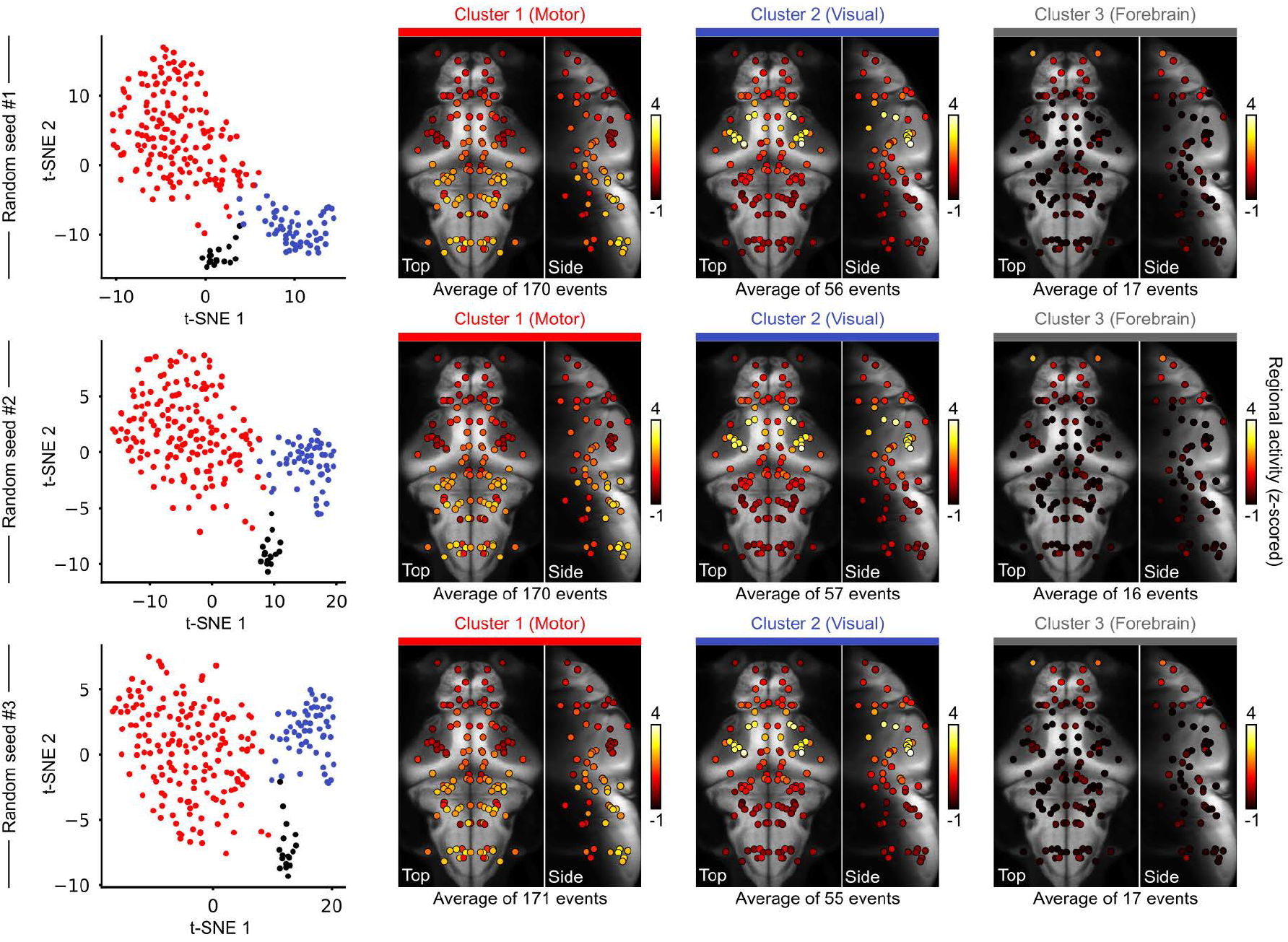
Three instances of randomly initialized clustering runs of high-amplitude regional co-fluctuation events. Note that the recovered embeddings and clusters are very similar across runs. Clusters 1 and 2 are displayed in Fig. 4, while the third forebrain cluster intercepts very few high-amplitude events and is not analyzed further. These forebrain events are better recovered in subsequent analyses using independent component analysis in the main text, or using clustering of low-amplitude brain activity configurations in Supp. Fig. S15.

**Figure S14:**
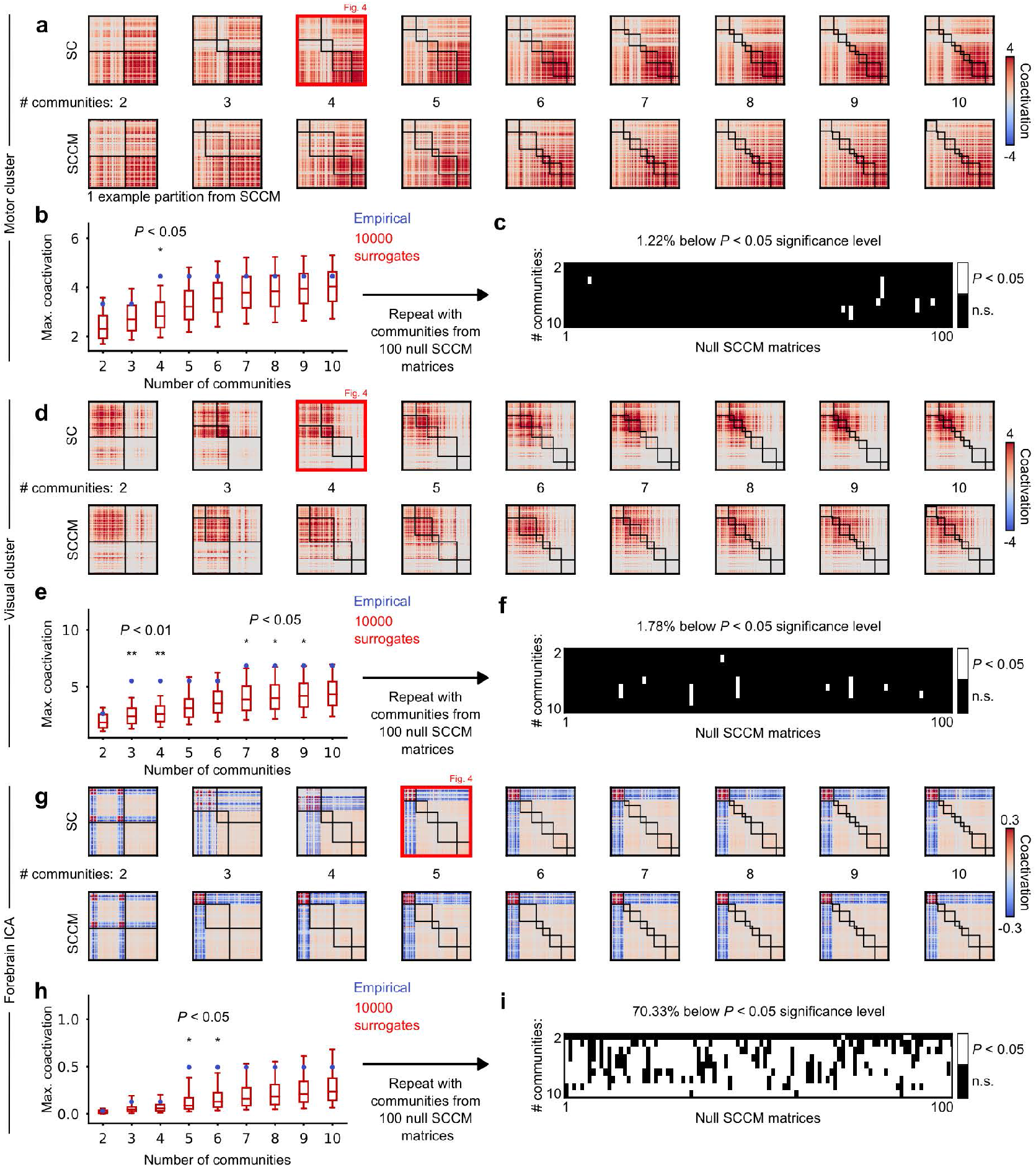
Statistical validation of the modularity analysis of regional co-fluctuation events. (**a-d-g**) Top row, average functional co-fluctuation matrices of motor, visual and forebrain activity configurations, sorted by increasing subdivisions of structural modules; bottom row, the same matrices sorted using null modules inferred from null SC matrices (spatially constrained configuration model, SCCM); one representative set of null modules is displayed for each activity cluster. (**b**-**e**-**h**) Red, distributions of co-activation values computed from the maximally co-activated module at each hierarchical level, using 1000 surrogates that preserve spatial autocorrelation; blue, the empirical co-activation values for unshuffled brain activity; statistical significance is indicated whenever empirical values exceed high percentiles of the null distribution. (**c**-**f** -**i**) Similar analysis to the previous panels, comparing empirical and shuffled activity, but repeated using 100 null parcellations inferred from SCCM matrices. White pixels indicate where a statistically significant spatial overlap is achieved for any hierarchical level in null parcellations. Note the very low false positive rates in **c**-**f**, demonstrating a very significant overlap between empirical SC modules and visual/motor activity configurations. However, a high false positive rate is observed in null SCCM modules for forebrain activity (which can be seen in one example parcellation in **g**, bottom row). This preserved overlap in shuffled networks can be explained by the spatial distance constraints imposed on edge swapping, combined with the protruding geometry of the forebrain. Most forebrain connections are preserved after shuffling, thus preserving the forebrain module in the null network models (see Supp. Fig. S8 **d**-**ef**, top left matrix corners, where the block-like structure of forebrain connectivity is partially preserved in SCCM but not in CM).

**Figure S15:**
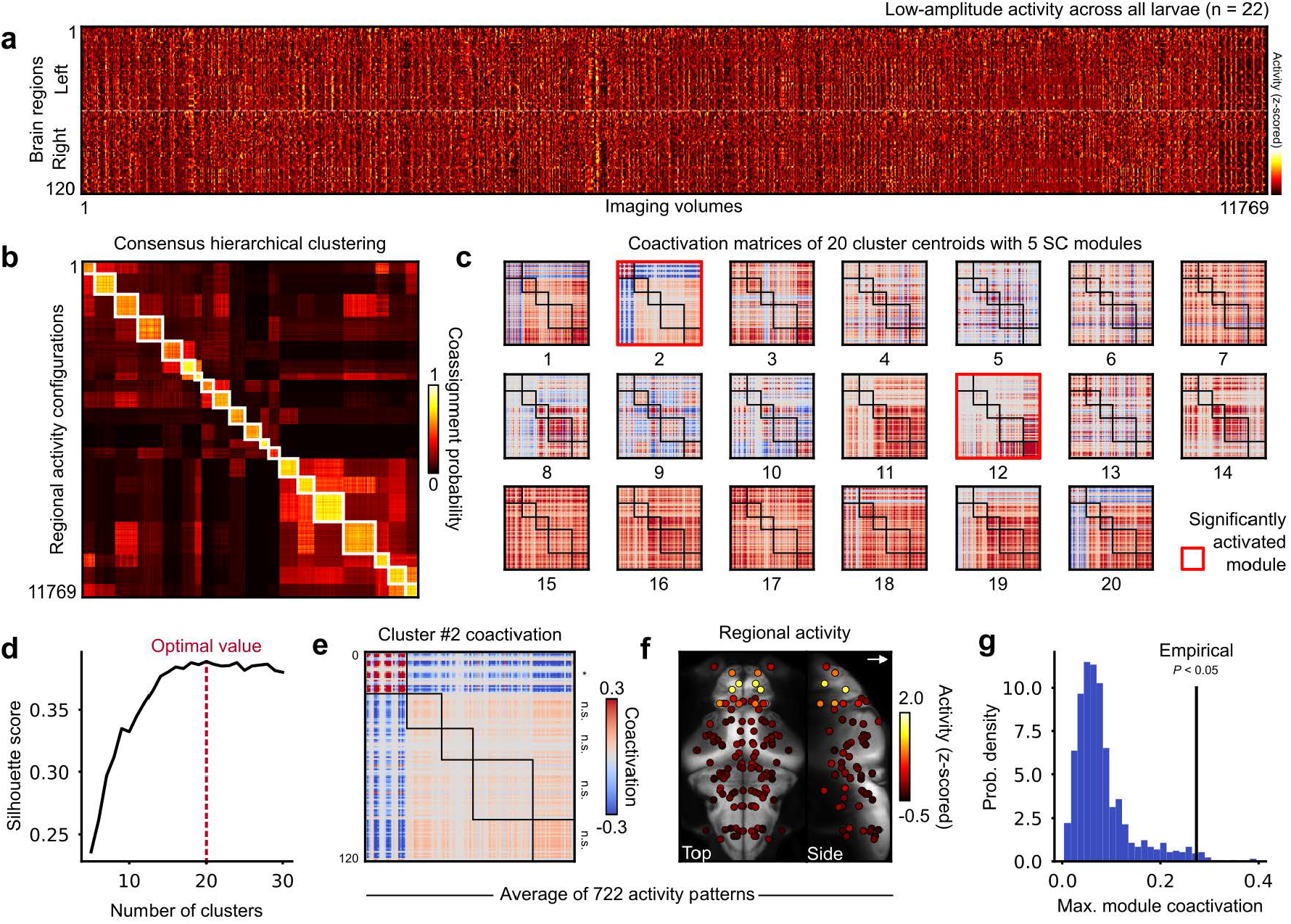
Supplemental clustering analysis of low-amplitude brain activity configurations, which recovers modular forebrain activity. (**a**) Concatenated regional activity matrix of 22 larvae, wherein high-amplitude co-fluctuation frames have been excluded. (**b**) Hierarchical clustering of a consensus matrix obtained through 500 K-means clustering runs of the previous matrix, with the number of clusters *k* ranging from X to Y. (**c**) Co-fluctuation matrices of 20 cluster centroids, each scaled arbitrarily for visualization. Clusters in which a significantly co-activated module is observed are highlighted in red; cluster 12 recovers lower-amplitude motor activity, while cluster 2 consists of forebrain activity. (**d**) Silhouette score computed on different subdivisions of the consensus matrix, which peaks at 20 clusters. (**e**) Close-up on the co-activation matrix of cluster 2, with statistical significance of the maximally co-activated module indicated on the right. (**f**) Average of cluster 2, which displays high forebrain activity. (**g**) Null distribution of 1000 activity patterns shuffled while approximately preserving autocorrelation, compared with the empirical forebrain co-activation value (*P <* 0.05).

**Figure S16:**
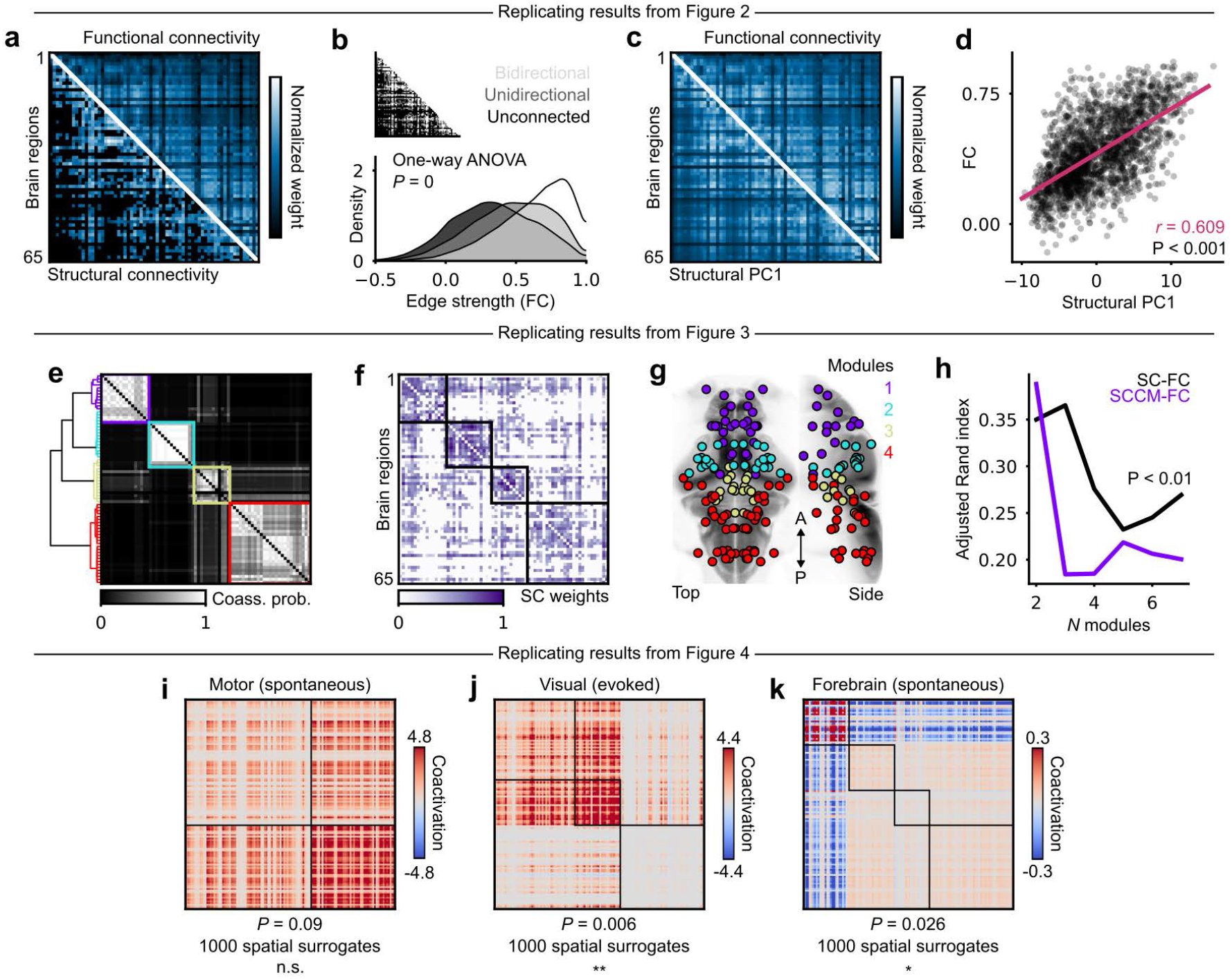
Replication of key results with a sparser SC matrix obtained using wiring method 3 in Supp. Fig. S7. All results remain significant, except the modularity of motor co-activation events due to the loss of modular subdivisions in the hindbrain. The first row (**a**-**d**) replicates panels **c, d, k, l** from Fig. 2, respectively. The second row (**e**-**h**) replicates panels **a, b, c, i** from Fig. 3, respectively. Note that SC-FC community overlap is only benchmarked against SCCM communities, as all other null models perform significantly worse. Third row (**i**-**k**) replicates panels **g, h, m** in Fig. 4, respectively. Peak modularity is observed at different hierarchical levels in this new partition of nodes.

**Figure S17:**
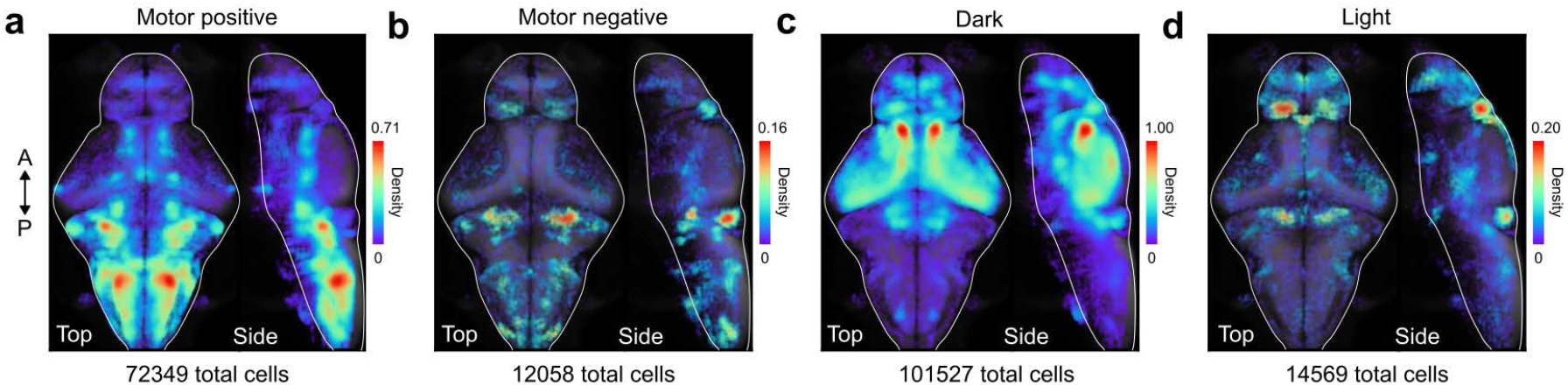
Four neuronal population densities displayed in Fig. 5, without applying a spatial *P* -value test. (**a**) Motor-correlated cells. (**b**) Motor-anticorrelated cells. (**c**) Dark-responsive cells. (**d**) Light-responsive cells. Sparse and spuriously correlated neurons are retained here, but many low-density clusters are also better defined, such as the inferior olive in dark-responsive cells. Some small brain nuclei can have more variability in the number of intersected cells per animal due to the z-spacing of the imaging planes and the angle of the brain, which can lead to slightly lower spatial reproducibility in cell identifications within these regions. The total number of cells indicated below each panel is the number of significantly correlated neurons compiled across all larvae to form a single spatial density.

**Figure S18:**
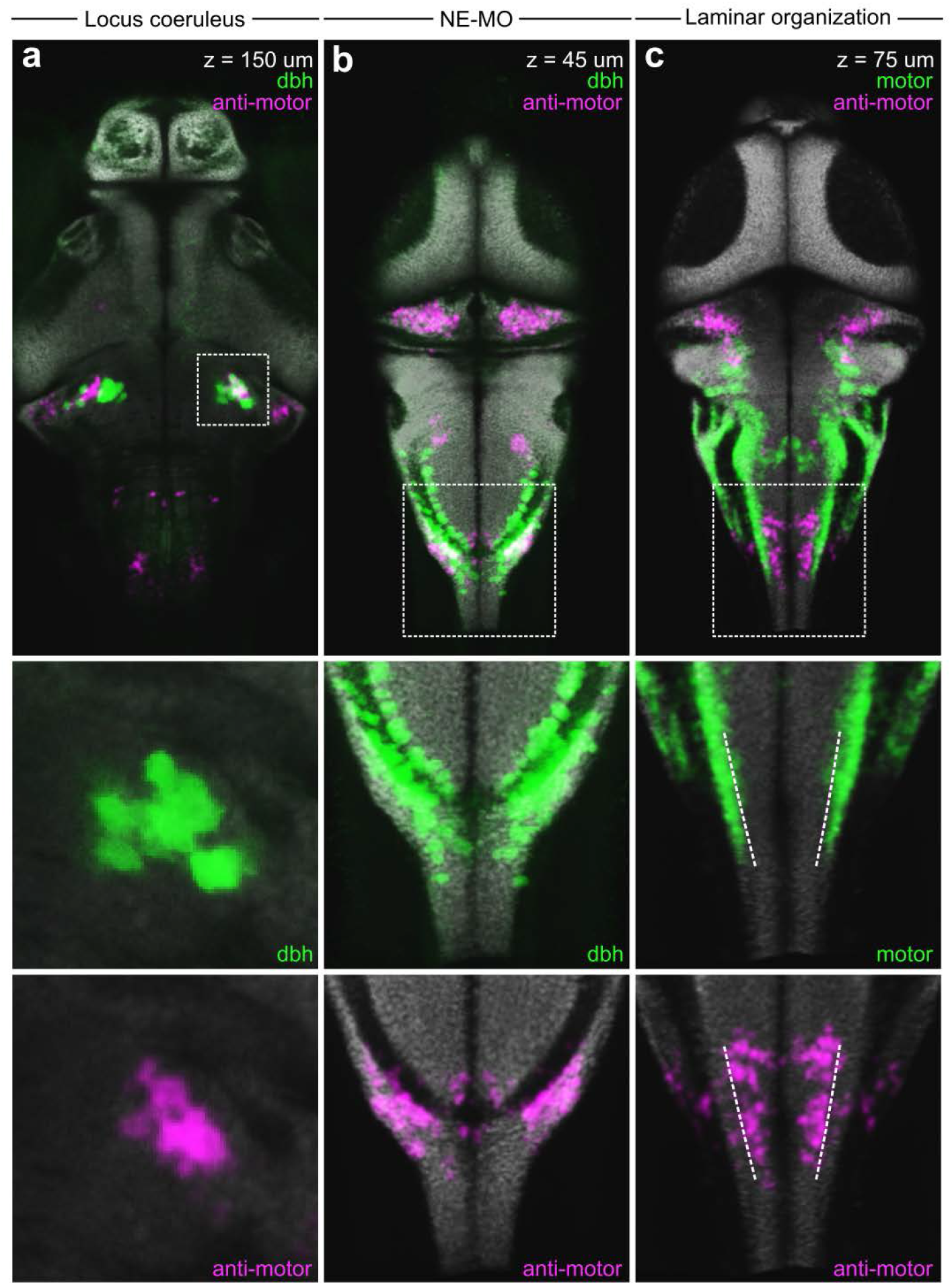
Anatomical validation of motor-anticorrelated cells. (**a**) Overlap between the density of motor negative neurons (anti-motor, magenta) and a dopamine beta-hydroxylase HCR marker (dbh, green) from mapZebrain ^33^. Good overlap is observed within the locus coeruleus (LC, white box), suggesting the involvement of LC noradrenergic cells in suppressing spontaneous behavior. (**b**) Overlap between motor negative neurons (magenta) and dbh-expressing neurons (green) at a more superficial depth, highlighting a strong correspondence between anti-motor cells and noradrenergic neurons of the medulla oblongata (NE-MO), whose initiating role in suppressing futile swimming has been characterized previously ^76,77^. (**c**) Laminar organization of positive (green) and negative (magenta) motor correlations in caudal hindbrain; dashed white lines indicate an abrupt boundary between both cell populations, suggesting a well-defined and organized neural substrate which opposes the hypothesis that these correlations could reflect movement artifacts.

**Figure S19:**
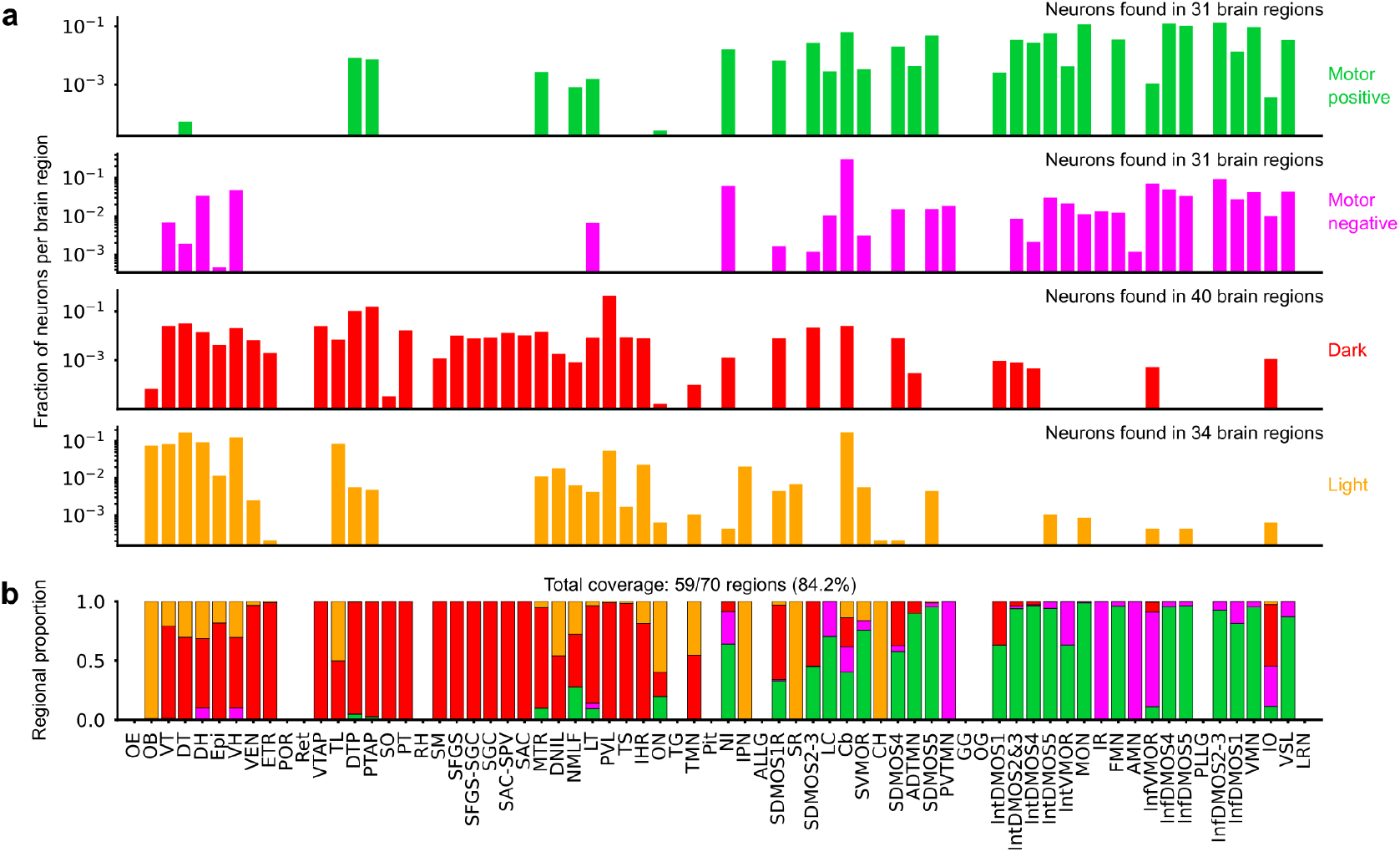
Distribution of the four main functionally defined cell populations in 70 mapZebrain atlas regions, after applying a spatial *P*-value threshold. (**a**) Average fraction of cells per brain region for each functional category (indicated on the right). (**b**) Regional proportions of the four cell types. Note the high diversity in certain regions, such as cerebellum (Cb) or inferior olive (IO), which is also reflected in Fig. 5 **h**. Collectively, the four populations can be found in 84.2% of brain regions. Regions are ordered from anterior to posterior.

**Figure S20:**
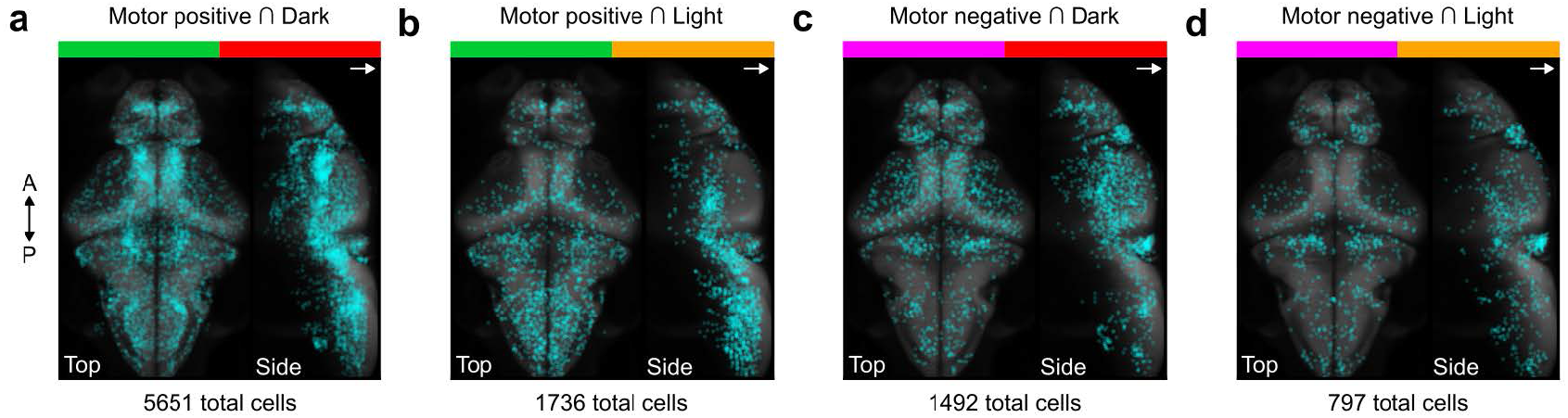
Anatomical distributions of polyfunctional cells obtained through intersections of the four main functionally defined cell populations, without spatial *P*-value thresholding. (**a**) Cells that are simultaneously correlated with motor activity and dark stimuli. (**b**) Cells that are simultaneously correlated with motor activity and light stimuli. (**c**) Cells that are simultaneously anti-correlated with motor activity and positively correlated with dark stimuli.(**d**) Cells that are simultaneously anti-correlated with motor activity and positively correlated with light stimuli. The total number of cells indicated below each panel is compiled across *n* = 18 larvae.

**Figure S21:**
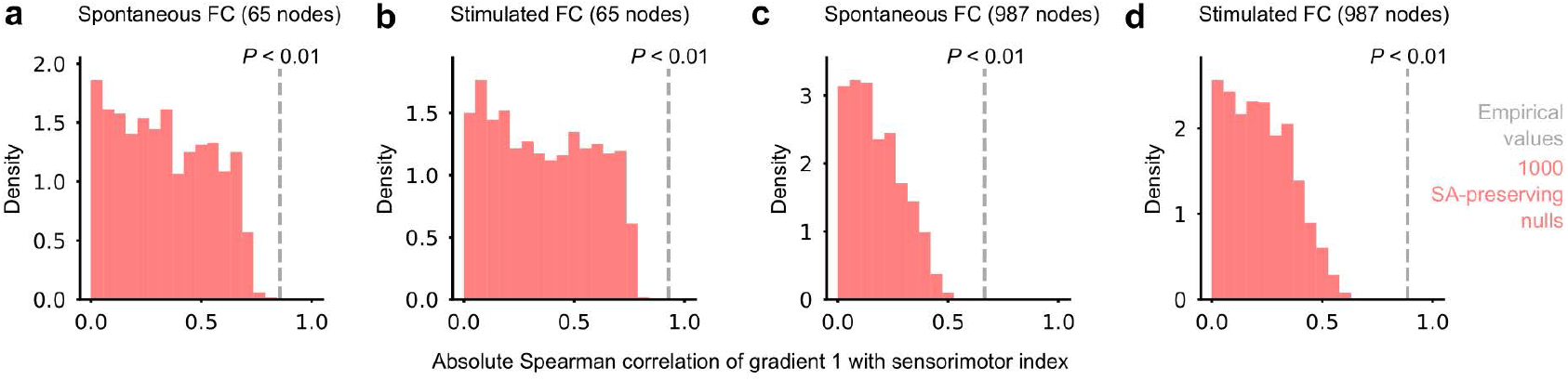
Statistical validation of the correlation between the first functional gradient and the sensorimotor index. (**a**-**d**) Distributions of 1000 spatially randomized sensorimotor indices (pink), compared to empirical correlations (gray dashed lines). All empirical correlations are significant (*P <* 0.01).

**Figure S22:**
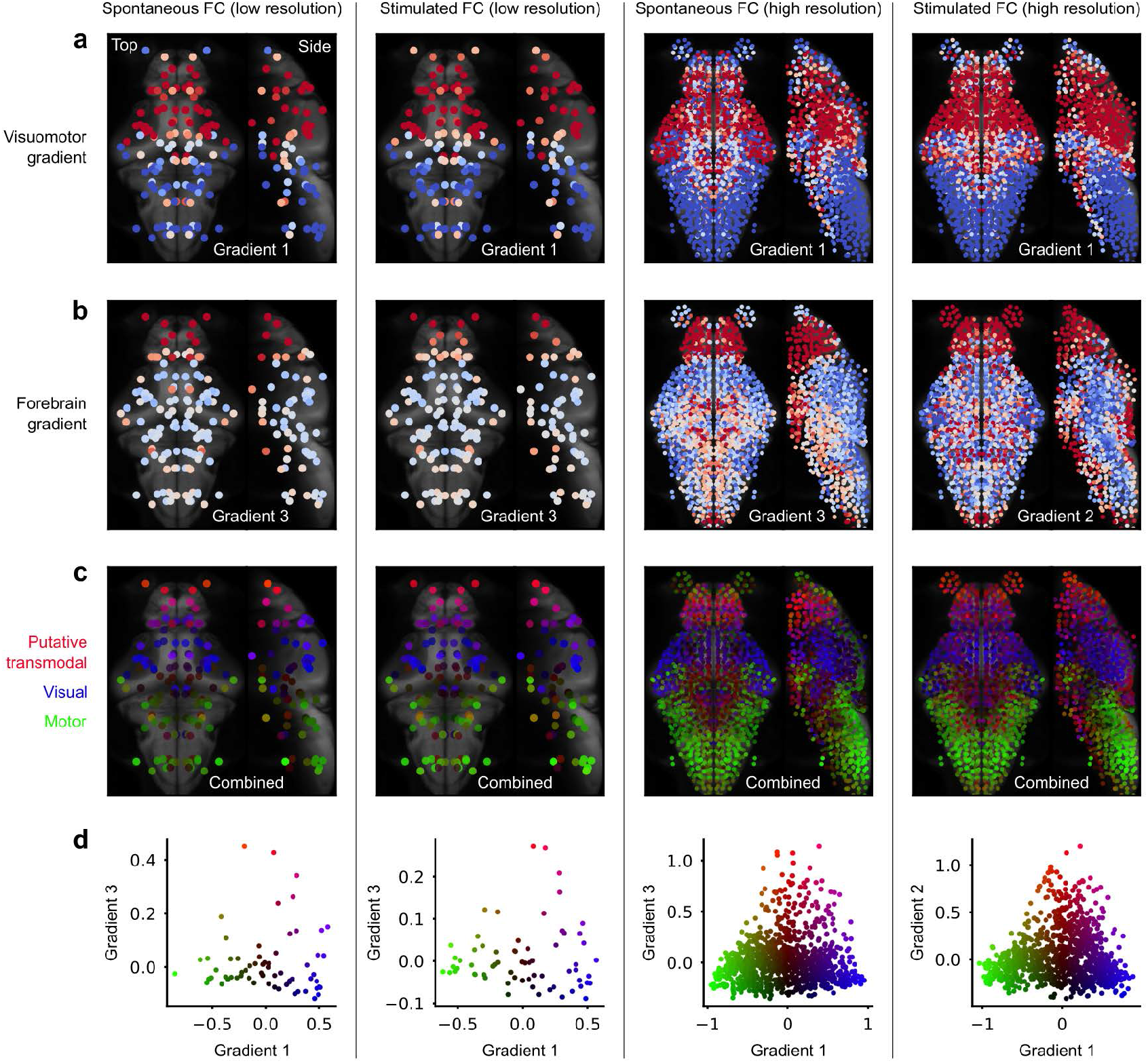
A second functional gradient begins in visuomotor regions and peaks in forebrain regions. (**a**) Main visuomotor gradient extracted from 4 different FC matrices (from left to right, similar to Fig. 6**f** -**i**). (**b**) Forebrain gradient of each FC matrix. (**c**) Combination of visuomotor and forebrain gradients with red, blue, and green colors associated with different extremities of the embedding in **d**, and reflecting putative transmodal, visual, and motor functions, respectively. (**d**) Bidimensional embedding of network nodes based on their values along the visuomotor and forebrain gradients. Note the clear emergence of a triangular embedding structure in higher resolution parcellations.

**Figure S23:**
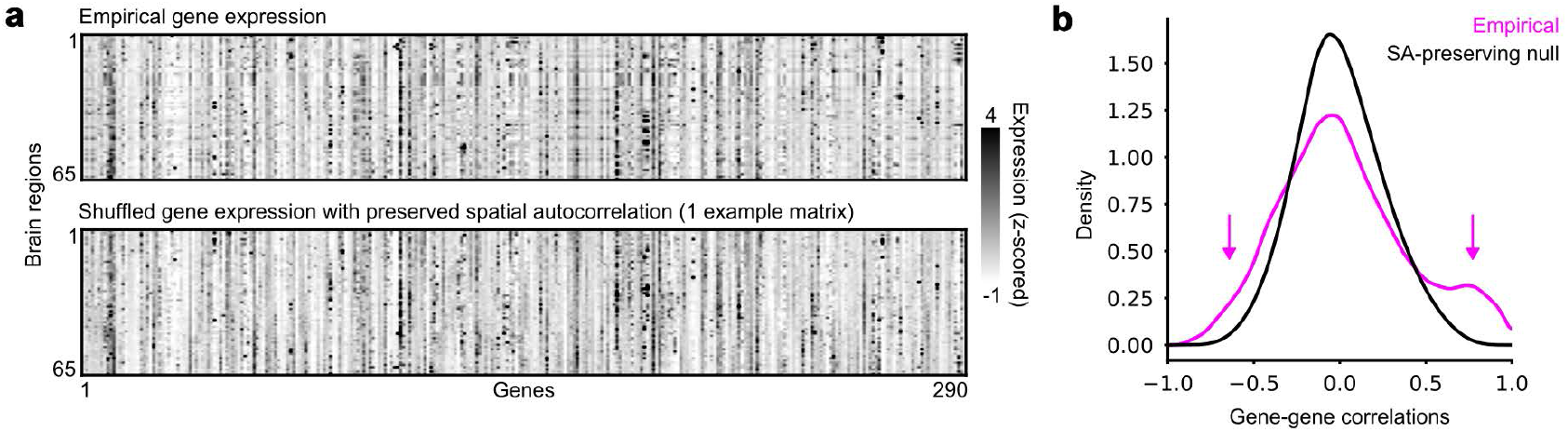
Reduced correlations in shuffled gene datasets. (**a**) Top, empirical gene expression matrix displayed in Fig. 7**b**; bottom, one example spatially shuffled gene expression matrix, where each gene is shuffled independently while approximately preserving its spatial autocorrelation. (**b**) Gene-gene correlation distributions obtained by correlating the spatial expression profiles of each gene to every other gene; the black distribution is generated by compiling correlation values across 1000 null gene expression matrices. Arrows indicate a negative heavy tail and a positive bump in real gene correlations, suggesting a higher level of spatial redundancy in gene expression patterns. This high redundancy explains the diminishing returns when considering a high number of genes in the simulated annealing optimization of gene co-expression to reproduce FC. This eventually leads to a better reconstruction of FC by null gene sets when the number of selected genes roughly exceeds 20 (see Fig. 7**d**).

